# Quantitative Method for Assessing the Role of Lysine & Arginine Post-Translational Modifications in Nonalcoholic Steatohepatitis

**DOI:** 10.1101/2020.01.17.910943

**Authors:** Aaron E. Robinson, Aleksandra Binek, Vidya Venkatraman, Brian C. Searle, Ronald J. Holewinski, George Rosenberger, Sarah J. Parker, Nathan Basisty, Xueshu Xie, Peder J. Lund, Gautam Saxena, José M. Mato, Benjamin A. Garcia, Birgit Schilling, Shelly C. Lu, Jennifer E. Van Eyk

**Author notes:** Corresponding author: Jennifer E. Van Eyk, 127 S. San Vicente Blvd, Advanced Health Sciences Pavilion, 9302, Los Angeles, CA 90048.

## Abstract

Proteoforms containing post-translational modifications (PTMs) represent a degree of functional diversity only harnessed through analytically precise simultaneous quantification of multiple PTMs. Here we present a method to accurately differentiate an unmodified peptide from its PTM-containing counterpart through data-independent acquisition-mass spectrometry, leveraging small precursor mass windows to physically separate modified peptidoforms from each other during MS2 acquisition. We utilize a lysine and arginine PTM-enriched peptide assay library and site localization algorithm to simultaneously localize and quantify seven PTMs including mono-, di-, and tri-methylation, acetylation, and succinylation in addition to total protein quantification in a single MS run without the need to enrich experimental samples. To evaluate biological relevance, this method was applied to liver lysate from differentially methylated non-alcoholic steatohepatitis (NASH) mouse models. We report altered methylation and acetylation together with total protein changes drive the novel hypothesis of a regulatory function of PTMs in protein synthesis and mRNA stability in NASH.

**Graphical Abstract:** 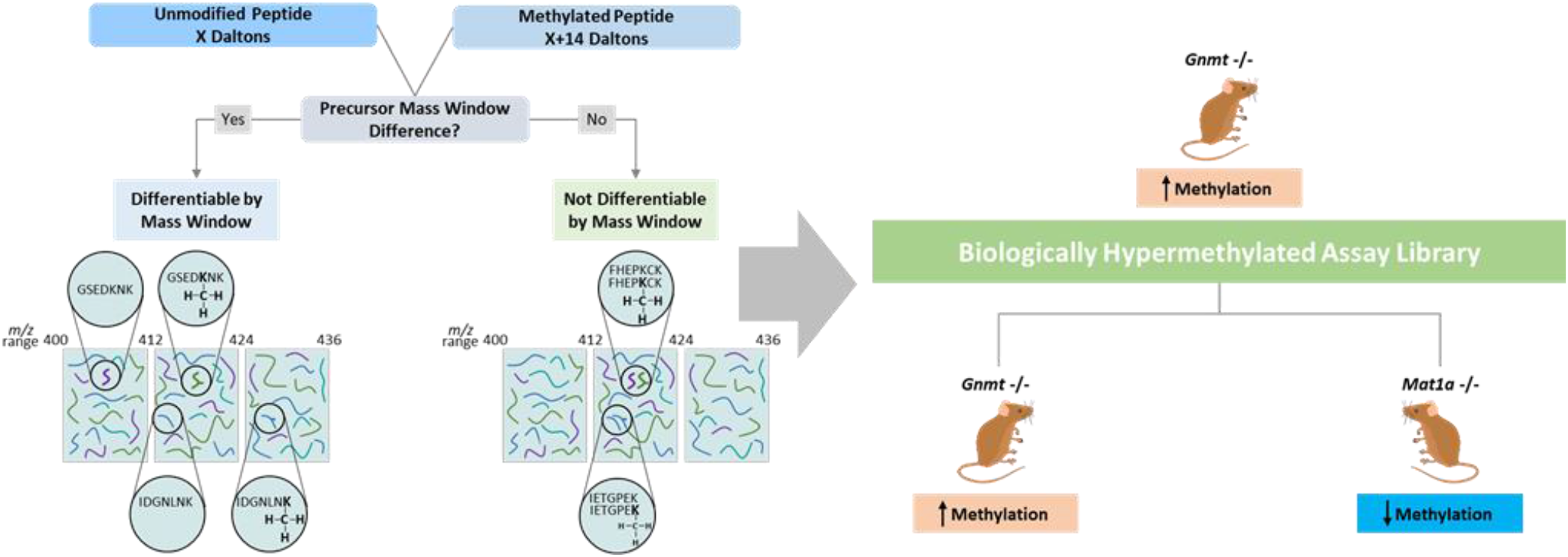

## Introduction

While there have been substantial advances in biochemical methods for the detection and subsequent mass spectrometry (MS) to quantify phosphorylation (e.g., metal affinity and immunoaffinity enrichments), detection of other protein translational modifications (PTMs), like methylation, acetylation and succinylation, which are also dynamic regulators of cellular signaling,^1–3^ has lagged behind.^4,5^ Technological advancements in MS instrument sampling speed have expanded the dynamic range of precursor sampling for fragmentation by data dependent acquisition-MS (DDA-MS), and concomitantly enabled reductions in precursor window widths for data independent acquisition-MS (DIA-MS) based fragmentation while still maintaining reasonable cycle times. More importantly, DIA-MS allows different peptidoforms to be physically segregated into different windows providing increased specificity. DIA-MS coupled with tissue-specific PTM-enriched assay libraries improves the consistency of peptide quantitation including low abundant peptides with PTMs.^6,7^ Recent published algorithms PIQUed,^8^ Peptide Collapse plugin for Perseus,^9^ Inference of Peptidoforms (IPF),^10^ and Thesaurus,^11^ claim reduced false site localization and greater PTM recovery. Taken together this provides an alternative to the classical approach to elucidate PTM dynamics in complex biological systems and a more complete understanding of the complexity of cellular signaling.^4,5,10,12–14^

In proteins, methylation generally occurs on Arg and Lys residues with the ε-amino group of Lys being either mono, di, or tri-methylated by protein lysine methyltransferases while Arg can be mono or di-methylated by protein arginine methyltransferases. In mammalian cells, S-Adenosylmethionine (SAMe) is the exclusive methyl donor for both Lys and Arg methylation.^3^ The majority of research in protein methylation has focused on histone methylation and its role in epigenetics. Histone methylation of Lys and Arg is an integral part of the histone code known to alter the physical properties of chromatin subsequently altering gene expression.^3^ Conversely, methylation of non-histone proteins has only recently emerged as a biologically relevant PTM and recent research has shown its importance in cellular signaling and function.^15–19^ This information has in part been obtained from global protein methylation proteomic analysis by the development of labeling and enrichment techniques like hM SILAC, high and low pH SCX fractionation, and peptide immunoprecipitations of methylated Lys and Arg and subsequent mass spectrometry.^20–23^ However, antibody based methyl-enrichments have a number of challenges and limitations: (1) they require a large amount of sample and multiple antibodies to cover all methyl forms; (2) they often contain a sequence bias for antigen recognition, thus selectively assaying a portion of the methylome while masking an unknown but likely significant portion of methyl-sites; and (3) by nature these approaches do not enrich the unmodified form of the peptide, hindering quantification of all peptidoforms together and thus impairing the ability to study the relative abundance of PTMs.^24,25^

To overcome these challenges and to facilitate a comprehensive and technically straightforward quantitative survey of the methylome in biological samples, we leveraged small precursor mass window DIA-MS to physically isolate modified peptidoforms from each other. Each experimental sample is then assayed and compared against a carefully curated peptide library, consisting of both unmodified and methylated peptides. In addition to methylation, lysine residues can be modified by various other PTM’s, including acetylation and succinylation.^26–30^ To understand the extent of lysine PTMs, we expanded our DIA assay library to include succinylated and acetylated peptides, thus providing an opportunity to quantify methylated, acetylated, succinylated, and unmodified peptides from the same sample and acquisition run. This not only allows for classical cellular protein quantification, but also provides the opportunity to quantify peptidoforms containing Lys methylation, succinylation, acetylation, and Arg methylation simultaneously from a single DIA acquisition. Traditional DDA and DIA based PTM-immunoprecipitation enrichment approaches rely on antibody enrichment of modified peptides and by nature are unable to quantify unmodified peptides in the same acquisition or enrich for a single protein which has limited utility.^4,29^ In our method, we quantify both modified and unmodified peptides using DIA acquisitions with small precursor mass windows requiring only five percent starting material than traditional PTM immunoprecipitations. The important distinction of small precursor mass windows is we can physically trap each modified peptidoform away from other modified peptidoforms and the unmodified peptide. Combined with an algorithm which provides localization scores and false localization rate (FLR) calculations for each modified amino-acid residue, our approach provides both protein quantification and quantitative data simultaneously on seven PTMs.

To highlight biological insight that can be gained using our small precursor mass window DIA method, two mouse models of nonalcoholic steatohepatitis (NASH) were employed, each chosen due to their extreme methylation potential. Glycine N-Methyltransferase knockout (*Gnmt* -/-) mice have increased SAMe, have higher methylation potential, and are biologically hypermethylated while methionine adenosyltransferase A1 knockout (*Mat1a* -/-) mice are SAMe deficient, have reduced methylation potential, and are therefore hypomethylated.^17 31–33^ Both animal models spontaneously develop NASH and resemble major subtypes of human disease.^31,34^ Interestingly, disease in both animal models can be treated by normalization of SAMe levels highlighting the importance of SAMe, methylation, and Lys PTMs in NASH.^31,35,36^

## Materials and Methods

### Methylated Peptide Synthesis

100 synthesized unmodified and mono-, di-, tri-methylated and acetylated peptides (JPT Peptide Technologies GmbH, Berlin, Germany) were supplied as lyophilized powder. Peptides were resuspended in 0.1% Formic Acid and pooled at 1:1 ratio, aliquoted, and stored at –80°C until used. Stabile Isotopically labeled (SIL) methyl-peptides were synthesized and were supplied as a lyophilized powder. The supplier provided product characterization (MALDI-TOF and HPLC traces) as proof of MW and purity accuracy. The peptides were of >95% purity (New England Peptide). Peptides were solubilized in 0.1% Formic Acid and pooled at 1:1 ratio, aliquoted, and stored at –80°C until used.

### Liver Tissue Sample Preparation

Livers were obtained from ten-month old methionine *Mat1a* -/- male mice in a C57Bl/6 background with hepatic lipid accumulation and their aged-matched wild-type (WT) male sibling littermates,^31^ and three-month old *Gnmt* -/- mice in a C57Bl/6 background with hepatic lipid accumulation with age matched WT littermates.^32,33^ Animals were bred and housed in the CIC bioGUNE animal unit, accredited by the Association for Assessment and Accreditation of Laboratory Animal Care International (AAALAC). Animals were fed with standard commercial chow animal diet (Ref. 2914, Envigo, Barcelona, Spain).

Frozen mouse livers (n=6/condition) were ground to powder under liquid N2 in a cryohomogenizer (Retsch). Tissue powder was thawed and lysed in 8M urea and 100mM TRIS-HCL, pH 8.0 and samples were ultrasonicated (QSonica) at 4 °C for 10 min in 10 second repeating on/off intervals of 10 seconds and centrifuged at 16,000 x g for 10min at 4 °C. The protein concentration of soluble supernatant was determined via Bicinchoninic Acid Assay (Thermo). 100μg of protein was reduced with DTT (15mM) for 1h at 37 °C, alkylated with iodoacetamide (30mM) for 30min at room temperature in the dark, diluted to a final concentration of 2M Urea with 100mM TRIS-HCL, pH 8.0 and digested for 16 hours on a shaker at 37 °C with a 1:40 ratio of Trypsin/Lys-C mix (Promega). Each sample was de-salted using HLB plates (Oasis HLB 30μm, 5mg sorbent, Waters).

### Orbitrap Fusion Lumos MS Acquisition

For DIA-MS analysis, a Orbitrap LUMOS Fusion mass spectrometer (Thermo Scientific) was equipped with an EasySpray ion source and connected to Ultimate 3000 nano LC system (Thermo Scientific). Peptides were loaded onto a PepMap RSLC C18 column (2 μm, 100 Å, 150 μm i.d. x 15 cm, Thermo) using a flow rate of 1.4 μL/min for 7 min at 1% B (mobile phase A was 0.1% formic acid in water and mobile phase B was 0.1 % formic acid in acetonitrile) after which point they were separated with a linear gradient of 5-20%B for 45 minutes, 20-35%B for 15 min, 35-85%B for 3 min, holding at 85%B for 5 minutes and re-equilibrating at 1%B for 5 minutes. For longer gradient method (120min) peptides were separated with a linear gradient of 5-20%B for 90 minutes, 20-35% for 30 min, 35-85%B for 3 min, holding at 85%B for 5 minutes and re-equilibrating at 1%B for 5 minutes. Each sample was followed by a blank injection to both clean the column and re-equilibrate at 1%B. The nano-source capillary temperature was set to 300 °C and the spray voltage was set to 1.8 kV. iRT Standards (Biognosys) were added to each sample before acquisition.

MS1 scans were acquired in the Orbitrap at a resolution of 60,000 FWHM from mass range 400-1000m/z. For MS1 scans the AGC target was set to 3×10^5^ ions with a max fill time of 50 ms. DIA MS2 scans were acquired in the Orbitrap at a resolution of 15000 FWHM with fragmentation in the HCD cell at a normalized CE of 30. The MS2 AGC was set to 5e4 target ions and a max fill time of 22ms. DIA was performed using 4 Da (150 scan events, 6.46 second cycle time with 1% standard deviation), 6 Da (100 scan events), or 12 Da (50 scan events) windows over the precursor mass range of 400-1000 m/z and the MS2 mass range was set from 100-1500 m/z. Multiplexed precursor mass window DIA acquisitions were acquired using the approach outlined in *Sidoli et al*.,^37^ using 4 Da and 6 Da precursor mass windows outlined above.

For DDA-MS methods used for the Orbitrap Fusion Lumos, see **Supplementary Methods**.

### 6600 TripleTOF MS Acquisition

For DIA-MS analysis, the 6600 TripleTOF (Sciex) was connected to Eksigent 415 LC system that was operated in micro-flow mode. The mobile phase A was comprised of 0.1% aqueous formic acid and mobile phase B was 0.1% formic acid in acetonitrile. Peptides were pre-loaded onto the trap column (ChromXP C18CL 10 x 0.3 mm 5 μm 120Å) at a flow rate of 10 μL/min for 3 min and separated on the analytical column (ChromXP C18CL 150 x 0.3mm 3 μm 120Å) at a flow rate of 5 μL/min using a linear A-B gradient composed of 3-35% A for 60 min, 35-85% B for 2 min, then and isocratic hold at 85% for 5 min with re-equilibrating at 3% A for 7 min. Temperature was set to 30°C for the analytical column. Source parameters were set to the following values: Gas 1 = 15, Gas 2 = 20, Curtain Gas = 25, Source temp = 100, and Voltage = 5500V. iRT Standards (Biognosys) were added to each sample before acquisition.

DIA MS1 scans were acquired using a dwell time of 250 ms in the mass range of 400-1250 m/z at 45,000 FWHM. DIA MS2 scans were acquired in high-sensitivity mode at 15,000 FWHM over the precursor range of 400-1250 m/z with the MS2 range of 100-1800 m/z using 100 variable windows with a dwell time of 30ms. Additionally, the same samples were acquired in DIA using 4 Da fixed windows over a precursor range of 400-1000 m/z with a dwell time of 22ms.

For DDA-MS methods used for the Sciex 6600 TripleTOF, see **Supplementary Methods**.

### Generation of Spectral Libraries and Database Searches

Sample preparation and MS acquisition settings for spectral library are located in **Supplementary Methods**.

PTM enriched DIA assay library was generated as described by *Parker et al.* 2016^6^ with some modifications. Briefly, all DDA files were converted to mzXML and searched through the Trans Proteomic Pipeline (TPP) using 3 algorithms, (1) Comet;^38^ (2) X!tandem! Native scoring;^39^ and (3) X!tandem! K-scoring^40^ against a reviewed, mouse canonical protein sequence database, downloaded from the Uniprot database on January 24^th^, 2019, containing 17,002 target proteins and 17,002 randomized decoy proteins. Precursor and fragment mass tolerance for all of these search algorithms were set to 10ppm. Peptide probability modeling was performed using the TPP peptide prophet “xinteract” and the results searches were combined using the TPP “interprophet parser”. Further filtering was done using Mayu to select peptide spectral match probability values consistent with a 1% peptide false discovery rate (FDR) and a spectral library was generated using the TPP SpectraST tool. Retention times were then normalized to ‘indexed’ retention time space^41^ using the custom python script spectrast2spectrast_irt.py, publicly available via the MSPROTEOMICSTOOLS python package (https://github.com/msproteomicstools/msproteomicstools). Biognosys internal retention time reference peptides were added to each sample immediately before acquisition and were used for retention time (RT) alignment. As needed, the expanded Common internal RT standards reference peptides^42^ was used to align RT in fractionated library samples, removing any outliers as described in *Parker et al. 2016*.^6,41^ Retention time normalized splib files were consolidated to a single spectrum entry for each modified peptide sequence using the “consensus spectrum” function in Spectrast. These spectral libraries were then filtered to generate peptide assay libraries by selecting the top twenty most intense b or y ion fragments.

The resulting file was imported into Skyline for DIA analysis as described in *Egertson et al.*^43^ and was simultaneously converted to the OpenSWATH library input TraML format with decoys appended. The Skyline and OpenSWATH assay libraries normalized to ‘indexed’ retention time space and all raw files used in this study will be available at Panorama (https://panoramaweb.org/methylation_methods_1.url, Proteome Exchange ID: PXD012621) as a public resource to the proteomics community.

### Quantitation of Individual Specimen by DIA-MS

Peak group extraction and FDR analysis was done as outlined in *Parker et al, 2016.*^6^ Briefly, raw intensity data for peptide fragments was extracted from DIA files using the open source OpenSWATH workflow^44^ against the sample specific peptide assay. Then, retention time prediction was made using the Biognosys iRT Standards spiked into each sample.

Target and decoy peptides were then extracted. scored and analyzed using the mProphet algorithm^45^ to determine scoring cut-offs consistent with 1% FDR. Peak group extraction data from each DIA file was combined using the ‘feature alignment’ script, which performs data alignment and modeling analysis across an experimental dataset.^46^ Finally, all duplicate peptides were removed from the dataset to ensure that peptide sequences are proteotypic to a given protein in our FASTA database.

### Development of Site Localization Scoring System for DIA-MS

We performed five individual data searches with Thesaurus search engine (version 0.9.4)^11^ presetting a different modification (acetylation, succinylation, mono-methylation, di-methylation or tri-methylation) in each analysis. For the Thesaurus searches, we used a filtered (modified peptidoforms only) version of the PTM enriched DIA Assay Library generated in this study. Localization strategy parameter was set to recalibrated (peak width only). Library precursor and fragment mass tolerance were both set to 10 ppm. Peptidoforms were selected for false localization rate (FLR or localization FDR) analysis if they passed user-specified localization p-value (p>0.05) and localization score (>2) thresholds. All the detected peptidoforms in the individually processed DIA-MS RAW files (at 5% global peptide FDR) were posteriorly integrated and processed with Percolator 3.01 (with the global peptide FDR filter set at 1%). Once all five modification-specific analyses were generated, five output results files (.elib) were integrated into one output and processed with Percolator (at 1% global peptide FDR filter). Localization p-values for passing peptidoforms are independently corrected using Benjamini-Hochberg FDR test (localization FDR). Peptidoforms meeting FLR <1% criteria were used for further biological analyses.

### Normalization and Quantitation

The total ion current (TIC) associated with the MS2 signal across the chromatogram was calculated for normalization using in-house software. This ‘MS2 Signal’ of each file was used to adjust the transition intensity of each peptide in a corresponding file. To obtain a measure of total protein quantity, all modified (methylated, acetylated, and succinylated) peptides were removed and the remaining peptides analyzed separately. Normalized transition-level data of the unmodified peptides were subsequently processed using mapDIA to obtain protein level quantitation.^47^ Normalized transition-level data of the all modified (methylated, acetylated, and succinylated) peptides were then analyzed individually through mapDIA to obtain peptide level quantitation of PTM containing peptides. Finally, to obtain a PTM/Total ratio, the intensity of the PTM containing peptide was divided by the intensity of its respective protein, calculated using mapDIA from an aggregate of all unmodified peptides for that protein.

### Data Visualization and Validation

The aligned OpenSWATH output was imported into Skyline^43^ which was used to visualize and manually validate methylated peptides. The Skyline documents containing DIA acquisitions extracted against their respective peptide assay libraries are available at Panorama (https://panoramaweb.org/methylation_methods_1.url, Proteome Exchange ID: PXD012621).

### Metabolomics Analysis

Liver metabolic profiles were semi quantified using an ultra-high performance liquid chromatography (UHPLC)-Time of Flight-MS based platform as described previously in *Alonso et al. 2017*.^31^ This platform provided coverage over polar metabolites, such as vitamins, nucleosides, nucleotides, carboxylic acids, coenzyme-A derivatives, carbohydrate precursors/derivatives, and redoxelectron-carriers was used. Briefly, proteins were precipitated from the liver tissue (15 mg) by adding methanol spiked with metabolites not detected in unspiked cell extracts (internal standards) and samples were homogenized using a Precellys 24 homogenizer (Bertin Technologies, Montigny-le-Bretonneux, France) at 6500 rpm for 23 seconds x 1 round, and centrifuged at 18,000 x g for 10 minutes at 4 °C. Then, 500 μl were collected, mixed with chloroform and vortexed. After 10 minutes of agitation, samples were centrifuged at 18,000 x g for 15 minutes at 4 °C. Supernatants were collected, dried under vacuum, reconstituted in water and resuspended with agitation for 15 minutes. After centrifugation at 18,000 x g for 5 minutes at 4 °C, samples were transferred to plates for UHPLC-MS analysis as described in *Barbier-Torres et al. 2015*.^48^ Data were pre-processed using the TargetLynx application manager for MassLynx 4.1 software (Waters Corp., Milford, USA). Metabolites were identified prior to the analysis. Peak detection, noise reduction and data normalization were performed as previously described in *Martinez-Arranz et al. 2015*.^49^

## Results

### Synthesized Methylated Peptides

The overall experimental design for quantifying protein methylation is shown in Figure 1. Although we initially focus on Lys methylation to develop the workflow, this approach is applicable to Arg methylation or any other methylated residue. To establish DIA-MS parameters to allow for differentiating a methylated peptide from an unmodified peptide, first a DDA-MS library was built on a Sciex 6600 Triple TOF based on the DDA acquisition of a pool of 400 synthesized peptides (400 femtomoles/peptide) containing either K[Unmodified], K[Monomethyl], K[Dimethyl], or K[Trimethyl]. DIA acquisitions of 400 femtomoles/peptide of the methyl peptide pool were then acquired using a 4 Dalton precursor mass windows and 100 variable mass windows (Figure 1A). A DDA library was additionally created from the methyl peptide pool acquired on the Orbitrap Fusion Lumos (200 femtomoles/peptide). DIA acquisitions of 200 femtomoles/peptide were then acquired with 4, 6, and 12 Dalton fixed precursor mass windows respectively (Figure1B). Generally, the unmodified peptide can be physically separated from its methylated form due to precursor mass difference and thus its fragmentation in a different precursor mass window allowing for differentiation of the unmodified and methyl peptide by precursor mass window alone. However, this is not always the case and depending on precursor mass window size and the charge on the peptide the methylated peptide will fragment in the same precursor mass window as its unmodified form. In this case, a site-specific transition or a co-eluting precursor trace is necessary to correctly identify the post-translational modification (Figure 1C, Table S1). A difference in retention time cannot reliably be used to differentiate an unmodified peptide and its methyl peptidoform as 58.5% of monomethyl peptides elute within 30 seconds of each other in a 60 minute gradient while 92.7% elute within 60 seconds making separation through LC alone unpractical for DIA-MS workflows (Figure S1). Due to this, we next looked into methods to physically separate peptidoforms using precursor mass window widths.

**Figure 1.**
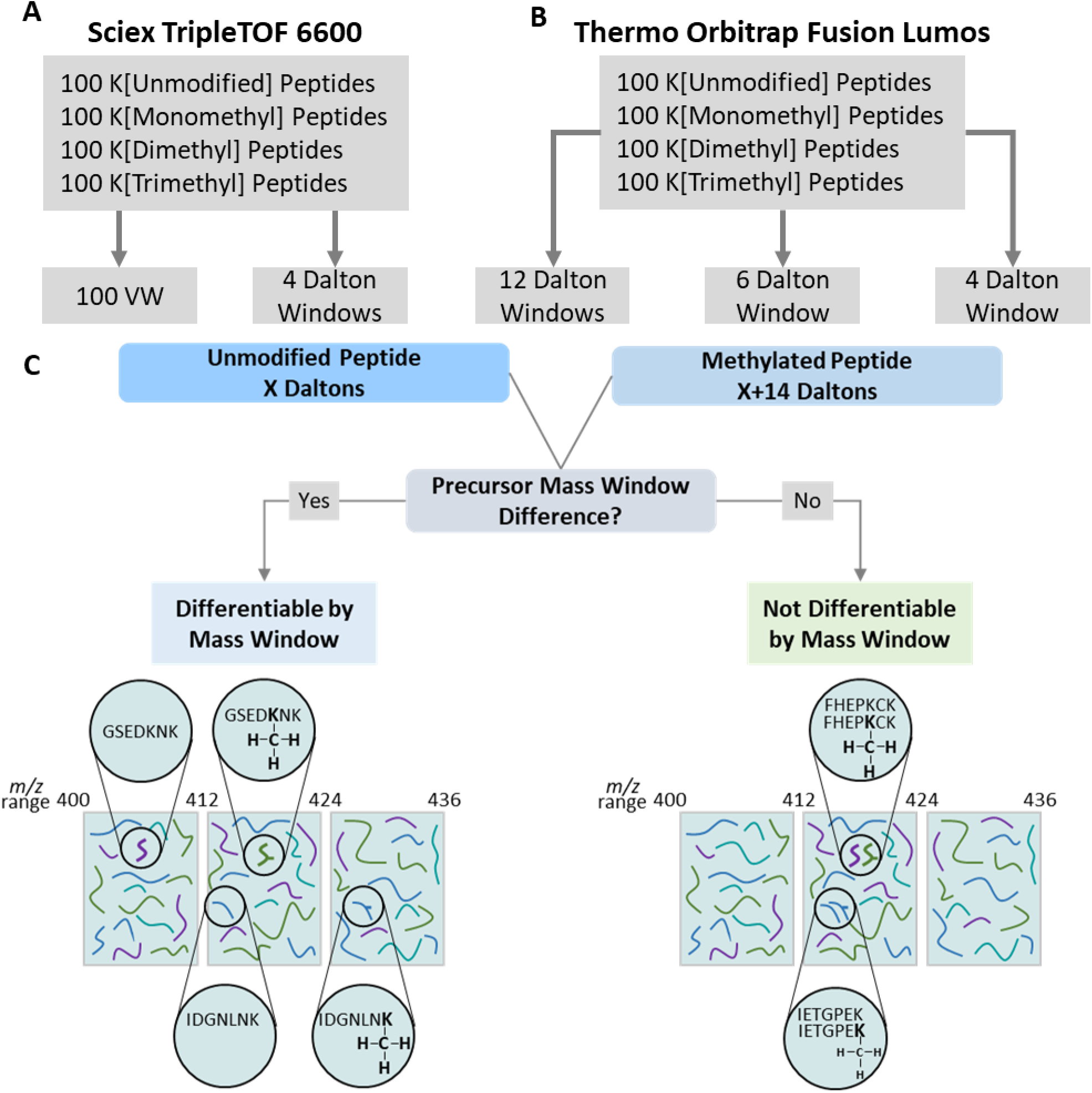
Schematic Overview of Experimental Design: (A) An equimolar pool of 400 synthesized peptides containing K[Unmodified]. K[Monomethyl]. K[Dimethyl], and K[Trimethyl] residues were acquired in DIA on the Sciex TripleTOF 6600 with 4 Dalton and 100 Variable Window (VW) precursor mass windows. (B) Same peptide pool was acquired in DIA on the Thermo Orbitrap Fusion Lumos with 4, 6, and 12 Dalton precursor mass windows. (C) Strategy to differentiate an unmodified peptide from it’s methylated form based off of precursor mass window difference. An unmodified precursor will be fragmented in a different mass window than its methylated form making it differentiable by precursor mass window.

First, we tested whether narrowing the MS1-precursor isolation window width affected the separation of different methylated peptidoforms from their unmethylated counterparts. For example, when the methyl peptide pool was acquired on the Orbitrap Lumos Fusion, we can see that with 12 Dalton precursor mass windows three species of the same peptide have co-fragmented in the same precursor mass window. With 6 Dalton precursor mass windows there are two species of peptide which have co-fragmented in the same window. With 4 Dalton windows, the precursor mass window is small enough that there is only one species of the peptide found (Figure 2 A, B). With the smallest acquisition mass window schema (4 Da) 100% of methylated peptides were separated from their corresponding unmodified forms. Increasing the mass window just 2 Da reduced the segregation slightly, and increasing mass window to 12 Da resulted in only ~ 50% segregation of methylated peptides from unmethylated. (Figure 2C). The 4 Dalton precursor mass windows separate 100% of both +2 and the smaller m/z +3 charged precursors from their methylated forms, whereas the commonly used 100 variable precursor mass window acquisition schema has multiple cases where 2 or more species of peptide co-fragment in the same window (Figure S2, Table S2).

**Figure 2.**
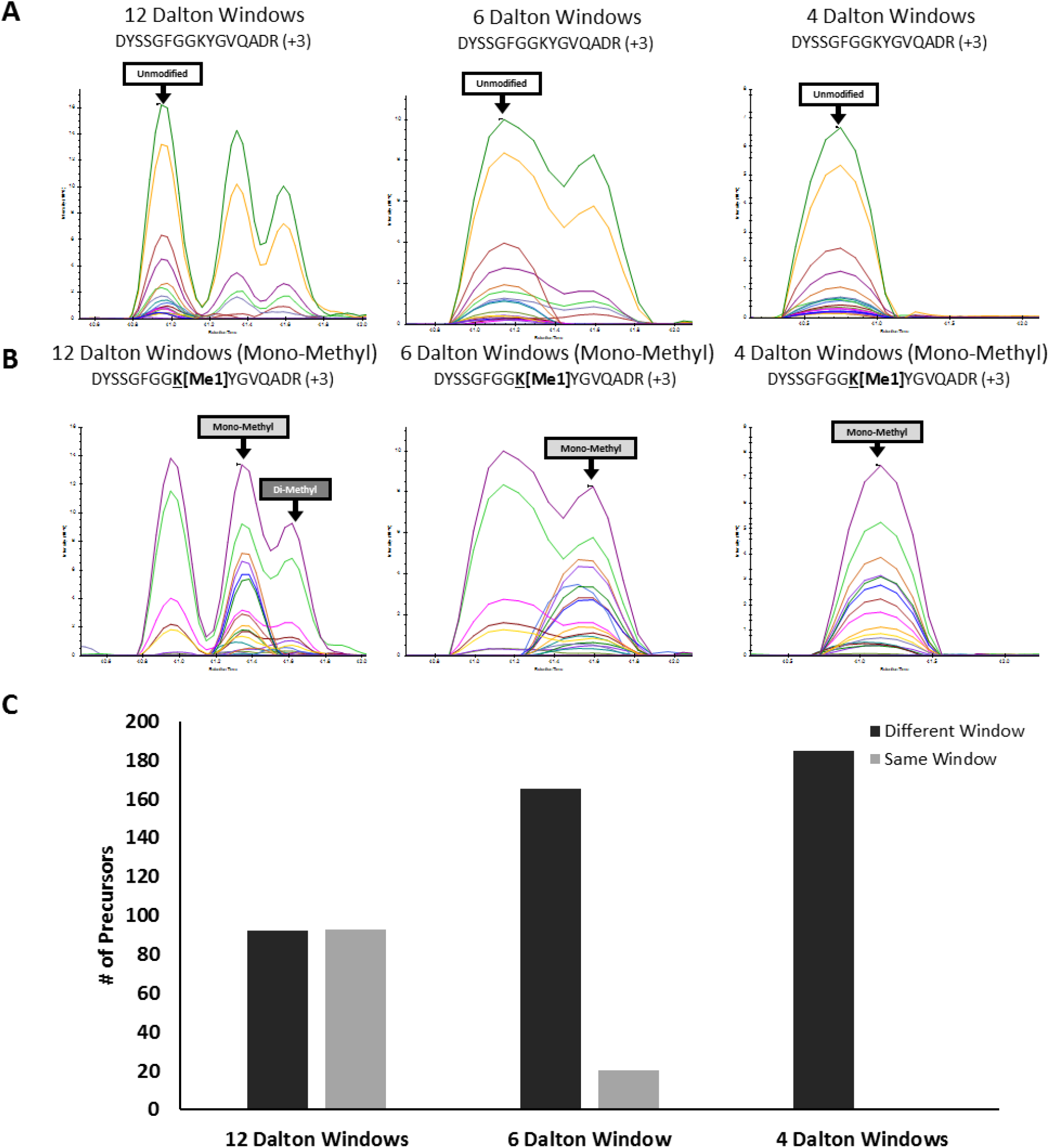
Synthesized Methylated Peptides: Presence of synthesized methylated and unmodified peptides acquired on the Thermo Orbitrap Fusion Lumos with 4, 6, and 12 Dalton precursor mass windows were visualized in Skyline (A) Quantified transitions of DYSSGFGGKYGVQADR (B) Quantified transitions of DYSSGFGG**<u>K</u>**[**Me1**]YGVQADR (C) Graphical representation of unmodified and monomethyl +2 and +3 precursors which can be differentiated from their unmodified forms using 4, 6, and 12 Dalton precursor mass windows. Black represents peptides eluting in different precursor mass windows while grey represents peptides co-eluting in the same precursor mass window.

### Accuracy of Peptide Quantitation in 4 Dalton DIA Method

To establish the accuracy the peptide quantitation obtained from our 4 Da precursor mass window DIA workflow, we performed a dilution series of 71 SIL peptides in complex liver lysate using both 60- and 120-minute gradients. Observation of enough MS2 spectra across the chromatographic elution profile, also termed ‘points across the peak’ is critical for estimating the area-under-the-curve (AUC) of intensity and elution time, and thus, the accuracy of the quantitative data for a given peptide. This is especially true for low abundant peptides which may contain PTMs. The more windows sampled, the more time spent in a given duty cycle, potentially sacrificing quantitative accuracy. The 4 Da DIA method used in this study has a median cycle time of 6.46 seconds with 1.0% standard deviation. We determined that the 4 Da precursor mass windows can detect an average of 6.94 points across a peak in a 60 minute LC gradient while expanding the LC gradient to 120 minutes, and thus widening chromatographic peaks, detected an average of 10.24 points across the peak (Figure 3 A,B). We next estimated the accuracy of quantification by examining linearity of SIL peptide AUC across the dilution series. We next determined the accuracy of quantitation using both 60- and 120-minute gradients by quantifying the SIL peptides in our dilution series. We have linearity in quantification (r^2^ > 0.95) of 87/91 (95.6%) of transitions corresponding to precursors for the SIL peptides in the 60-minute gradient and linearity in quantification (r^2^ > 0.95) for 91/91 (100%) of the transitions corresponding to precursors for the SIL peptides in the 120-minute gradient.

**Figure 3.**
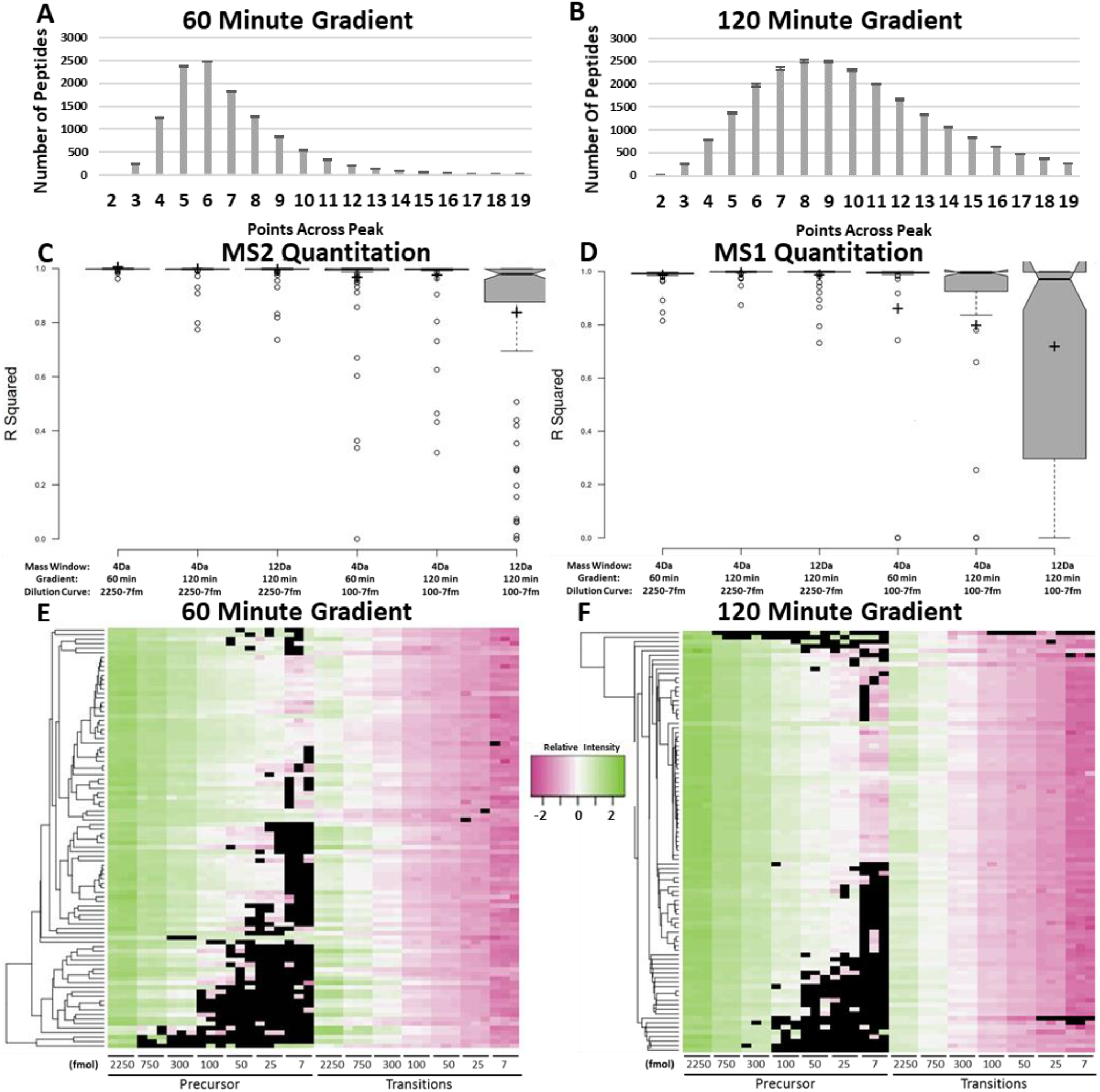
SIL Peptide Dilution Series: 71 SIL peptides were spiked into complex liver lysate in varying concentrations and acquired on the Thermo Orbitrap Fusion Lumos in DIA with 4da precursor mass windows (A) Distribution of points across the chromatographic peak of all endogenous peptides with a 60 minute LC gradient and (B) a 120 minute LC gradient. (C) Box plots showing the accuracy of quantitation using of a dilution series of SIL peptides in complex liver lysate acquired with different DIA acquisition methods and gradients for MS2 quantitation and (D) MS1 quantitation. (E) Heatmap corresponding to intensity of SIL peptides in dilution series with a 60 minute gradient. (F) Heatmap corresponding to intensity of SIL peptides in dilution series with a 120 minute gradient. For A/B: All data are mean ± s.e.m. of three technical replicates. For CZD: For all data, center lines show the medians; box limits indicate the 25th and 75th percentiles as determined by R software; whiskers extend 1.5 times the interquartile range from the 25th and 75th percentiles, outliers are represented by dots; crosses represent sample means, n = 91 SIL precursors. For E/F: Color represents intensity of peptide, black represents missing data. Data is three replicates per concentration of SIL peptides.

We additionally looked at the lower end of our dilution curve (7 fmol to 100 fmol on column) to see how our method performs on lower abundant peptides, which are a better representation of the biological concentration of PTMs in complex lysate. We found a (r^2^ > 0.95) for 84/91 (92.3%) and 85/91 (93.4%) of the transitions corresponding to SIL precursors for the 60-minute gradient and the 120-minute gradient respectively. Furthermore, we performed a dilution series of the same 71 SIL peptides in complex liver lysate with 12 Da precursor mass windows in a 120-minute gradient and found linearity in quantification (r^2^ > 0.95) of 87/91 (95.6%) of transitions corresponding to precursors for the SIL peptides. Unexpectedly, looking at the lower end of our dilution curve for 12 Da precursor mass window acquisitions, we found only 59/91 (64.8%) of the transitions corresponding to SIL precursors show linearity in quantification showing that our 4 Dalton precursor mass window DIA acquisition method provides more accurate quantitation of lower abundant peptides than a 12 Dalton precursor mass window method (Figure 3C, Table S3).

### Total Protein Normalization Allows for More Accurate Quantification of Methylation in NASH

After extracting 4 Dalton precursor mass window DIA acquisition maps of *Gnmt* -/- and *Mat1a* -/- mouse liver cellular lysate against the hypermethylated DIA assay library described in **Supplementary Results**, we quantified 78 methylated peptides in OpenSWATH with unmodified peptides quantifiable for the same protein. To determine the relative abundance of PTMs, having total protein quantification is required. Since we are using a DIA assay library containing both methylated and unmodified peptides extracted against an unenriched complex lysate, we have the ability to quantify both methylated Arg and Lys containing peptides and unmodified peptides in a single DIA acquisition. For example, a unique mono-methylated peptide corresponding to EF1A1 K165 was found to be significantly increased in the hypermethylated *Gnmt* -/- NASH model compared to WT (p<0.0005), whereas the same site was significantly decreased in the hypomethylated *Mat1a* -/- NASH model compared to WT (p<0.05) (Figure 4A). After normalizing to total protein, by taking the ratio of the intensity of the methylated peptide compared to the intensity of an aggregate of all unmodified EF1A1 peptides as determined by MapDIA, *Mat1a* -/- hypomethylation compared to WT was more significant (p<0.0005) (Figure 4B). By being able to normalize to the aggregate protein intensity derived from unmodified peptides in the same MS acquisition we have the ability to look at Methyl/Total ratio which allows for easy normalization of the methylated peptide to the aggregate protein abundance in one DIA acquisition, providing similar data as a protein immunoblot for a PTM and total protein, which can further the biological insight that one can take away from this type of data. We tested our approach versus library free approaches and found significantly better quantitation of low abundant modified peptides with the described methodology (Supplementary Results, Figure 3-4, Table S3-4)

**Figure 4.**
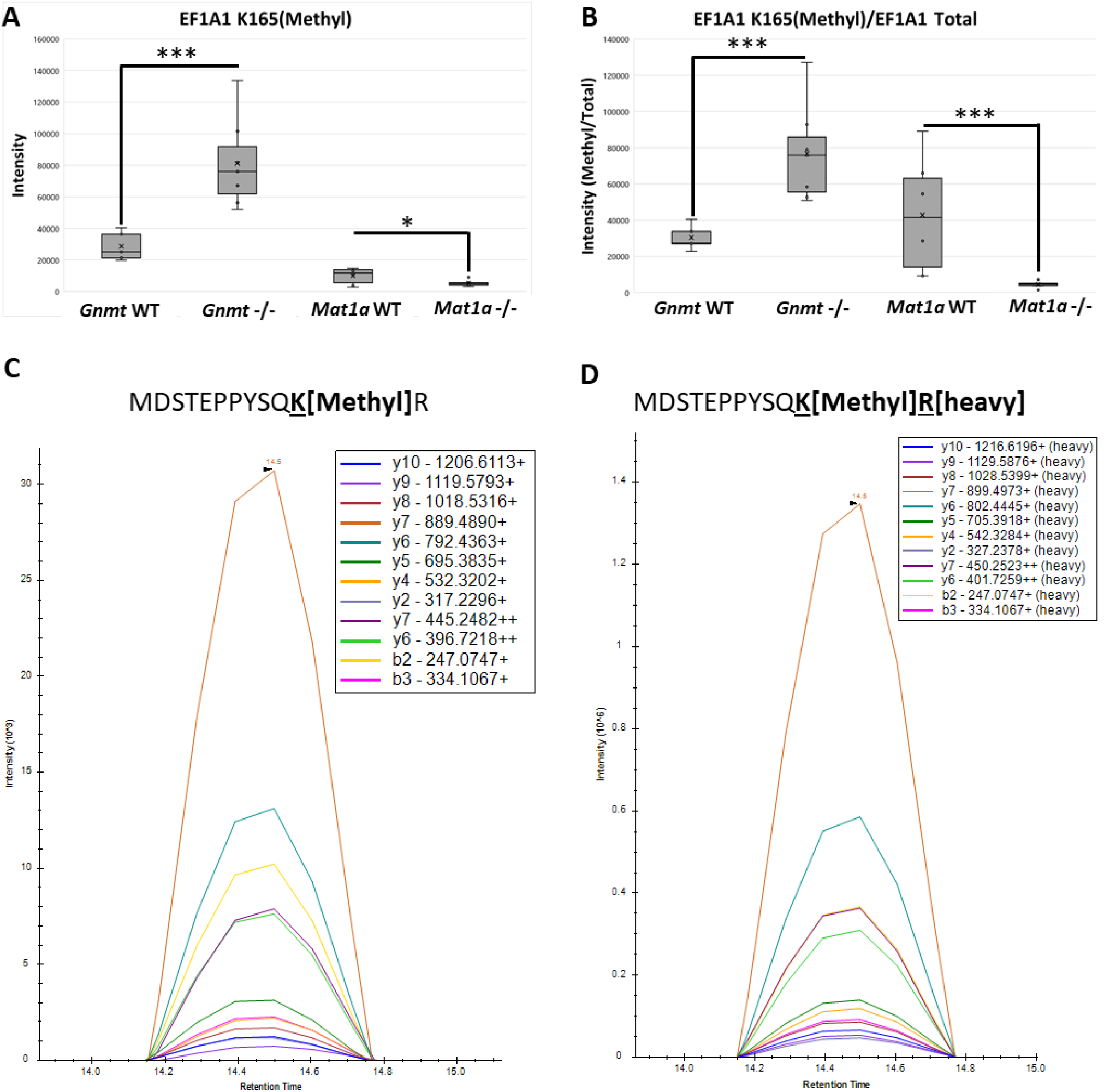
*In-vivo* Methylation: (A) Quantification of the MS2 TIC normalized intensity of peptide containing EF1A1 K[I65] Mono-Methyl in *Gnmt -/-* and wild-type littermates and *Mat1a* -/- and wild-type littermates Data are box and whisker plots of six biological replicates. Two-tailed Student’s t-test. \*\*\**P* < 0.005. *\*P* < 0.05 (B) Quantification of the intensity of the MS2 TIC normalized peptide containing EF1A1 K[I65] MonoMethyl divided by the intensity of EF1A1 total protein, determined by EF1A1 unmodified peptides using MAPDIA in *Gnmt -/-* and wild-type littermates and *Mat1a -/-* and wild-type littermates Data are box and whisker plots of six biological replicates per condition. Two-tailed Student’s t-test. \*\*\**P* < 0.0005 (C) Skyline Visualization of the XIC of die peptide MDSTEPPYSQK[Metliyl]R which corresponds to EF1A1 K[I65] Mono-Methyl in *Gnmt -/-* complex liver lysate (D) Skyline Visualization of the XIC of a stably isotopically labeled peptide MDSTEPPYSQK[Metliyl]R[lieavy] which corresponds to EF1A1 K[I65] Mono-Methyl spiked into *Gnmt -/-* complex liver lysate

To further validate our methyl library, we synthesized heavy methyl-peptides to match three proteins localized in different cellular compartments which were present in complex liver lysate from *Gnmt* -/- mice with NASH. Next, these heavy peptides were added to the *Gnmt* -/- complex liver lysate and 4 Dalton precursor mass window DIA-MS was carried out. We found co-eluting peaks from all of the endogenous methyl-peptides and their corresponding synthetic heavy peptides (Figure 4D, Figure S3). From our SCX fractionated peptide assay library in this study, 53.7% of unmodified peptide precursors are +2 and 38.5% of precursors are +3. However, looking at only methylated precursors, only 22.2% are +2 while 58.1% of the methylated precursors are +3 (Figure S4, Table S5). This is likely due to more missed cleavages due to methylation inhibiting trypsin’s ability creating longer peptides with a higher charge.^50^ We can differentiate 80.3% of methylated precursors in our library from their unmodified forms due to precursor charge state of +2 and +3. When looking further into the 19.7% of peptides that remain with a charge of +4 or higher and calculating the respective precursor mass of the methylated peptide and its corresponding unmodified peptide, we can differentiate 99.1% of the remaining methylated peptides from their unmodified form by which mass window each precursor falls in leaving 0.24% (2/808) unresolvable by 4 Dalton precursor mass windows (Table S5). To this point, our approach solves the common problem of false positive or ambiguous identification of methyl peptides (Supplementary Results, Figure S5-6)

### Acetylation and Succinylation are Differentially Changed in NASH Subtypes

Next, looking at Lys modifications acetylation and succinylation, and we quantified 176 acetylated and 59 succinylated peptides with OpenSWATH in the DIA-MS runs on the mouse tissue which had unmodified peptides quantifiable of the same protein. As with methylation, to determine relative abundance of the modified peptidoform, having total protein quantification is required. By using a DIA assay library containing both modified and unmodified peptides and assaying against complex lysate, we have the ability to quantify both acetylated and succinylated Lys containing peptides and unmodified peptides in a single DIA acquisition.

A unique succinylated peptide corresponding to mitochondrial malate dehydrogenase (MDHM) K165(Succinyl) was significantly increased in the *Gnmt* -/- NASH model compared to WT (p<0.005) while the *Mat1a* -/- NASH model compared to WT was not significantly changed (Figure 5A). When normalizing to total protein, by taking a ratio of the intensity of the succinylated peptide compared to the intensity of unmodified MDHM peptides as determined by MapDIA, *Mat1a* -/- compared to WT became significantly decreased in the *Mat1a* -/- NASH model compared to WT (p<0.005) while *Gnmt* -/- NASH model compared to WT remained significant (p<0.05) (Figure 5B). Furthermore, a peptide proteotypic for Histone H4 (H4) with three acetyl sites, K8, K12 and K16 respectively was found in the DIA-MS datasets. This peptide was not significantly changed in either the *Gnmt* -/- NASH model compared to WT or the *Mat1a* -/- NASH model compared to WT (Figure 5D). When normalizing to total protein, by taking a ratio of the intensity of the acetylated peptide compared to the intensity of unmodified H4 peptides, *Gnmt* -/- compared to WT became significantly decreased versus WT (p<0.05) and *Mat1a* -/- became significantly increased compared to WT (p<0.005) (Figure 5E). As with methylation, being able to normalize to the protein intensity derived from unmodified peptides in the same MS acquisition we have the ability to look at Succinyl/Total or Acetyl/Total ratio which provides an additional piece of information, similar to a protein immunoblot for total protein, which allows for a representation of the biological concentration of the PTM in complex lysate (Figure 5F).

**Figure 5.**
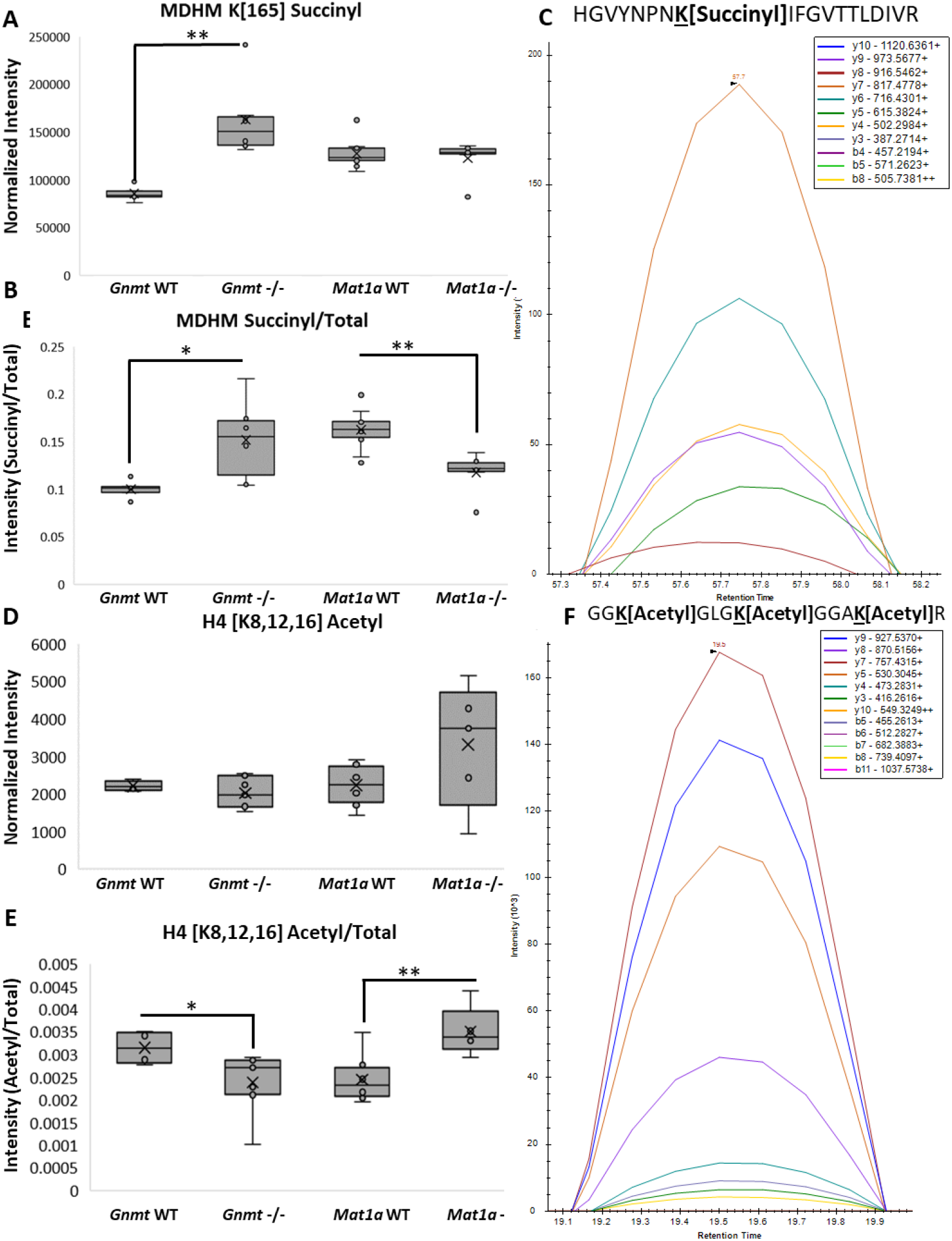
*In-vivo* Succinylation and Acetylation: (A) Quantification of the MS2 TIC normalized intensity of peptide containing MDHM K[I65] Succinyl in *Gnmt* and wild-type littermates and *Mat1a* -/- and wild-type littermates (n=6/condition) Data are box and whisker plots of six biological replicates per condition. Twotailed Student’s t-test, \*\**P* < 0.005 (B) Quantification of the intensity of the MS2 TIC normalized peptide containing MDHM K[I65] Succinyl divided by the intensity of MDHM total protein, determined by MDHM unmodified peptides using MAPDIA. Data are box and whisker plots of six biological replicates per condition. Two-tailed Student’s t-test, \*\**P* < 0.005, \**P* < 0.05 (C) Skyline Visualization of the XIC of the peptide containing MDHM K[I65] Succinyl (D) Quantification of the MS2 TIC normalized intensity of a proteotypic peptide mapping to H4 K[8,I2,16] Acetyl in *Gnmt -/-* and wild-type littermates and *Matla* -/- and wild-type littermates. Data are box and whisker plots of six biological replicates per condition. (E) Quantification of the intensity of the MS2 TIC normalized peptide containing H4 K[8,I2,16] Acetyl divided by the intensity of H4 total protein, determined by H4 unmodified peptides using MAPDIA in *Grant -/-* and wild-type littermates and *Matla* -/- and wild-type littermates. Data are box and whisker plots of six biological replicates per condition. Two-tailed Student’s t-test, ** *P* < 0.005, \**P* < 0.05 (F) Skyline Visualization of the XIC of the peptide containingH4 K[8,I2,16] Acetyl

We confirmed the presence of these succinylated and acetylated peptides by showing the highest abundant transitions found in DDA and DIA acquisitions of ‘one-pot’ acetyl-Lys and succinyl-Lys immunoaffinity enrichment from WT mouse liver were also found in complex *Gnmt -/-* liver lysate (Figure S7, S8). Furthermore, for the tri-acetylated H4 K[8,12,16] peptide, the relative abundance of these transitions was constant between DIA acquisitions acquired on the Orbitrap Fusion Lumos and the Sciex TripleTOF 6600. These acquisitions were run on instruments located at two different research institutes, and yet yielded the same confident identifications of an acetylated and succinylated peptide respectively.

### Modified Peptidoform Localization with 4 Dalton DIA Method

We modified the Thesaurus algorthim^11^ and developed a new version (0.9.4) which can localize all of the PTMs assayed in the described method. We tested the new version of Thesaurus using 400 synthesized peptides containing either K[Monomethyl], K[Dimethyl], K[Trimethyl], or K[Acetyl] on the same a non-terminal Lys residue of 100 peptide backbones. After creating a DDA Library containing the MS2 spectra of synthesized peptides, we performed peptidoform localizations of 4Da precursor mass window DIA acquisitions using Thesaurus 0.9.4. Out of peptidoforms which were detected with less than 1% global peptide FDR, we were able to localize with less than 1% FLR 62/76 (81.6%) of mono-methylated peptides, 60/77 (78.0%) of di-methylated peptides, 52/65 (80%) of tri-methylated peptides, and 61/76 (80.0%) of acetylated peptides (Figure 6A, Table S6). Overall, 42 peptide backbones were correctly localized with all 4 modifications (Figure 6B) across 3 replicates of DIA-MS acquisitions.

**Figure 6.**
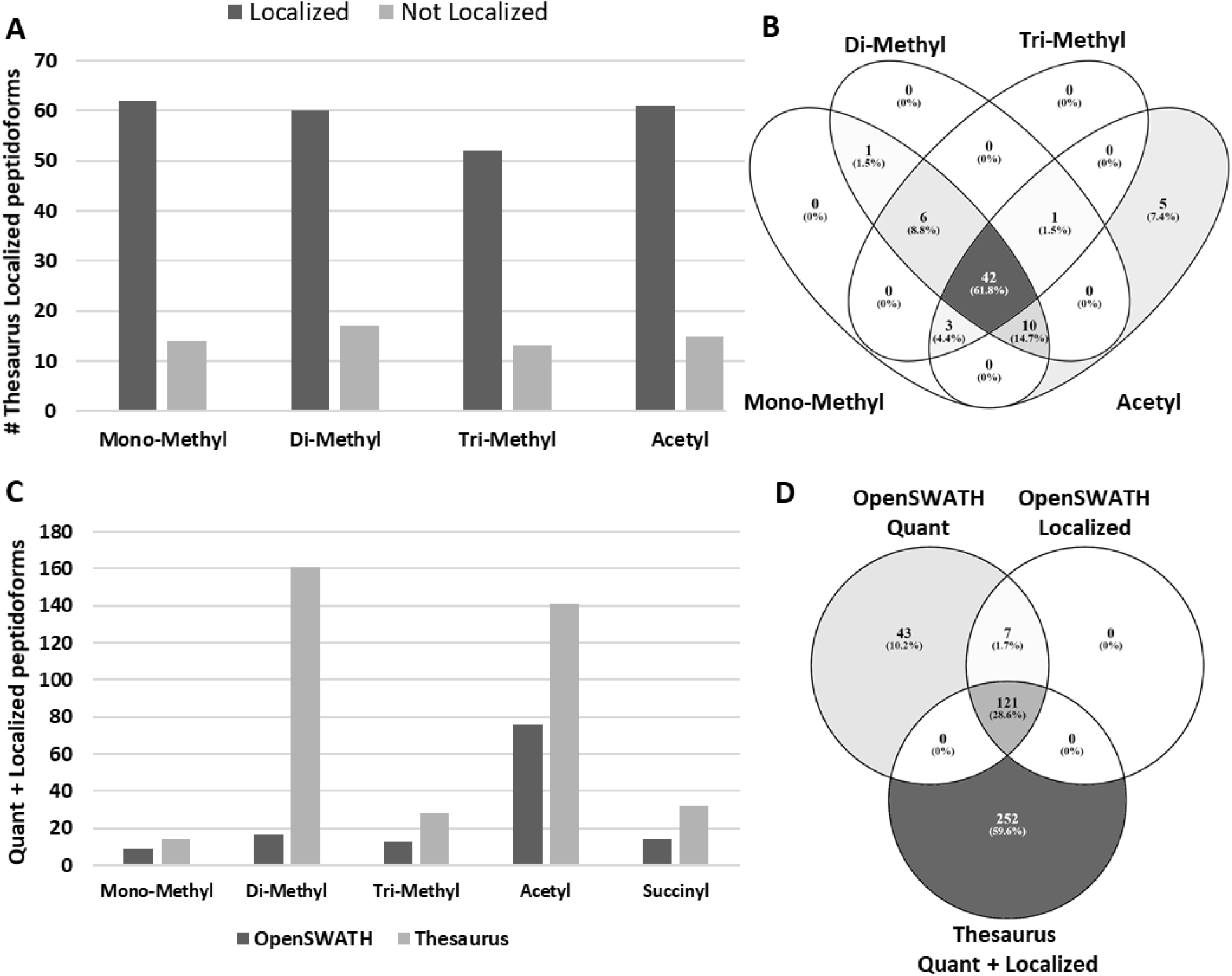
Modified Peptidoform Localization: **(A)** Localization results of modified peptidoforms using Thesaurus 0.9.4 froma mixture of synthesized peptides containing the same peptide backbone but with modifications (mono-, di-, tri-methylated and acetylated) on the same a noil-terminal Lys residue. **(B)** Venn diagram showing the overlap of Thesaurus localization results from each modification type from mixture of synthesized peptides **(C)** Quantified and localized modified peptidoform from 4da DIA acquisitions of complex cellular lysate from Gnmt -/- and Gnmt WT livers assayed against the PTM enriched DIA Assay Library in either OpenSWATH 2.3.0 or Thesaurus 0.9.4 and identified with >1% FDR in ≥4/12 (≥33%) of samples hi each respective approach in addition to >1% FLR localization calculated in Thesaurus 0.9.4 and applied to both the OpenSWATH and Thesaurus results. (D) Venn diagram showing the overlap of quantifiable peptidoforms from OpenSWATH. quantifiable peptidoforms from OpenSWATH which were localized using Thesaurus, andpeptidoforms which were both quantifiable and localization from Thesaurus.

After assessing the performance of Thesaurus 0.9.4 on our ground truth dataset of synthesized peptides, we moved on to PTM localizations in a complex cell lysate. Using our PTM enriched DIA-MS Assay Library and performing analysis in Thesaurus against the 4 Da precursor mass window DIA acquisitions of complex cellular lysate from *Gnmt* -/- and *Gnmt* WT livers, we were able to identify with >1% global peptide FDR in ≥4/12 (≥33%) of samples and localize with >1% FLR 14 mono-methyl peptides, 161 di-methyl peptides, 28 tri-methyl peptides, 141 acetylated peptides, and 32 succinylated peptides (Figure 6C). The same cohort of samples was assayed against the PTM enriched DIA Assay Library in OpenSWATH 2.3.0 with >1% FDR, identification in ≥4/12 (≥33%) of samples, and >1% FLR localization in Thesaurus. This resulted in localization and quantitation of 9 mono-methyl peptides, 17 di-methyl peptides, 13 tri-methyl peptides, 76 acetylated peptides, and 14 succinylated peptides (Figure 6C). Additionally, we were able to localize 128/171 (74.6%) of the modified peptidoforms quantified in OpenSWATH with >1% FLR in Thesaurus and found 121 quantifiable peptidoforms in common between Thesaurus and OpenSWATH (Figure 6D).

### Regulatory Role of Protein Methylation During Translation: mRNA Stability in NASH

Two NASH models which knock out *Mat1a* and *Gnmt* (*Mat1a* -/- and *Gnmt* -/-, respectively) were analyzed to determine the role that differential methylation potential plays in NASH by manipulating SAMe concentration. Since cellular methylation potential is decreased overall in *Mat1a* -/- and increased in *Gnmt* -/-, we focused on methylated residues and proteins which show this trend. We found differentially methylated peptides in *Mat1a* -/- and *Gnmt* -/- which are associated with protein translation in addition to differential cytosolic and mitochondrial ribosomal protein levels, the protein machinery needed for translation. As an example, there is a cluster of differential methylated peptides at known amino acid residues in *Mat1a* -/- and *Gnmt* -/- which are proteotypic for three RNA binding proteins involved in translation (EF1A1), RNA stability (poly(A)-binding protein (PABP1)), and transcription and RNA stability (Heterogeneous nuclear ribonucleoprotein U (HNRPU)) (Figure 7A). Based on differential methylation patterns of these residues in both mutants we hypothesize that decreased methylation in these proteins will result in decreased messenger RNA stability and subsequent decreased protein translation which should be observed as differentially expressed proteins between *Gnmt* -/- and *Mat1a* -/- NASH. 627 proteins which were decreased in *Mat1a* -/- (FDR < 0.01, log2 FC < −0.32) and 735 proteins which were significantly increased in *Gnmt* -/- (FDR < 0.01, log2 FC > 0.32) (Table S7). Of these differentially dysregulated proteins we found significant enrichment in: mitochondrial electron transport chain complexes (ETC, OxPhos) (Figure 7C,D), mitochondrial ribosomes (MRPLs) and cytosolic ribosomes (Figure 7B). We also analyzed our previously published microarray data^31,32^ from the same models of NASH as a measure for mRNA stability of transcripts from the pathways which were differentially altered at the protein level (Figure S9 A, B). The trends in mRNA levels of the altered pathways (decrease in mRNA in *Mat1a* -/- and increase of mRNA in *Gnmt* -/-) are concordant with our hypothesis that PABP1 methylation effects their mRNA stability. Quantifying methylation and total protein from the same DIA acquisition allows for a much better understanding of how a pathway can be regulated by PTMs from a single experiment.

**Figure 7.**
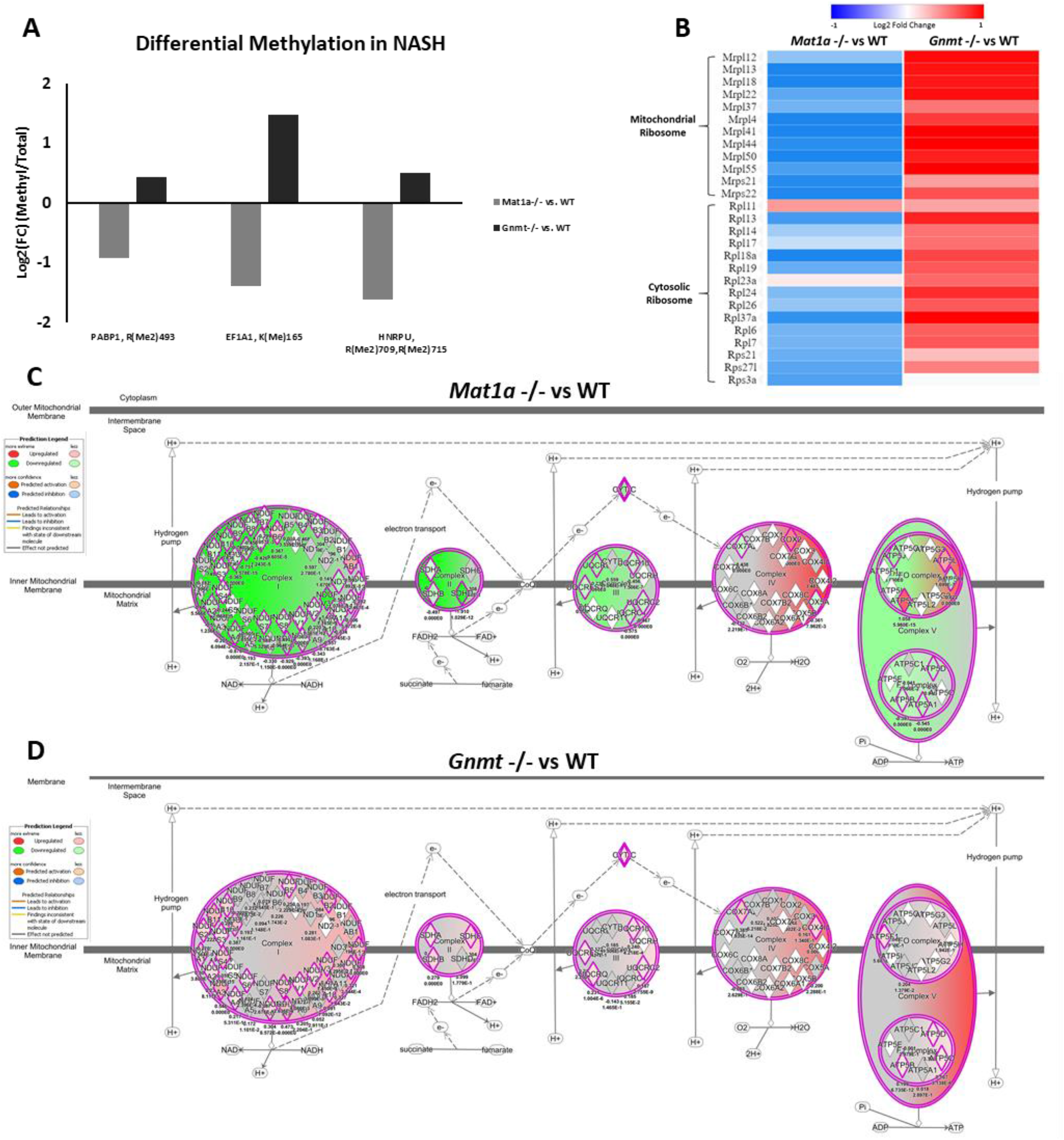
Methylation’s Differential Effect on Translation: **(A)** Quantification of the intensity of the MS2 TIC normalized methylated peptide divided by the intensity of then respective total protein, determined by unmodified peptides using MAPDIA in Gnmt -/- and wild-type littermates and Matla -/- and wild-type littermates **(B)** Total protein quantitation of mitochondrial ribosomes and cytosolic ribosomes determined using MAPDIA in Gnmt -/- and wild-type littermates and Mat1a -/- and wild-type littermates. All data has a FDR<0.01. **(C)** Predicted inhibition of oxidative phosphorylation complex 1,2,3, and 5 in *Matla -/-* vs WT from total protein using Ingenuity Pathway Analysis (activation z-score −4.61) **(D)** Predicted activation of oxidative phosphorylation complex 1.2,3, 4. and 5 in *Gnmt -/-* vs WT (activation z-score 6.403) All data has FDR<0.01.

### Disparate Mechanisms Drive Decreased Acetylation in Murine NASH Models

Protein acetylation was decreased in both models of NASH (Figure S10A) compared to controls. This could be due to two different mechanisms which should also be observed by changes in the proteome. First, metabolomic analysis of Acetyl-CoA levels from livers from *Mat1a* -/- and wild-type littermates show a 54% decrease in Acetyl-CoA (p-value = 0.0002) in *Mat1a* -/-, (Figure S10D) supporting globally decreased acetylation in *Mat1a* -/-. Interestingly, there is no change in the total protein quantity of sirtuins SIRT2 and SIRT5, a class of protein de-acetylases, in the proteomics analysis of *Mat1a* -/-. Second, in *Gnmt* -/-, there is a 61% increase of the deacetylase SIRT3 at the total protein level (FDR = 5.4E-18). In addition to increased SIRT3 levels in *Gnmt* -/- mice, we also find significant increases in total protein abundance of IDH2 and SOD2, both of which are known to be activated via SIRT3 mediated deacetylation^51–53^ (Figure S10B). In our acetyl-peptide dataset, we found a known target of SIRT3 deacetylation, ECHA K411-Acetyl, significantly decreased in *Gnmt* -/-, in agreement with SIRT3 activation in *Gnmt* -/- (Figure S10C). In addition at the total protein level, we see substantial increase of all five complexes of the electron transport chain in *Gnmt* -/-, also consistent with downstream effects of SIRT3 mediated deacetylation^51–53^(Figure 7D).

## Discussion

In this study, we have developed a method utilizing 4 Dalton precursor mass windows that can differentiate modified peptidoforms from each other and their corresponding unmodified peptide sequence. Combined with Thesaurus, updated to provide localization scores for PTMs involved in NASH *in-vivo*, now available as a community resource, our approach provides both total protein quantification and quantitative data for seven PTMs from the same sample and acquisition run.

A previous study by *Sidoli et. al.*, applied a small precursor mass windows approach to analyze methylation from histone enriched lysate using multiplexed 6 Dalton precursor mass windows with the ability to differentiate a +2 precursor but not a +3 precursor from its unmodified form.^37^ We compared multiplexed 4 and 6 Dalton precursor mass windows using the approach outlined in *Sidoli et al.*, to non-multiplexed windows on complex liver lysate. Although we were able to detect methylated peptides, the intensity was on average 10-fold lower than in non-multiplexed windows method. Additionally, fragment ions do not overlap perfectly causing issues with identification and quantitation of peaks (Figure S11). Combined with the long time needed to de-multiplex windows with current tools, we chose to not use that method for this study. We have shown using a mixture of synthesized unmodified and methylated peptides that we have the ability to physically separate 100% of the observable +2 and +3 methylated precursors from their unmodified forms. Furthermore, we applied this method to complex cellular lysate and have shown that it can simultaneously quantify protein methylation, Lys acetylation and succinylation, as well as unmodified peptides in a single LCMS injection.

Growing interest in studying residue specificity of PTMs using DIA approaches has created a need for tools that can accurately localize modified amino acid residues and provide a PTM-site localization score. It has recently become possible to computationally assess the strength of PTM localization of peptidoforms which contain positional isomers using tools which score fragment ions distinguishing between isomers to assess the correct site of modification.^8,10,11^ However, these tools were all tested and optimized for phospho-peptide positional isomer localization using synthesized peptides and designed to be used on phospho-peptide enrichments; a significantly less complex sample matrix than complex cellular lysate. We chose to use Thesaurus 0.9.4 for modified peptidoform localizations, a graphical interface easily operated by an end-user which has the ability to score and localize Lys and Arg PTM’s pursued in the study of NASH increasing the confidence of PTMs identified with our approach.

We have shown that our 4 Da precursor mass window DIA method provides linear protein quantitation in both the 60- or 120-minute LC gradient and that the quantitation of the method significantly outperforms common DIA acquisition methods for low abundant peptides, and therefore will be more useful to quantify the signatures of low abundant PTMs in complex lysate. We recommend using a 120-minute LC gradient, as with the described method’s cycle time; more points across the chromatographic peak can be achieved. Furthermore, we have shown that small precursor isolation windows together with spectrum-centric data analysis workflow used in our method outperforms MS1 centric workflows in its ability to quantify low abundant peptides, including endogenous low abundant and modified peptides (Figure S12). In the majority of cases, as the abundance of a peptide decreases, the co-eluting precursor trace is lost. Instead of relying on a co-eluting precursor trace to differentiate a methylated peptide from an unmodified peptide, our method physically separates the unmodified and methylated peptides by acquiring their precursors into separate mass windows allowing for the quantitation of low abundant modified peptides without the presence of a MS1 trace.

To better understand the role of PTMs in NASH, we tested this methodology in differentially methylated mouse models where the primary methyl donor, SAMe, is genetically manipulated.^33,35^ We linked differential methylation potential and altered acetylation together with total protein changes to propose a hypothesis of regulatory function of PTMs on proteins involved in mRNA stability, protein synthesis, and mitochondrial function found in the *Mat1a* -/- and *Gnmt* -/- NASH models. Differential methylation detected in RNA binding proteins indicates that decreased methylation could impair mRNA stability and protein translation. PABP1 is known to be controlled by post-translational modifications and the proline-rich linker region is crucial for its polymerization, for mature mRNA nuclear export, protein synthesis, and its capacity to bind and stabilize mRNA.^54–56^ We identified PABP1 R493 di-methyl, a known site in this critical proline-rich linker region with unknown function and not described in NASH. Differential methylation of PABP1 via destabilized mRNA could be responsible for dysregulated protein synthesis in observed significantly altered pathways between *Gnmt* -/- and *Mat1a* -/- NASH. (Figure 7B-D).

While cellular methylation potential is solely controlled by SAMe concentration, acetylation is not genetically manipulated in either of these *in-vivo* NASH models and has multiple points of control. Globally decreased protein acetylation observed in both NASH models could be due to two different mechanisms. Acetylation mainly occurs non-enzymatically,^29,57^ therefore observed decreased concentration of Acetyl-CoA may explain the global protein acetylation decrease in Mat1a -/-. Alternatively, downregulation of protein acetylation could be driven by increased deacetylase activity of sirtuins.^58,59^ While there is no increase in observable sirtuins abundance in Mat1a -/-, we observe targets concordant with (SIRT3) activation in Gnmt -/-. We find alterations in downstream targets known to be activated upon SIRT3 deacetylation,^51–53,57,58^ including a substantial increase in total protein quantity of all five mitochondrial complexes in *Gnmt -/-*, which could be regulated by both increased SIRT3 activity and PABP1 hypermethylation.

There are some limitations with our approach where in isolated cases manual inspection for a site-specific transition or co-eluting precursor trace is needed to verify peptidoform identification. Nonetheless, quantitation with 4 Da precursor mass windows physically separates peptidoforms in the mass spectrometer which allows for the use of all transitions to identify and quantify respective peptidoforms, a significant benefit over large precursor mass window DIA where only few site-specific transitions can be quantified, often leading to decreased accuracy of quantitation. Additionally, large precursor mass windows can lead to ambiguity, especially in cases where only one site specific is transition detected (Supplemental Results, Figure S5). Finally, care should be taken when comparing tri-methylated and acetylated peptides, a challenging task with most MS approaches. See **Supplementary Discussion** for additional comments on the above limitations.

In conclusion, the small precursor mass window DIA approach outlined in this study which is suitable for both TripleTOF and Orbitrap mass spectrometers along with the corresponding publicly available PTM-enriched mouse liver peptide assay library provides an easily adoptable framework needed for quantification and localization via Thesaurus of protein methylation, acetylation, and succinylation as well as total protein quantification in any DIA proteomic experiment without the need for enrichment of each experimental sample. This approach only requires enough sample for one DIA acquisition which opens up the possibility for studying PTMs from small experimental samples like tissue biopsies where there isn’t enough sample for traditional PTM immunoprecipitations paving the way towards understanding the biological significance and functional diversity of PTM containing proteoforms.

## Author Contributions

The manuscript was written through contributions of all authors. All authors have given approval to the final version of the manuscript.

## Acknowledgment

The authors thank Helen Choi Robinson for helping with figure design. This work was supported by USA National Institutes of Health (NIH) grants R01DK107288 (SC Lu and J Van Eyk), NIH GM110174 and AI118891 (B Garcia), and Agencia Estatal de Investigación MINECO SAF 2017-88041-R, ISCiii PIE14/00031 CIBERehd-ISCiii, and Severo Ochoa Excellence Accreditation SEV-2016-0644) (J Mato).

## ABBREVIATIONS

PTMs: Post translational modifications
NASH: Non-alcoholic steatohepatitis
MS: Mass spectrometry
DDA-MS: Data dependent acquisition
DIA-MS: Data independent acquisitions
IPF: Inference of peptidoforms
SAMe: S-Adenosylmethionine
LC: Liquid chromatography
*Mat1a*: Methionine Adenosyltransferase A1
*Gnmt*: Glycine *N*-methyltransferase
FLR: False localization rate
SCX: Strong cation exchange
FDR: False discovery rate
SIL: Stabile isotopically labeled
TIC: Total ion current
RT: Retention time
fm: Femtomoles
AUC: Area-under-the-curve

