## Supplementary Material for "Quantitative Method for Assessing the Role of Lysine & Arginine Post-Translational Modifications in Nonalcoholic Steatohepatitis"

^1^Advanced Clinical Biosystems Research Institute, The Smidt Heart Institute, Cedars Sinai Medical Center, Los Angeles, CA 90048, USA; ^2^Department of Genome Sciences, University of Washington, Seattle, WA, USA; ^3^Department of Systems Biology, Columbia University, New York, NY, USA; ^4^Buck Institute for Research on Aging, Novato, CA 94945, USA;­­ ^5^Department of Biochemistry and Biophysics, Epigenetics Institute, University of Pennsylvania School of Medicine, Philadelphia, PA 19104, USA; ^6^DeepDIA, USA ^7^CIC bioGUNE, Centro de Investigación Biomédica en Red de Enfermedades Hepáticas y Digestivas (Ciberehd), Technology Park of Bizkaia, 48160 Derio, Bizkaia, Spain; ^8^Division of Digestive and Liver Diseases, Cedars-Sinai Medical Center, Los Angeles, CA, 90048, USA

**Supplementary Results**

**Generation of In-vivo PTM DIA Assay Library**: Two mouse models of NASH where employed, each chosen due to their extreme methylation potential. *Gnmt* -/- mice have increased SAMe and are biologically hypermethylated while *Mat1a* -/- mice are SAMe deficient and therefore hypomethylated.^1,2^ First, we created a hypermethylated DIA assay library by DDA acquisitions of 5 SCX fractions from *Gnmt* -/- mouse livers. We supplemented this library with a DDA acquisition of a methyl-Lys immunoprecipitation from a hepatoma cell line derived from *Gnmt* -/- mice.^3^ This gave us an assay library of 808 methyl Arg and Lys peptides. We further supplemented the library with DDA acquisitions of a ‘one-pot’ acetyl-Lys and succinyl-Lys immunoaffinity enrichment of mitochondrial extract and whole liver lysate isolated from WT mouse liver as described in *Basisty et al*^4^ which provided us with 2428­­ succinylated peptides and 2547 acetylated peptides.

**Simultaneous Total Protein and 7 different PTM Quantification**: Total protein quantification was obtained at the same time as the PTM data because our DIA assay library was built with SCX fractionated complex cellular lysate, containing both modified peptides as well as unmodified peptides. Focusing on the total protein data from *Gnmt* -/- mice with NASH and comparing the 12 Dalton precursor mass window data to the 4 Dalton precursor mass window data, we achieve marginally better results when searching the 4 Dalton precursor mass window data through OpenSWATH. Assaying against the same SCX fractionated DDA library, 5,111 proteins, 26,293 peptides, and 341,241 transitions where identified with our 4 Dalton Precursor mass Windows. With the 12 Dalton Precursor Mass Window acquisitions, we matched 4,595 proteins, 22,782 peptides, and 304,365 transitions. (Figure S13).

**Comparison of the Performance of Alternate DIA Data Analysis Methods**: MS1 centric workflows like DIA-Umpire^5^ and the “DirectDIA” function in Spectronaut (Biognosys) use of MS1 masses in a “spectral-centric” workflow have the ability to differentiate a modified peptide from an unmodified peptide by the precursor mass alone. However, these tools rely on a coeluting precursor trace to accurately identify a peptide. Using the SIL peptides dilution series to mimic low abundant peptides in complex liver lysate and visualizing the results in Skyline we found cases whereas we lowered the concentration of SIL peptides, a coeluting precursor trace was lost while transitions remained. Although 100% of SIL peptides had a coeluting precursor trace at 2250 femtomoles (fm) on column in both the 60- and 120-minute gradient, as the concentration of SIL peptides decreased so did their coeluting precursor traces. When we approached 7 fm on column, only 36% had coeluting precursor traces in the 60 min gradient and 46% had coeluting precursor traces in 120 minute gradient (Figure 3 E,F). Accordingly, using MS1 traces of SIL peptides to determine linearity of their quantitation, we found a sharp decrease in accurate quantitation looking at the lower end of our dilution curve in 4 Da and 12 Da DIA acquisition methods (Figure S3, Table S3). Furthermore, looking at endogenous modified peptides in these same acquisitions from our DIA assay library, methylated, acetylated, or succinylated, we found 60.4% do not have a coeluting precursor trace, making them undetectable by “MS1-centric” approaches and in the case of methylated peptides undifferentiable from their unmodified peptidoform without 4 Da precursor mass windows (Figure S12 A,B). To further this point, we next analyzed DIA acquisitions of complex liver lysate using different data analysis software. We chose to use Spectranaut’s software suite as Spectranaut has the ability to analyze DIA data with a targeted library based method as well as DirectDIA, a MS1-centric library free approach. Assaying complex liver lysate DIA acquisitions against our PTM enriched DIA assay library resulted in quantitation of 188 endogenous modified peptides from our DIA assay library; methylated, acetylated, or succinylated. DirectDIA analysis of the same DIA acquisitions resulted in 28 endogenous modified peptides. We also assayed the same complex liver lysate acquired with 12 Da windows using DirectDIA which resulted in the 35 endogenous PTM containing peptides (Figure S12C, Table S4). Instead of relying on a coeluting precursor mass to identify a modified peptide and differentiate a methylated peptide from unmodified peptide, our method physically separates the unmodified and methylated peptides by acquiring their precursors into separate mass windows. In addition, using a PTM enriched DIA assay library, our method allows for the quantitation of low abundant modified peptides without a MS1 trace.

**Limited Number of Ambiguous Methylation Quantitation in DIA-MS**: Another important consideration is the extent of ambiguous transitions and their impact on methyl-peptide quantitation. In other words: peptides that are in the same precursor mass window as their unmodified form, and where transitions from the peak of a co-eluting unmodified peptide are being assigned as methylated. Looking at complex cellular liver lysate from *Gnmt ­*-/- mice acquired with 12 Da precursor mass windows, we have an example of an ambiguous identification; the same peak is being called for the methylated peptide and the unmodified peptide with a single site-specific transition (b7) for the methyl-site identified (Figure S5A,B). Looking at the same sample acquired with 4 Da precursor mass windows, although we can still detect the unmodified peptide, we no longer can detect the methylated peptide, leading us to believe that the quantified intensity of the methylated peptide with 12 Da windows is incorrect (Figure S5C,D). Additionally, due to this co-eluting site-specific transition, FLR algorithms will incorrectly assign a high probability to this ambiguous modified peptidoforms. 4 Dalton precursor mass windows separate 100% of +2 and +3 charged precursors from their methylated forms leading to zero modified peptides with ambiguous transitions leading to incorrect quantitation in complex lysate while 12 Da precursor mass windows have over 50% of methylated peptides which fall in the same precursor mass window and are likely to contain transitions which will lead to ambiguous FLR assignment and quantitation. (Figure S5E). In addition to ambiguous quantitation, false positive identification also occurs for methyl and dimethyl peptides where the unmodified peak was being called as methylated based on the same criteria but in this case zero site specific transitions are observed (Figure S6).

**Supplementary Discussion**

Lys (and Arg) residues can be modified by multiple PTMs and crosstalk between these PTMs constitutes a major regulatory mechanism of protein function.^6–8^ Despite its importance in biology, PTM crosstalk is difficult to study and currently not possible without enrichment of each respective PTM. Using our approach, we can assay 946 Lys residues which contain multiple PTMs; 24 Lys residues contain multiple methyl forms (Figure S14A) and 922 Lys residues where crosstalk between acetylation, succinylation, or methylation occurs, 872 of which have crosstalk between acetylation and succinylation (Figure S14B). Having the ability to simultaneously identify and quantify peptides with different Lys PTM crosstalk in one DIA run will further our understanding of the role of PTM crosstalk in biological systems.

**Limitations:** DIA-MS relies on a pre-existing peptide assay library and compares the MS data obtained from each experimental sample to this library. As such, PTMs not contained within this assay library are not assayed in experimental samples using our approach and the library is species specific and often sample (e.g. cell type) specific. We have shown the limitation of current library free approaches at detecting PTMs (Figure S12C). To this point, this is why so much work went into maximizing PTMs in our murine PTM enriched DIA assay library. As with all MS analysis including DDA analysis of complex samples, the proteome depth is limited unless fractionation of some form is used. The hereby proposed methodology is limited to the dynamic range of unfractionated complex lysate, and we are most like only able to detect the highest abundant methylated, succinylated, and acetylated peptides. However, it is important to recognize that, DIA-MS approaches display a number of favorable properties, by systematically fragmenting all precursor ions in precisely defined (peptidoform-tailored) width of mass to charge windows, and concurrently overcome the stochastic ion selection of data-dependent-acquisition (DDA), providing larger completeness to the dataset. The ability to quantitate highly abundant modified peptides as well as total protein quantitation from the same DIA acquisition provides a complementary and analytically valuable alternative despite the limited depth. In addition, we have shown that our method performs quite well with peptide of low abundance and in fact outperforms existing DIA acquisition methods for low abundant peptides at concentrations similar to biological PTMs in complex lysate (Figure 3).

Although unmodified peptides always fall in different mass windows as methylated peptides, if the peptide contains a methionine the oxidized species may end up in the same precursor mass window as the methylated peptide. In this case, a site-specific transition or co-eluting precursor trace is needed to verify peptidoform identification. Retention time difference is a valuable method for peptidoform differentiation, as peptides containing an oxidized methionine elute before their corresponding unmodified peptide by 2.37% ACN ‘gradient units’.^9^ Using the retention time difference from the unmodified peptide, a co-eluting precursor trace, or a site specific transition, one can differentiate a methylated and oxidized peptidoforms.

Additional consideration should be taken when comparing tri-methylated and acetylated peptides, a challenging task with most MS approaches. There are two major places for ambiguous or misidentification when differentiating these isobaric modifications in library-based DIA-MS. 1) During DDA acquisitions for DIA library build the correct modification needs to be assigned by computational spectral searching algorithms and 2) During DIA acquisitions care needs to be taken to prevent misidentification of acetylated and tri-methylated peptidoforms.

Trimethylated and acetylated peptides have isobaric nature (Tri-Methyl = 42.0470 Da, Acetyl = 42.0106 Da), which may confound peptide characterization by means other than high-resolution or high-mass accuracy mass spectrometry. To limit the misassignment of tri-methylated and acetylated spectra during DIA library build, we utilized immunoprecipitations for both acetyl and methyl peptides, enriching for each modification separately and only searching the immuno-enrichment products for their respective target. We additionally used 10ppm precursor and fragment mass tolerances during the spectral library build to further confidence of a correct identification. However, it must be acknowledged that precursor mass tolerance of 10 ppm makes trimethylation distinguishable from acetylation only at a precursor masses lower than 3,600 Da, where their mass can be distinguished on the third decimal point. For example, precursor ions at charge state 2+, 3+, +4 larger than 1800m/z, 1200m/z and 900m/z, respectively, would need additional attention. Peptide acetylation results in increased retention in reversed-phase chromatography, while methylation, including trimethylation, shows little change in retention when compared to unmodified peptide sequence, hence RT of these modified peptides can be used to determine the nature of the modification.^10^ When applying 120-min gradient in LCMS runs acetylated peptides elute 1.5 ~ 3.5 min later than the unmodified isoforms of the same peptides.^11–14^ Therefore mass accuracy of the methodology is applicable to modifications residing on peptides with masses well below above mentioned limits and for the larger species elution time inspection should be implemented additionally. To limit misidentification in our DIA acquisitions, we extracted peaks in 150 second windows on either side of library matches, sufficient to differentiate these isobaric modifications with longer gradients as long as spectra are correctly assigned during DDA library build.

Finally, a methylated Lys or Arg often inhibits trypsin’s ability to cleave.^15^ In order to determine the stoichiometric site specific ratio of a modified peptidoform, we ideally would use an enzyme that would give a methylated peptidoform without any missed cleavages and normalize to the intensity of the unmodified peptidoform which would give a better representation of the quantity of the modified peptidoform than our approach. However, to make our method more amendable for adoption by the proteomics community, we use trypsin for our study as it is currently the most commonly used endoproteinase but, only after quantifying the number of potential missed-cleavages due to the Lys/Arg being modified. Importantly, missed cleavages of unmodified peptides do still occur even with Trypsin/LysC mix 5-10% of the time.^16^ An unmodified peptide with a missed cleavage can occur due to incomplete digestion, although generally it would be at a much lower abundance than the tryptic version of the peptide. Since the PTMs in this study are also low abundant, it would be possible to misidentify an incompletely digested unmodified peptide containing a missed cleavage with a PTM containing peptide especially if it had only few low-abundant site-specific transitions. However, this is not the case when we apply our 4 Da window method which physically separates the unmodified from the modified version within the MS instrument. There are also plenty of methylated peptides that do not have a missed cleavage in our database, and which have been reported. For example, in Geoghegan et al, 2015^17^ they find (682/5781) 11.8% of methylated peptides without a missed cleavage (tryptic peptides). In these situations, one would run into the issue where the tryptic unmodified and methylated peptide would occur and co-elute in the same precursor mass window, unless 4Da window DIA acquisitions were applied.

**Supplementary Methods**

**SCX Fractionation Sample Preparation:** To establish a broad methylation peptide assay library, 500µg of lysate from mouse livers (n=1/condition) was digested as described above. Each sample were de-salted (Oasis HLB, 10 mg sorbent cartridges) and Strong Cation Exchange (SCX) fractionated as described in *Kooij et al*.^18^ with some modifications. De-salted peptides were dried and re-suspended in SCX buffer A (7 mM KH_2_PO_4_, 30% ACN, pH 2.65). The SCX columns were prepared by adding 100 mg of SCX bulk media (polysulfoethyl aspartamide, Nest Group) suspended in 1ml of 30% ACN to an empty 1 ml cartridge (Applied Separations), allowing the slurry to settle and sealing the column bed with an additional frit. The column was wet with 3 ml of 80% ACN, followed by a 3 ml H_2_O wash and then equilibrated with 6 ml of SCX buffer A. Samples were loaded on the columns and the flow through was collected. Columns were washed two more times with SCX buffer A and the flowthroughs were combined as the first fraction. Samples were sequentially eluted with 1 ml SCX buffer A containing four different salt concentrations (10mM KCL, 40 mM KCl, 60mM KCL, and 150 mM KCl). All 5 fractions were dried, re-suspended in 1ml 1% FA, and desalted on using HLB plates (Oasis HLB 30µm, 5mg sorbent, Waters).

**SCX Fractionation MS Acquisition Orbitrap Fusion Lumos:** For DDA-MS analysis, a Orbitrap LUMOS Fusion mass spectrometer (Thermo Scientific) was equipped with an EasySpray ion source and connected to Ultimate 3000 nano LC system (Thermo Scientific). Peptides were loaded onto a PepMap RSLC C18 column (2 µm, 100 Å, 150 µm i.d. x 15 cm, Thermo) using a flow rate of 1.4 µL/min for 7 min at 1% B (mobile phase A was 0.1% formic acid in water and mobile phase B was 0.1 % formic acid in acetonitrile) after which point they were separated with a linear gradient of 5-20%B for 45 minutes, 20-35%B for 15 min, 35-85%B for 3 min, holding at 85%B for 5 minutes and re-equilibrating at 1%B for 5 minutes. Each sample was followed by a blank injection to both clean the column and re-equilibrate at 1%B. The nano-source capillary temperature was set to 300 °C and the spray voltage was set to 1.8 kV. MS1 scans the AGC target was set to 4x10^5^ ions with a max fill time of 50 ms. MS2 spectra were acquired using the TopSpeed method with a total cycle time of 3 seconds and an AGC target of 5x10^4^ and a max fill time of 22 ms, and an isolation width of 1.6 Da in the quadrapole. MS1 scans were acquired in the Orbitrap at a resolution of 60,000 FWHM from mass range 400-1000m/z. Precursor ions were fragmented using HCD with a normalized collision energy of 30% and analyzed in the Orbitrap at 15,000K resolution. Monoisotopic precursor selection was enabled and only MS1 signals exceeding 50000 counts triggered the MS2 scans, with +1 and unassigned charge states not being selected for MS2 analysis. Dynamic exclusion was enabled with a repeat count of 1 and exclusion duration of 15 seconds.

**SCX Fractionation MS Acquisition SCIEX TripleTOF 6600:** For DDA-MS analysis, the 6600 TripleTOF (Sciex) was connected to Eksigent 415 LC system that was operated in micro-flow mode. The mobile phase A was comprised of 0.1% aqueous formic acid and mobile phase B was 0.1% formic acid in acetonitrile. Peptides were pre-loaded onto the trap column (ChromXP C18CL 10 x 0.3 mm 5 µm 120Å) at a flow rate of 10 µL/min for 3 min and separated on the analytical column (ChromXP C18CL 150 x 0.3mm 3 µm 120Å) at a flow rate of 5 µL/min using a linear A-B gradient composed of 3-35% A for 60 min, 35-85% B for 2 min, then and isocratic hold at 85% for 5 min with re-equilibrating at 3% A for 7 min. Temperature was set to 30^o^C for the analytical column. Source parameters were set to the following values: Gas 1 = 15, Gas 2 = 20, Curtain Gas = 25, Source temp = 100, and Voltage = 5500V.

DDA MS1 scans were acquired at 45,000 FWHM, using a dwell time of 250 ms in the mass range of 400-1250 m/z of 100 counts per second were selected for fragmentation. DDA MS2 scans were acquired in high-sensitivity mode at 15,000 FWHM with dynamic accumulation with a dwell time of 25 ms for ions ranging from +2 to +5 using rolling collision energy and a collision energy spread of 5. Ions were excluded for fragmentation after one occurrence for a duration of 15 seconds.

**Methyl-Enrichment Sample Preparation:** A cell pellet (~1 ml volume) of a hepatoma cell line derived from a *Gnmt* -/- mouse^3^ was resuspended in 4 ml urea lysis buffer (8 M urea, 100 mM NaCl, 50 mM Tris pH 8) supplemented with 1X protease inhibitors (Halt, Thermo) and sonicated on ice with a probe sonicator (Fisher Sonic Dismembrator Model 100, setting 2, 3 x 15 sec). Whole cell extract was then cleared by centrifugation for 10 mins at 4°C and 14,000 rpm (15,996 x g). A total of 10 mg of protein was reduced with 10 mM dithiothreitol at 56°C for 30 mins and alkylated with 50 mM iodoacetamide for 40 mins at room temperature in the dark. This incubation protocol is unique to this set of samples, while the rest of the study’s samples were subjected to reduction at 37°C. After adding 4 volumes of 25 mM Tris pH 8, proteins were digested with trypsin (1/200 ratio by mass) overnight at 37°C. The digest was acidified by addition of trifluoroacetic acid (TFA) to 1% and then desalted with SepPak cartridges (3cc, 200 mg). Dried peptides were resuspended in 1 ml of phosphate-buffered saline (PBS) and dibasic sodium phosphate was added to neutralize the pH (~ 50 mM final concentration). Insoluble debris was pelleted by centrifugation for 10 mins at 17,000 x g at room temperature and the supernatant was used for immunoprecipitation of methylated peptides. Peptide concentration was estimated based on UV absorbance. Anti-mono-methyl lysine and anti-di-methyl lysine antibodies (Proteintech Group Inc.) using a synthetic peptide library immunogen consisting of CX6KX6, where X is any amino acid except cysteine and the central lysine is unmodified, mono-methylated, or di-methylated was used to further enhance the peptide library. A total of 50 µg of each antibody per 1 mg of input peptides were combined and incubated with magnetic protein A beads (GE, 70 µl slurry per 112 µg antibody) for several hours on a rotator at 4°C. The charged beads were washed 4 x 1 ml with PBS and then incubated with 1 ml (~3.8 mg) of peptides overnight at 4°C with rotation. The beads were washed 1 x 1 ml with PBS, 1 x 1 ml with PBS supplemented with 0.8 M NaCl, 1 x 1 ml with PBS, and 1 x 1 ml with water. Captured peptides were eluted by incubation with 100 µl of 1% TFA for 5 mins at room temperature and then desalted with C18 stage tips prior to analysis by LC-MS/MS.

**Methyl-Enrichment MS Acquisition:** Peptides were analyzed by LC-MS/MS using a nano EasyLC 1000 system (Thermo) fitted with a fused silica column (Polymicro Tech, 75 µm i.d. x 20 cm) packed with ReproSil-Pur 120 C18-AQ (3 µm, Dr. Maisch GmbH) and positioned in line with an Orbitrap Fusion mass spectrometer (Thermo). The chromatography run consisted of a 120 min gradient increasing from 2% to 40% solvent B over 105 mins, 40% to 80% solvent B over 15 mins, and then holding at 80% solvent B for 10 mins. Water containing 0.1% formic acid and acetonitrile containing 0.1% formic acid served as solvents A and B, respectively. The mass spectrometer was programmed to perform MS1 scans in positive profile mode in the Orbitrap at 120,000 resolution covering a range of 300-1500 m/z with a maximum injection time of 100 ms and AGC target of 1e6. MS2 scans on the most intense ions, fragmented by HCD, were performed in the Orbitrap in positive centroid mode at 15,000 resolution with a maximum injection time of 200 ms, target of 1e5, and NCE of 27. Quadrupole isolation was enabled with an isolation window of 2 m/z. Dynamic exclusion was set to 30 sec. MS2 scans were performed over a 3 sec cycle time.

**Acetyl and Succinyl Enrichment Sample Preparation:** Mitochondrial extracts and total protein lysate were prepared from mouse tissue as previously described.^4^ Briefly, crude mitochondrial fractions were isolated from the livers of SIRT5-/- (C57BL/6) mice by differential centrifugation. Total protein lysate was also isolated from the livers of WT (C57BL/6) mice. Protein concentration of mitochondrial extracts and total protein lysate was determined by BCA assay and 1 mg of protein per sample were brought to equal volumes in 8M urea in 50 mM TEAB buffer and vortexed for 10 min. Samples were reduced with 20 mM Dithiothreitol (DTT) for 30 min at 37C, alkylated in 40 mM iodoacetamide for 30 min at room temperature, diluted 10-fold in 50mM TEAB, and digested overnight at 37C with trypsin used at 1:50 enzyme protein. Digestion was quenched with formic acid and samples were desalted with Oasis HLB 10 mg Sorbent Cartridges, vacuum concentrated to dryness, and resuspended in 1.4mL immunoaffinity purification (IAP) buffer. Immunoaffinity enrichments of modified peptides were done using the PTMScan® Acetyl-Lysine Motif [Ac-K] Kit (#13416) and PTMScan® Succinyl-Lysine Motif [Succ-K] Kit (#13764) (Cell Signaling Technology, Danvers, MA), using the ‘one-pot’ affinity enrichment previously described.^4^ For one-pot enrichments, 1 mg peptide digests were incubated in a tube containing equal parts (~62.5 µg immobilized antibody) of both the succinyl- and acetyl-lysine antibodies.

**Acetyl and Succinyl Enrichment MS Acquisition:** Samples were analyzed by reverse-phase HPLC-ESI-MS/MS using the Eksigent Ultra Plus nana-LC 2D HPLC system (Dublin, CA) combined with a cHiPLC System, and directly connected to a quadrupole time-of-flight SCIEX TripleTOF 5600 or a TripleTOF 6600 mass spectrometer (SCIEX, Redwood City, CA). Typically, mass resolution in precursor scans was ~35,000 (TripleTOF 5600) or 45,000 (TripleTOF 6600), while fragment ion resolution was ~15,000 in ‘high sensitivity’ product ion scan mode. Injected peptide mixtures initially flow into a C18 pre-column chip (200 µm x 6 mm ChromXP C18-CL chip, 3 µm, 300 Å, SCIEX) and washed at 2 µl/min for 10 min with the loading solvent (H2O/0.1% formic acid) for desalting. Subsequently, peptides flow to the 75 µm x 15 cm ChromXP C18-CL chip, 3 µm, 300 Å, (SCIEX), and eluted at a flow rate of 300 nL/min using a 3 or 4 hr gradient using aqueous and acetonitrile solvent buffers.

For the creation of a spectral library for analysis by the library-based DIA workflow, data-dependent acquisitions (DDA) were carried out on all PTM enrichments to obtain MS/MS spectra for the 30 most abundant precursor ions (100 ms per MS/MS) following each survey MS1 scan (250 ms), yielding a total cycle time of 3.3 sec. For collision induced dissociation tandem mass spectrometry (CID-MS/MS), the mass window for precursor ion selection of the quadrupole mass analyzer was set to ± 1 m/z using the Analyst 1.7 (build 96) software.

**Supplementary Figures**

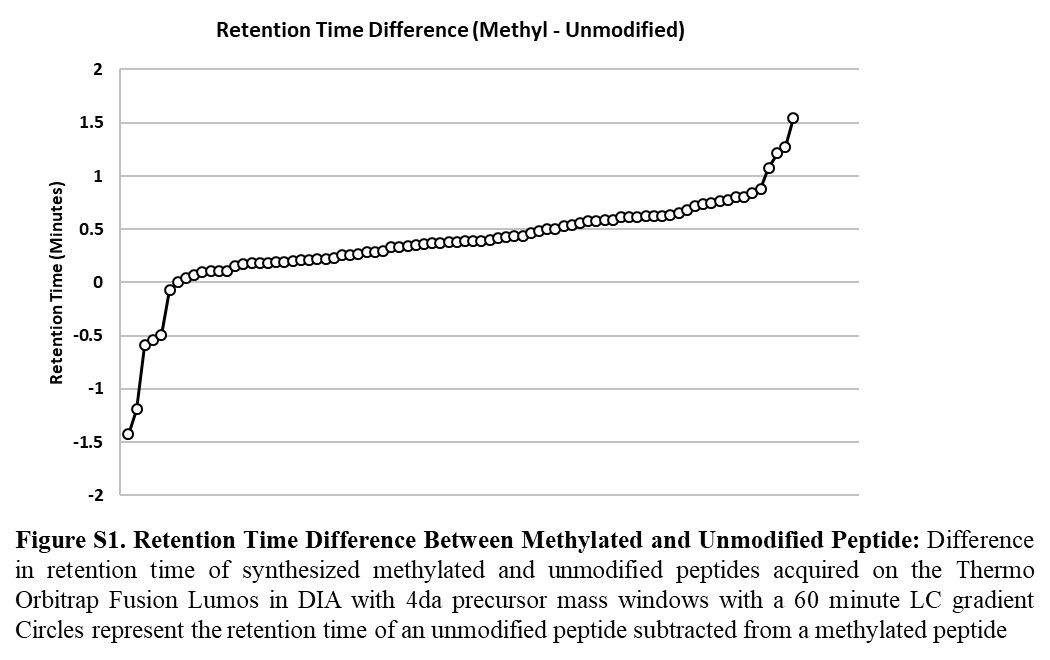

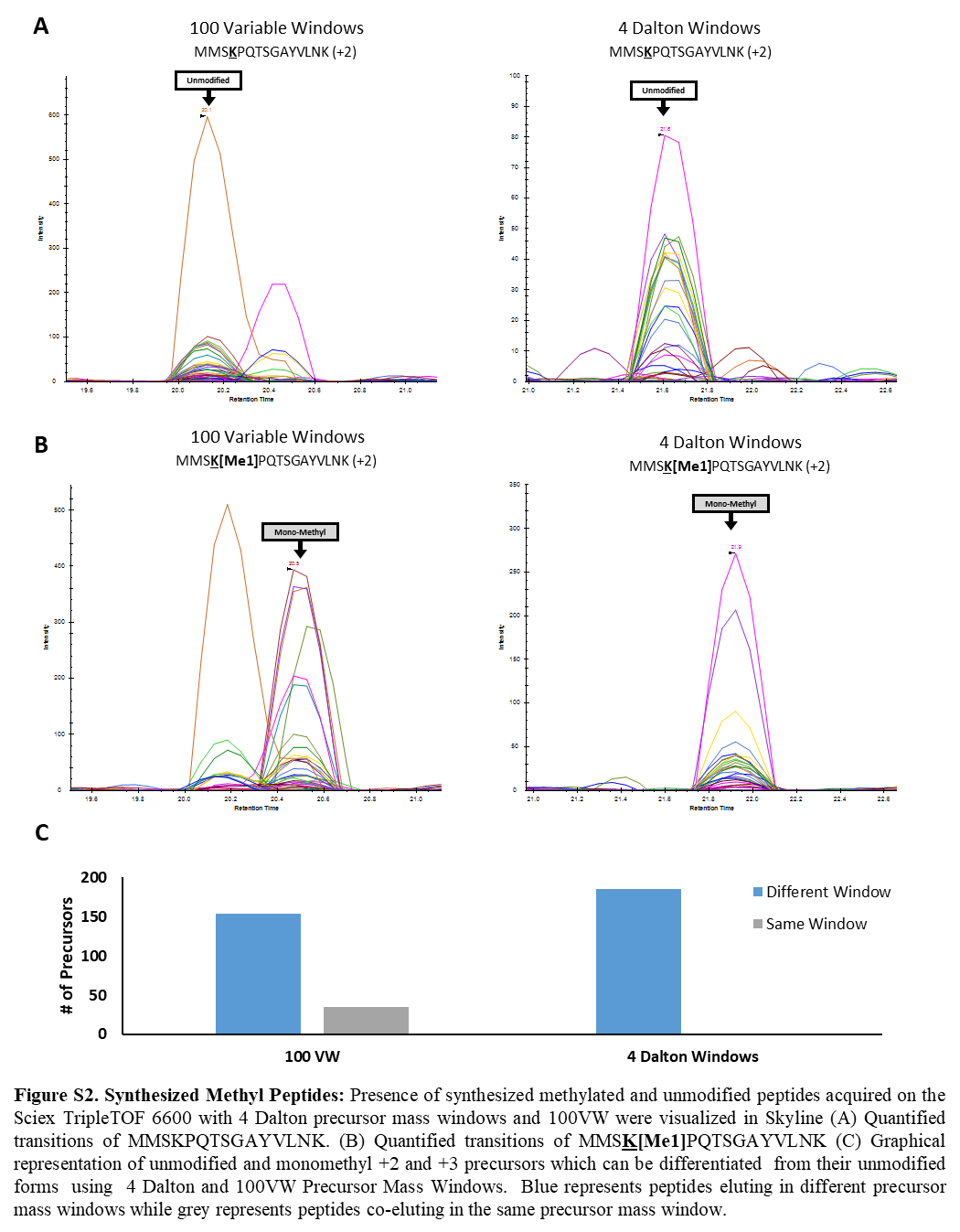

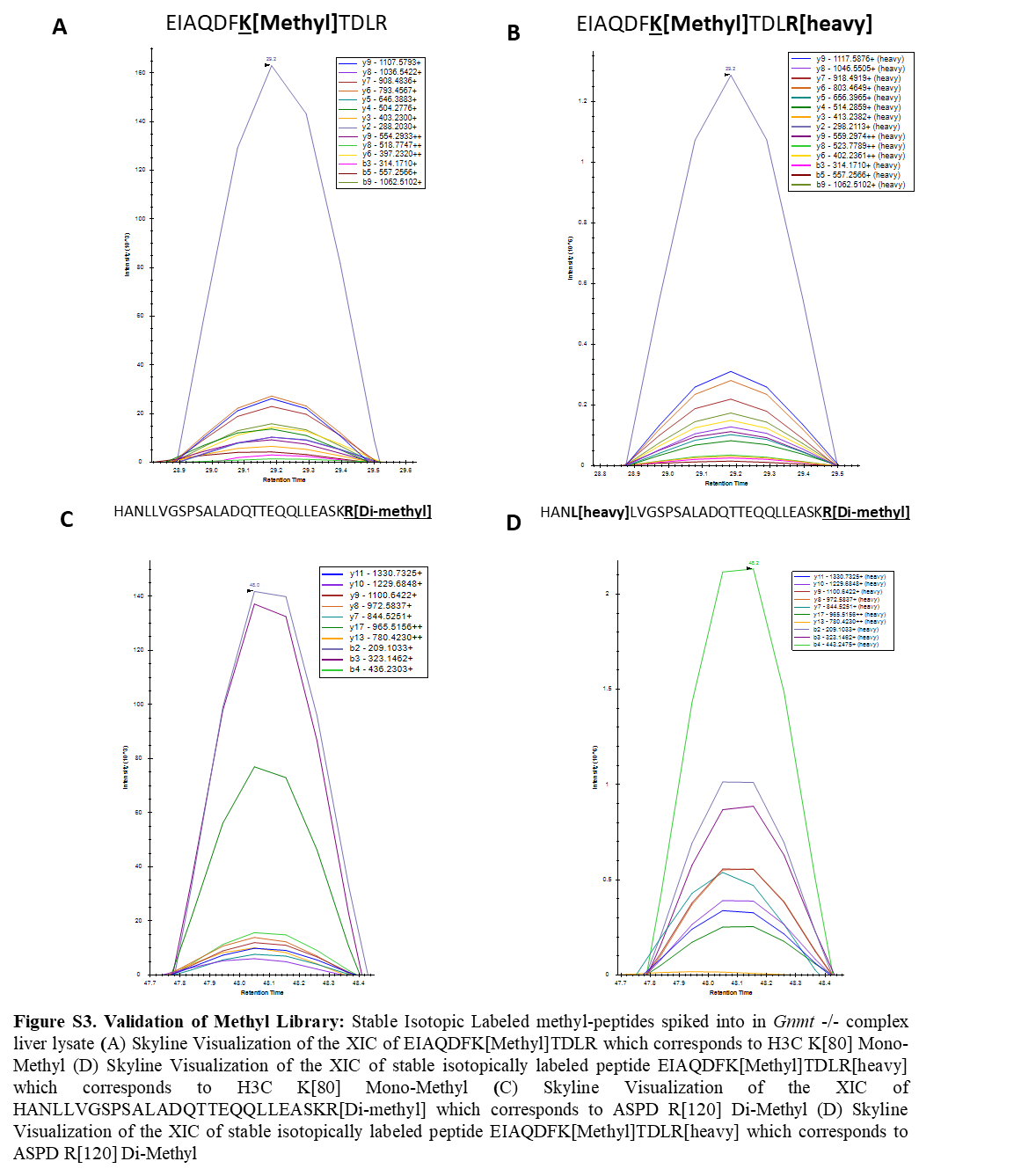

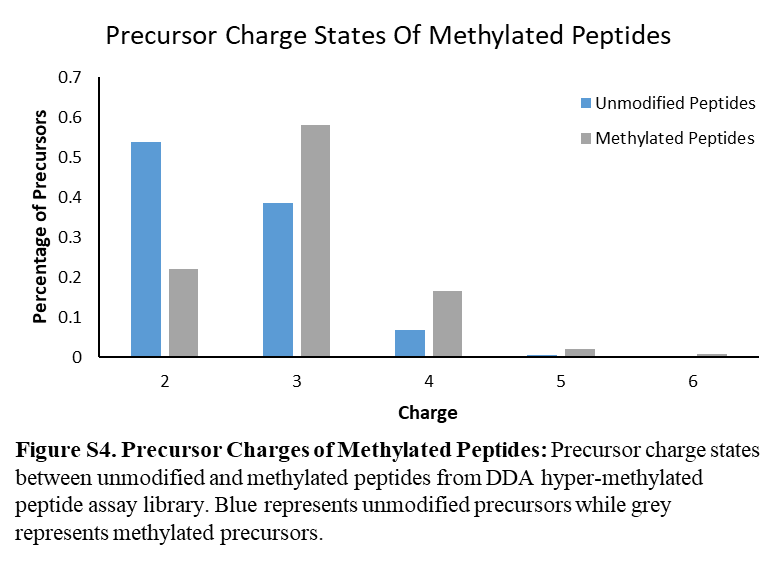

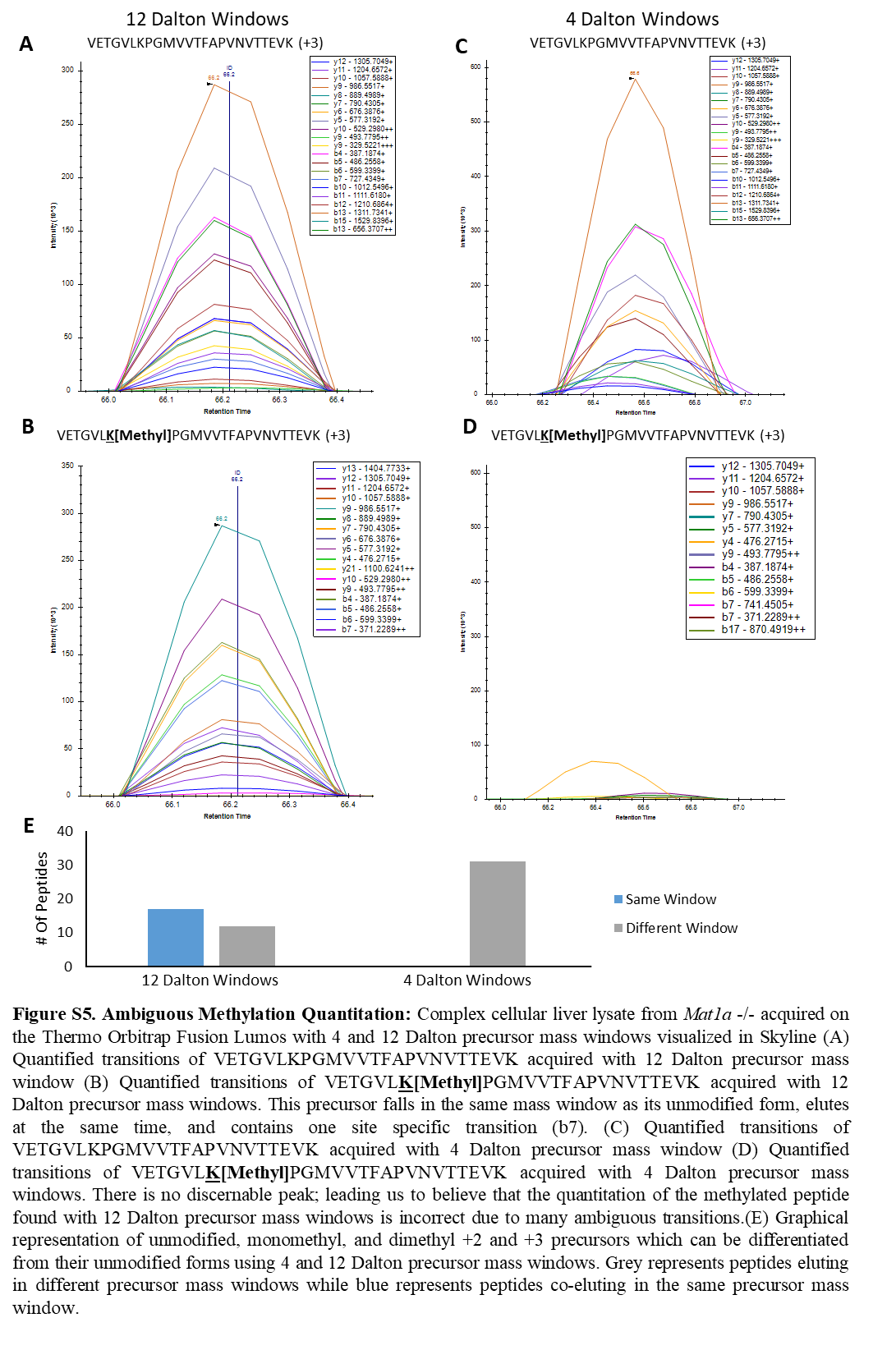

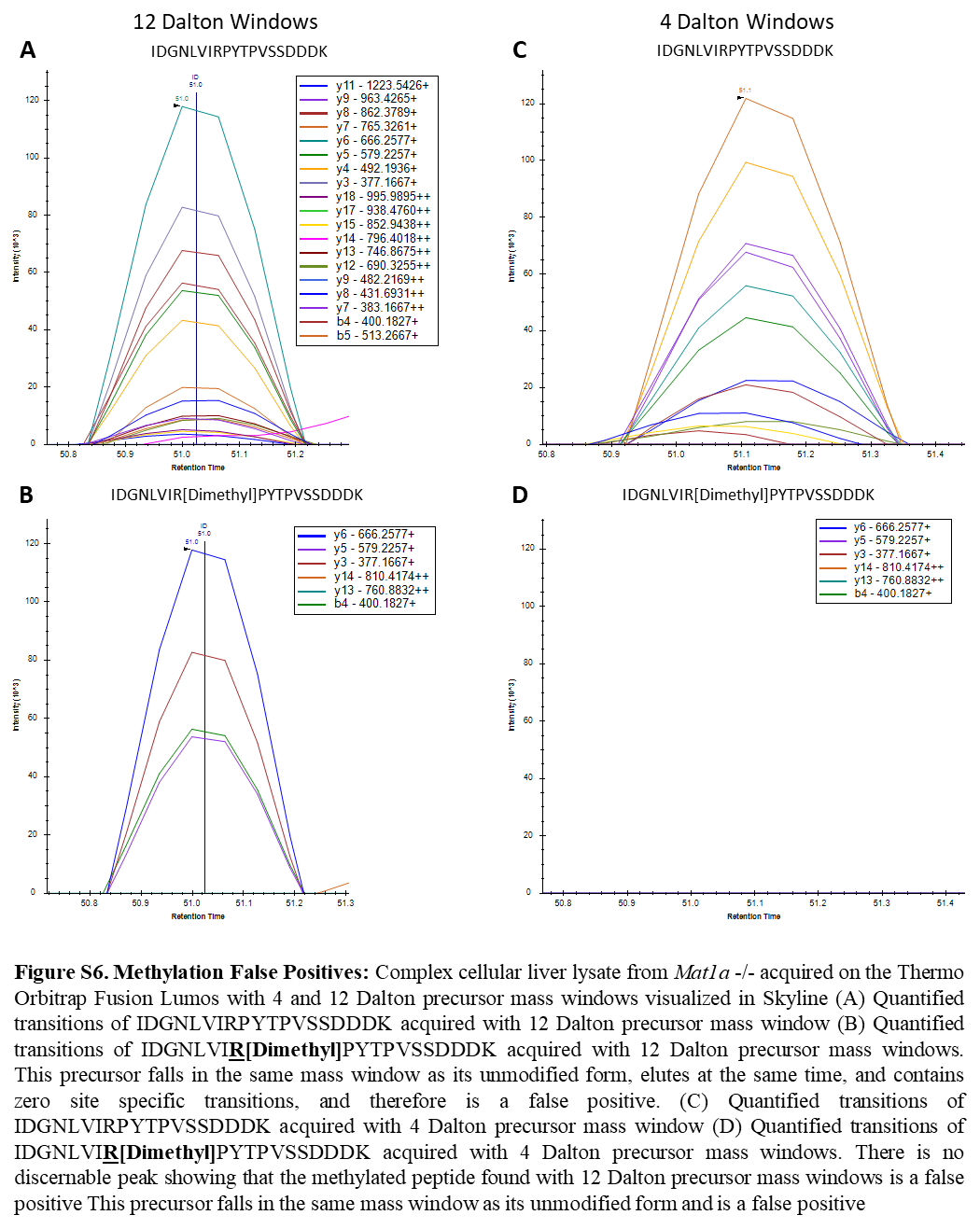

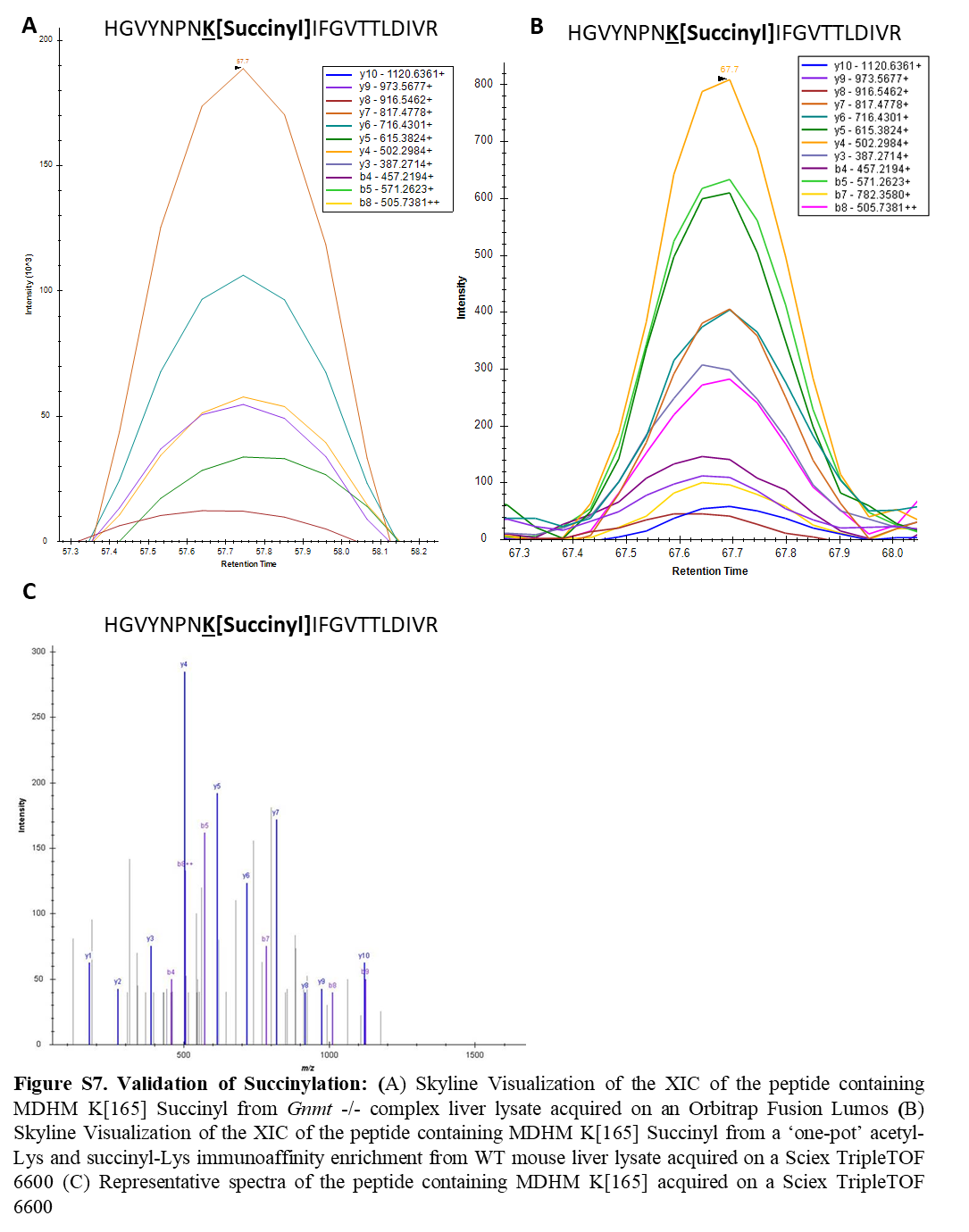

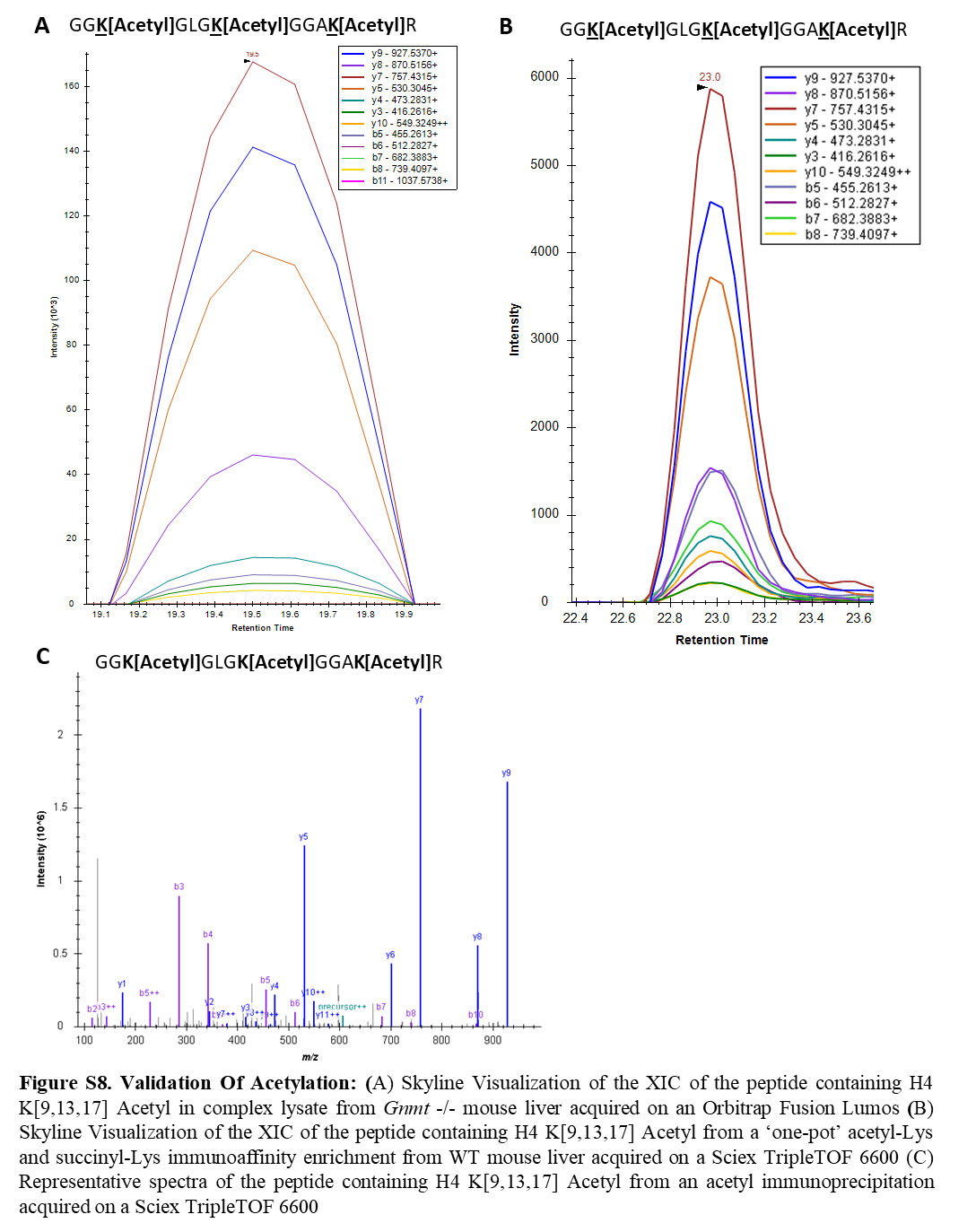

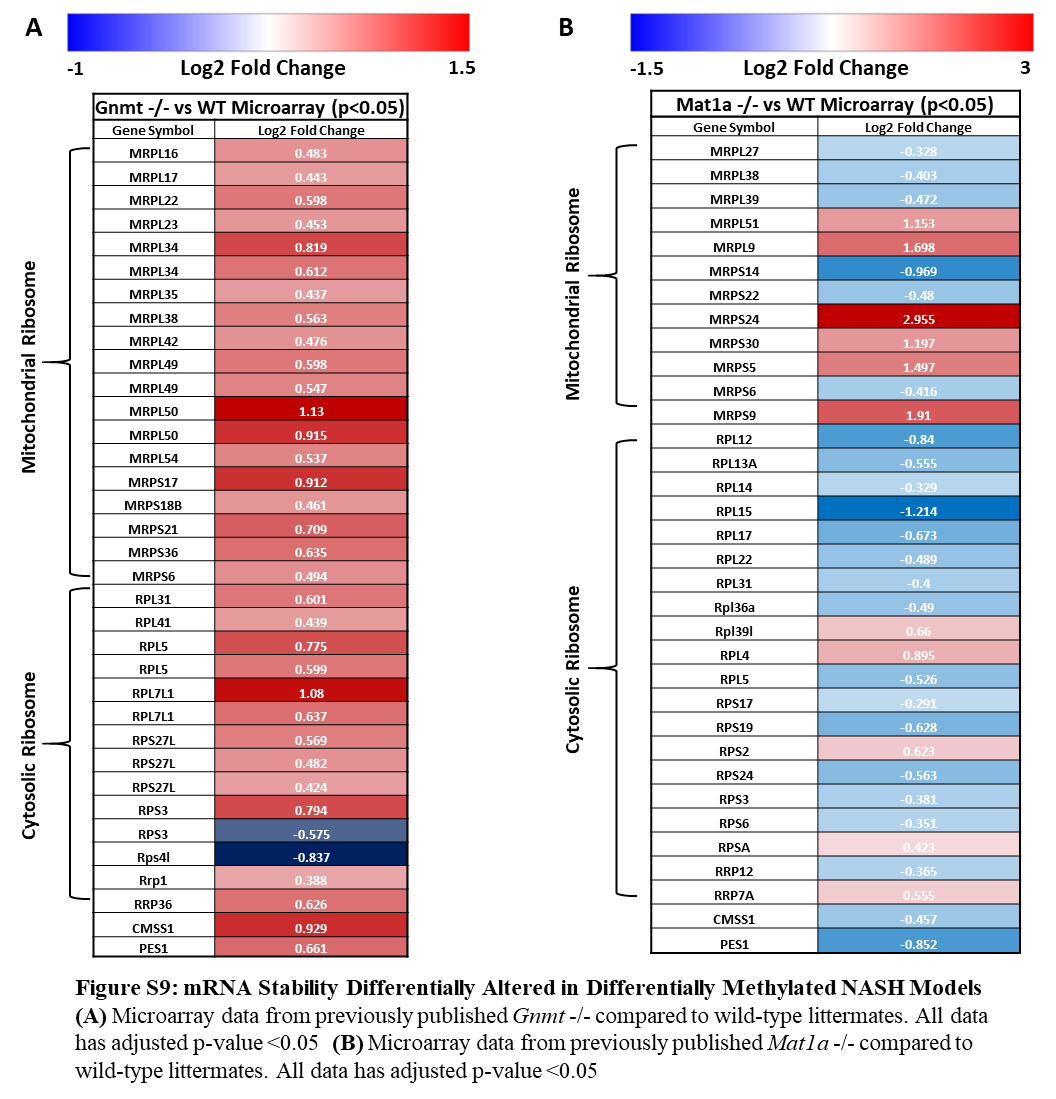

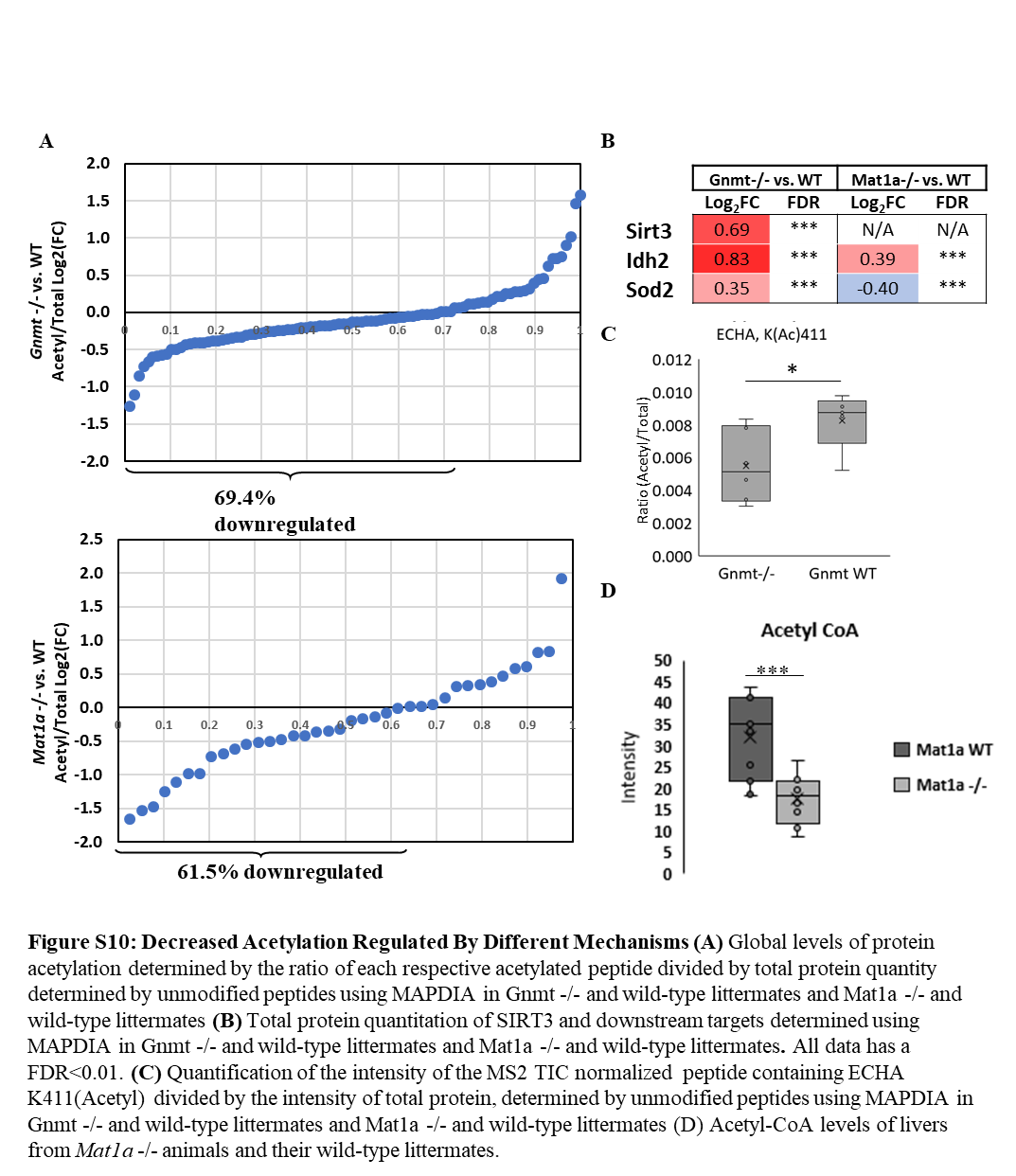

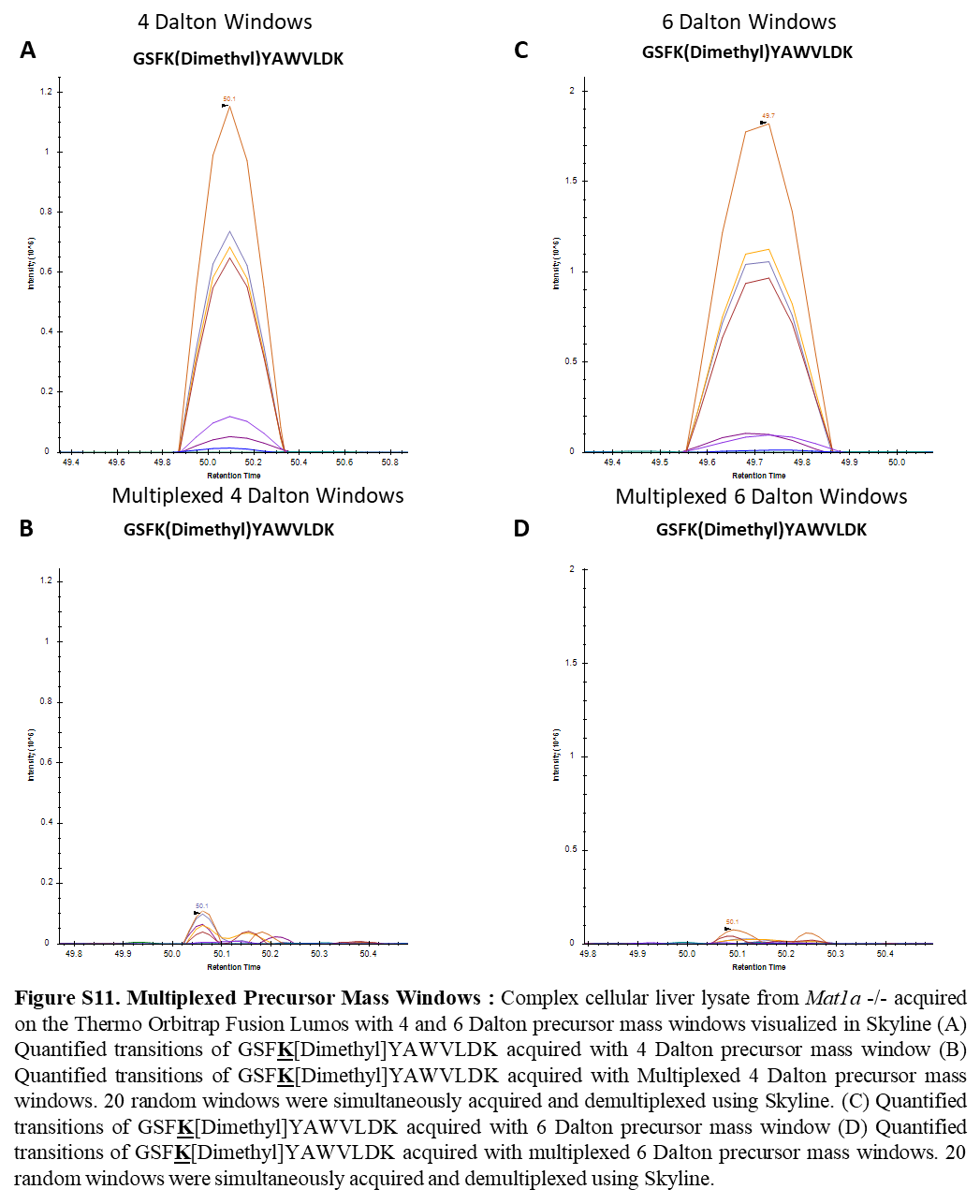

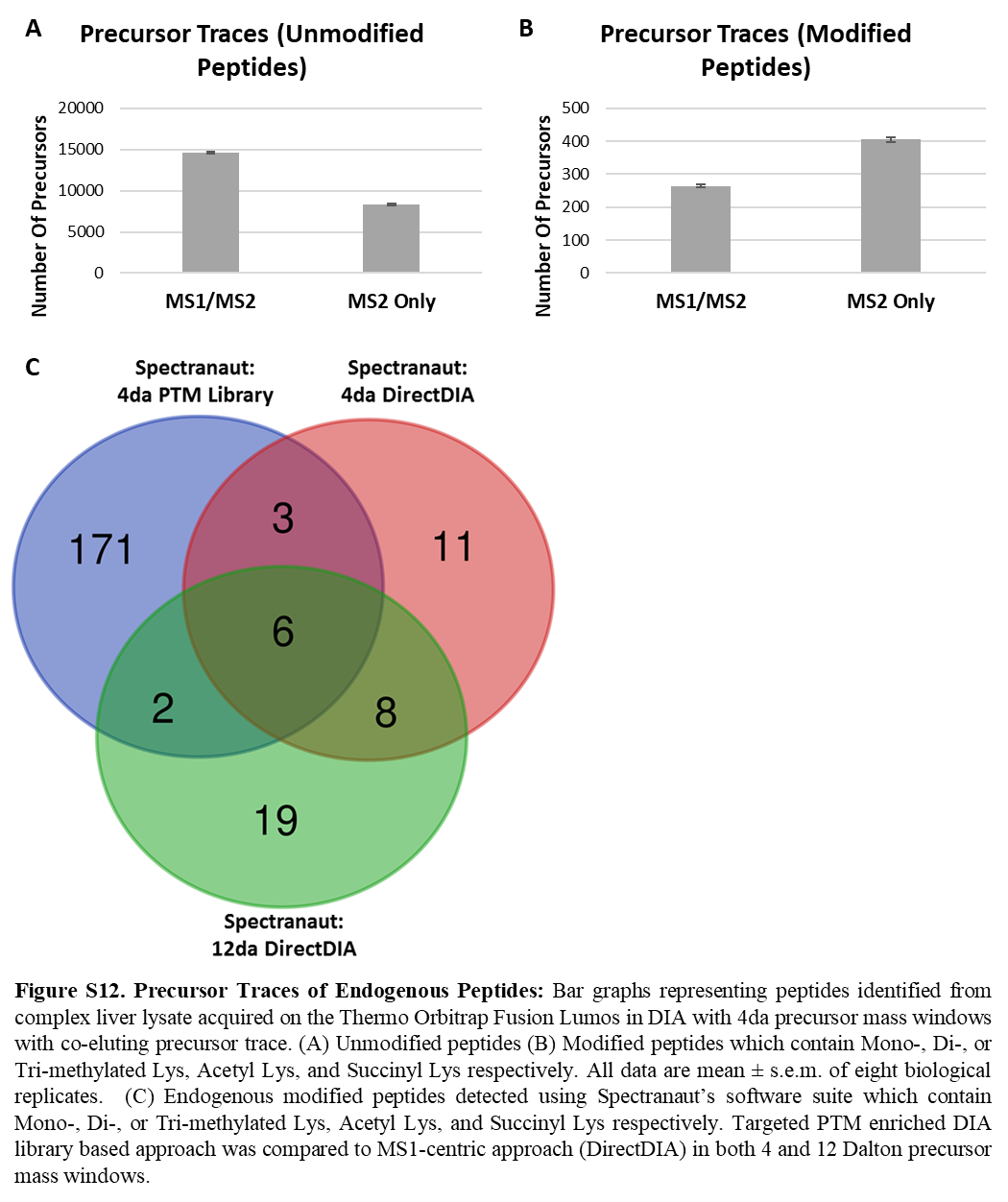

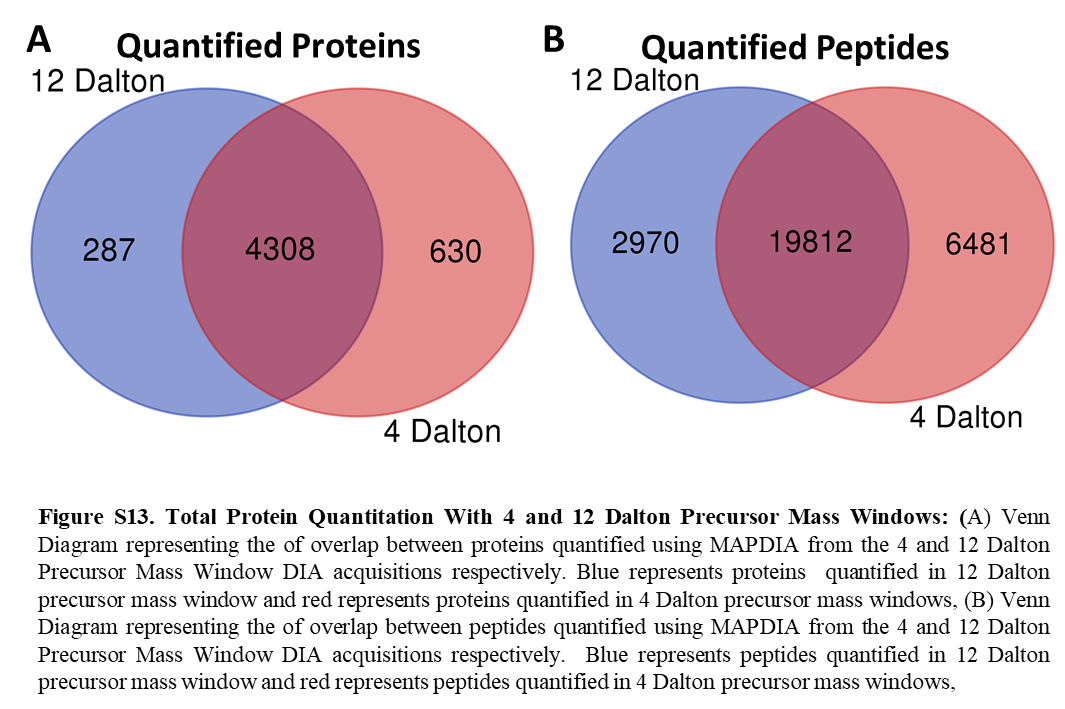

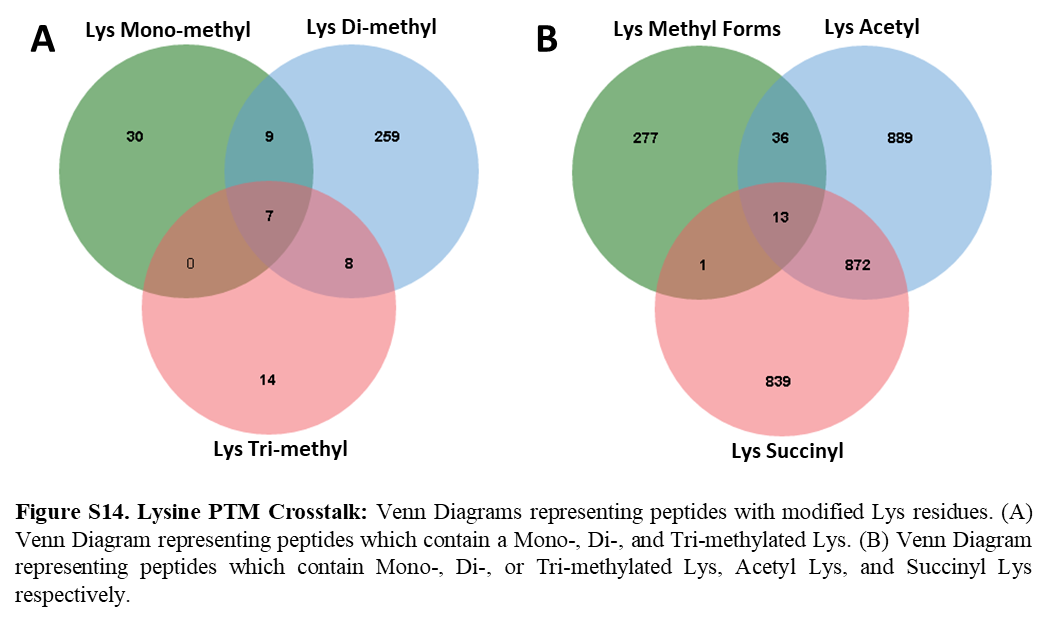
