## Supplementary material for "Quantitative Method for Assessing the Role of Lysine & Arginine Post-Translational Modifications in Nonalcoholic Steatohepatitis": Figures and Supplemental Figures

### Slide 1
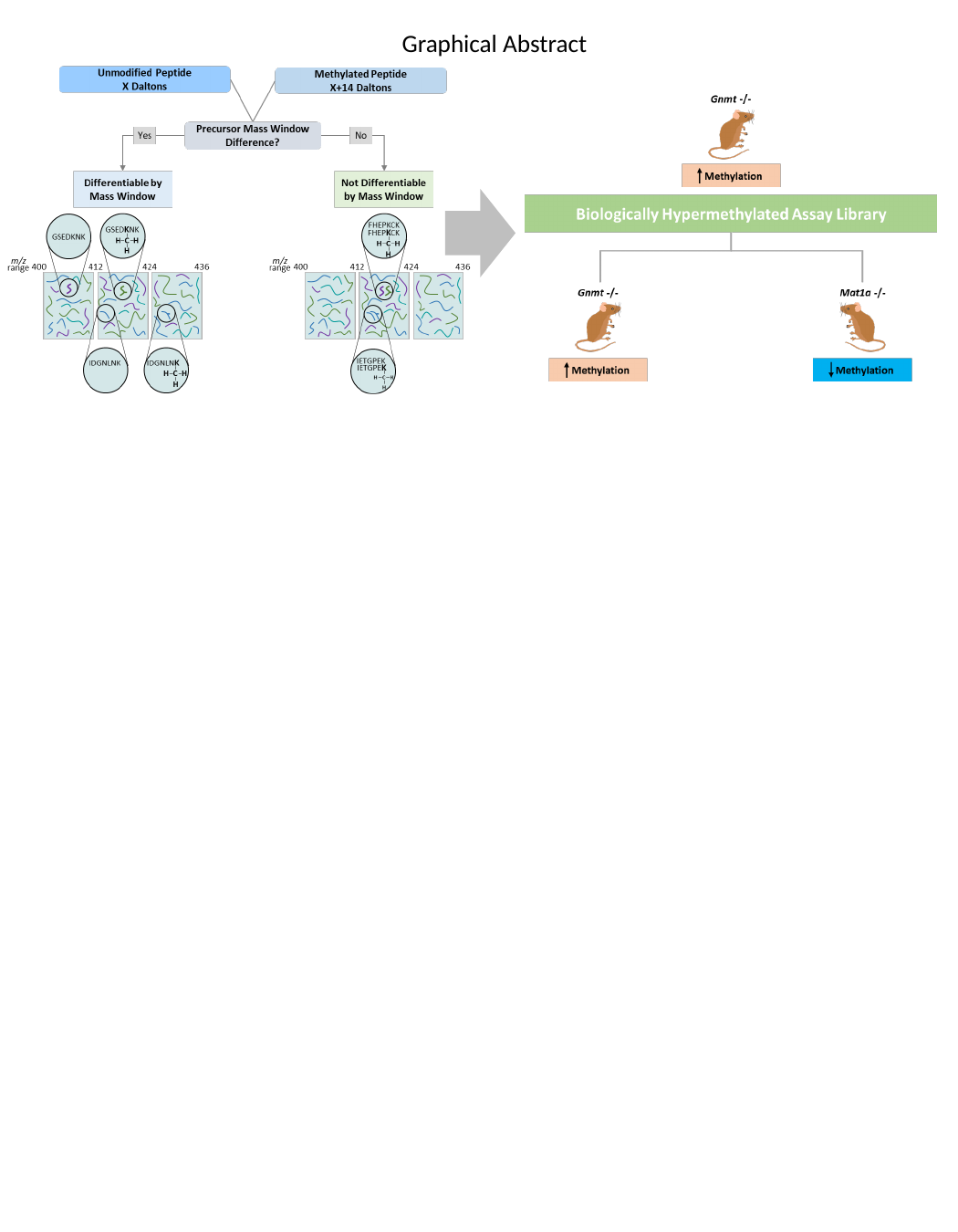

Graphical Abstract

### Slide 2
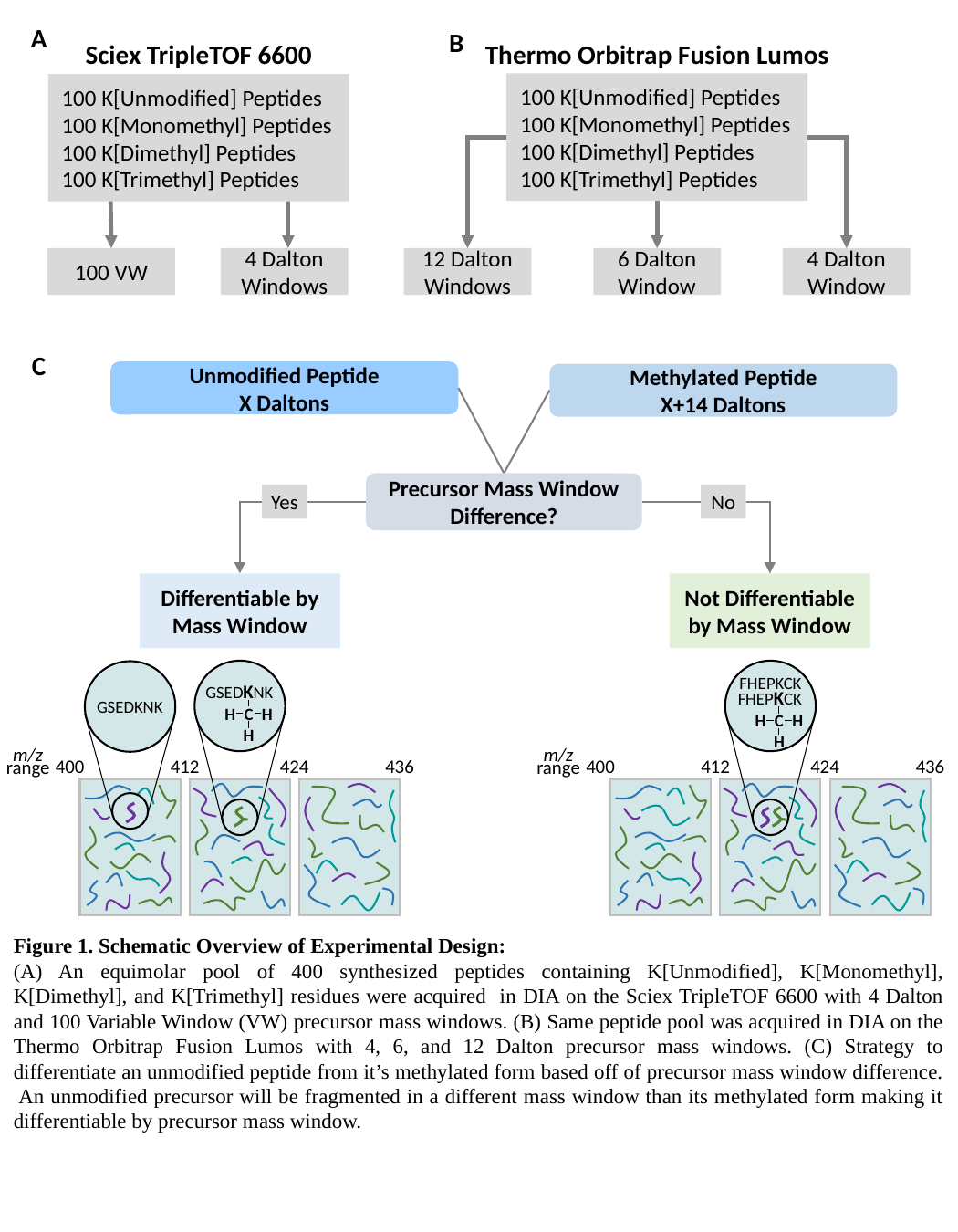

A
B
Sciex TripleTOF 6600
Thermo Orbitrap Fusion Lumos
100 K[Unmodified] Peptides
100 K[Monomethyl] Peptides
100 K[Dimethyl] Peptides
100 K[Trimethyl] Peptides
100 K[Unmodified] Peptides
100 K[Monomethyl] Peptides
100 K[Dimethyl] Peptides
100 K[Trimethyl] Peptides
100 VW
4 Dalton
Windows
12 Dalton Windows
6 Dalton Window
4 Dalton Window
C
Unmodified Peptide
X Daltons
Methylated Peptide
X+14 Daltons
Precursor Mass Window Difference?
Yes
No
Differentiable by Mass Window
Not Differentiable by Mass Window
GSEDKNK
H
C
H
H
GSEDKNK
m/z
range
400
412
424
436
FHEPKCK
FHEPKCK
H
C
H
H
m/z
range
400
412
424
436
Figure 1. Schematic Overview of Experimental Design:
(A) An equimolar pool of 400 synthesized peptides containing K[Unmodified], K[Monomethyl], K[Dimethyl], and K[Trimethyl] residues were acquired in DIA on the Sciex TripleTOF 6600 with 4 Dalton and 100 Variable Window (VW) precursor mass windows. (B) Same peptide pool was acquired in DIA on the Thermo Orbitrap Fusion Lumos with 4, 6, and 12 Dalton precursor mass windows. (C) Strategy to differentiate an unmodified peptide from it’s methylated form based off of precursor mass window difference. An unmodified precursor will be fragmented in a different mass window than its methylated form making it differentiable by precursor mass window.

### Slide 3
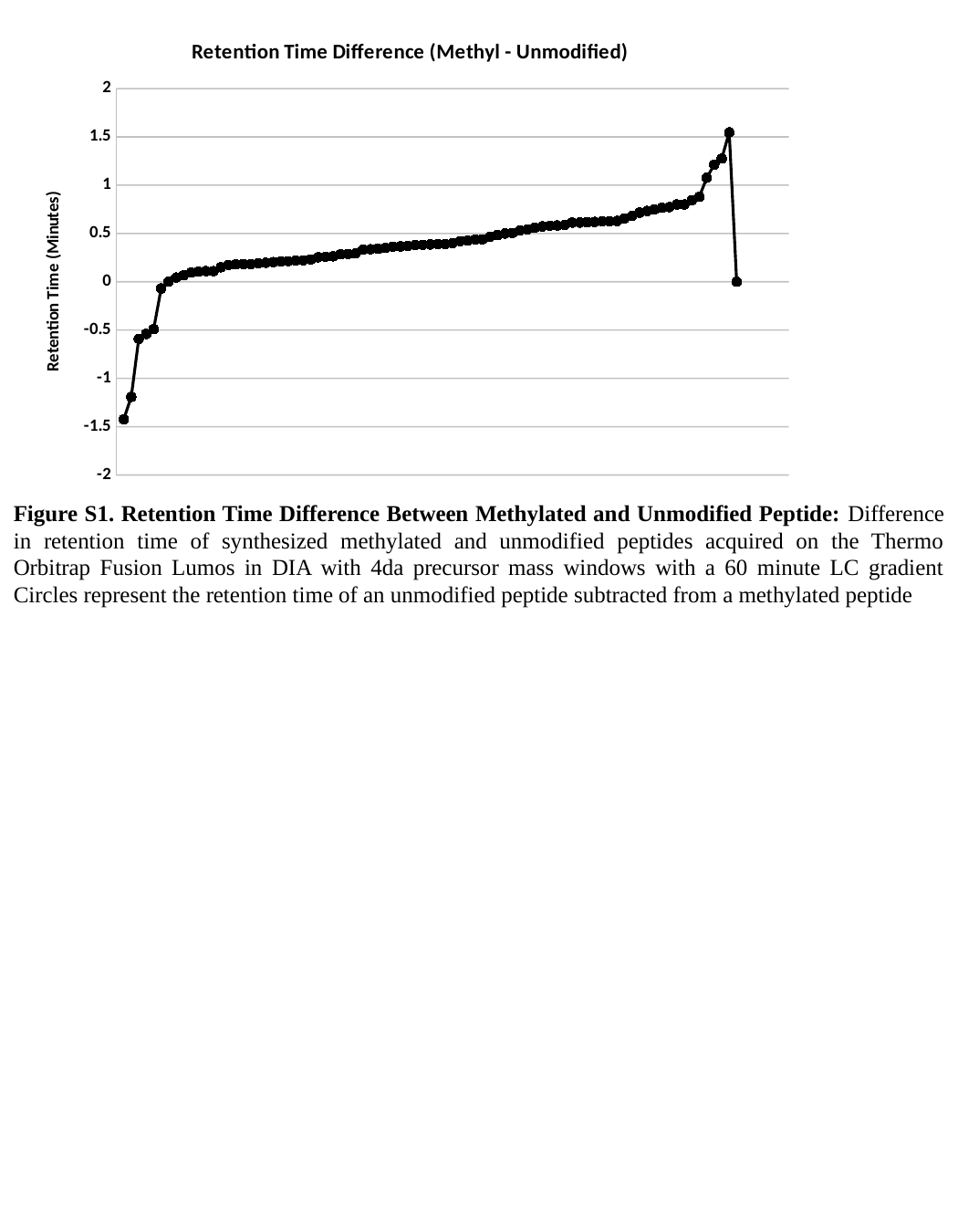

#### Chart: Retention Time Difference (Methyl - Unmodified)
| Category | |
|---|---|Figure S1. Retention Time Difference Between Methylated and Unmodified Peptide: Difference in retention time of synthesized methylated and unmodified peptides acquired on the Thermo Orbitrap Fusion Lumos in DIA with 4da precursor mass windows with a 60 minute LC gradient Circles represent the retention time of an unmodified peptide subtracted from a methylated peptide

### Slide 4
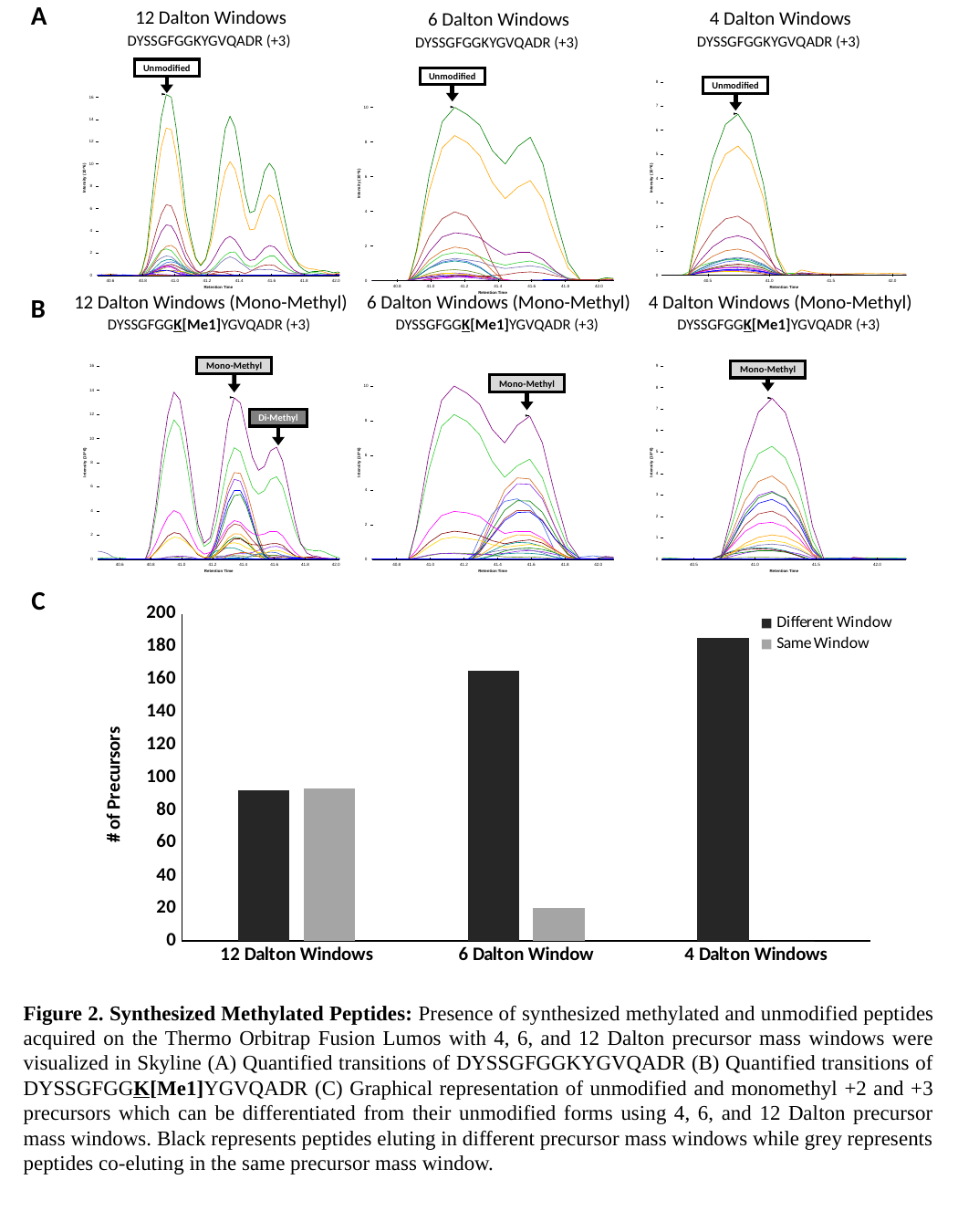

A
12 Dalton Windows
DYSSGFGGKYGVQADR (+3)
4 Dalton Windows
DYSSGFGGKYGVQADR (+3)
6 Dalton Windows
DYSSGFGGKYGVQADR (+3)
Unmodified
Unmodified
Unmodified
B
12 Dalton Windows (Mono-Methyl)
DYSSGFGGK[Me1]YGVQADR (+3)
6 Dalton Windows (Mono-Methyl)
DYSSGFGGK[Me1]YGVQADR (+3)
4 Dalton Windows (Mono-Methyl)
DYSSGFGGK[Me1]YGVQADR (+3)
Mono-Methyl
Mono-Methyl
Mono-Methyl
Di-Methyl
C
#### Chart
| Category | Different Window | Same Window |
|---|---|---|
| 12 Dalton Windows | 92.0 | 93.0 |
| 6 Dalton Window | 165.0 | 20.0 |
| 4 Dalton Windows | 185.0 | 0.0 |Figure 2. Synthesized Methylated Peptides: Presence of synthesized methylated and unmodified peptides acquired on the Thermo Orbitrap Fusion Lumos with 4, 6, and 12 Dalton precursor mass windows were visualized in Skyline (A) Quantified transitions of DYSSGFGGKYGVQADR (B) Quantified transitions of DYSSGFGGK[Me1]YGVQADR (C) Graphical representation of unmodified and monomethyl +2 and +3 precursors which can be differentiated from their unmodified forms using 4, 6, and 12 Dalton precursor mass windows. Black represents peptides eluting in different precursor mass windows while grey represents peptides co-eluting in the same precursor mass window.

### Slide 5
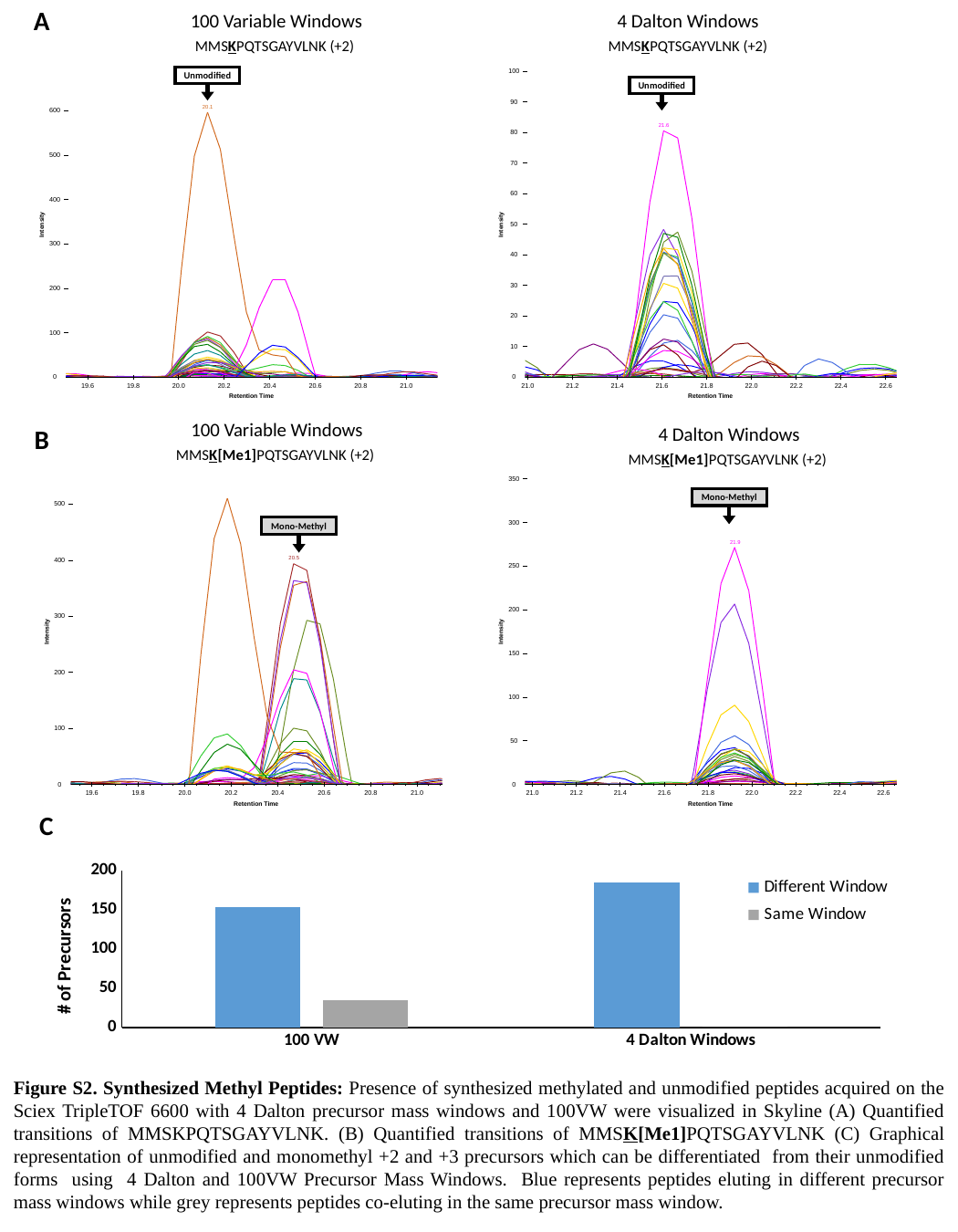

A
100 Variable Windows
MMSKPQTSGAYVLNK (+2)
4 Dalton Windows
MMSKPQTSGAYVLNK (+2)
Unmodified
Unmodified
B
100 Variable Windows
MMSK[Me1]PQTSGAYVLNK (+2)
4 Dalton Windows
MMSK[Me1]PQTSGAYVLNK (+2)
Mono-Methyl
Mono-Methyl
C
#### Chart
| Category | Different Window | Same Window |
|---|---|---|
| 100 VW | 154.0 | 35.0 |
| 4 Dalton Windows | 185.0 | 0.0 |Figure S2. Synthesized Methyl Peptides: Presence of synthesized methylated and unmodified peptides acquired on the Sciex TripleTOF 6600 with 4 Dalton precursor mass windows and 100VW were visualized in Skyline (A) Quantified transitions of MMSKPQTSGAYVLNK. (B) Quantified transitions of MMSK[Me1]PQTSGAYVLNK (C) Graphical representation of unmodified and monomethyl +2 and +3 precursors which can be differentiated from their unmodified forms using 4 Dalton and 100VW Precursor Mass Windows. Blue represents peptides eluting in different precursor mass windows while grey represents peptides co-eluting in the same precursor mass window.

### Slide 6
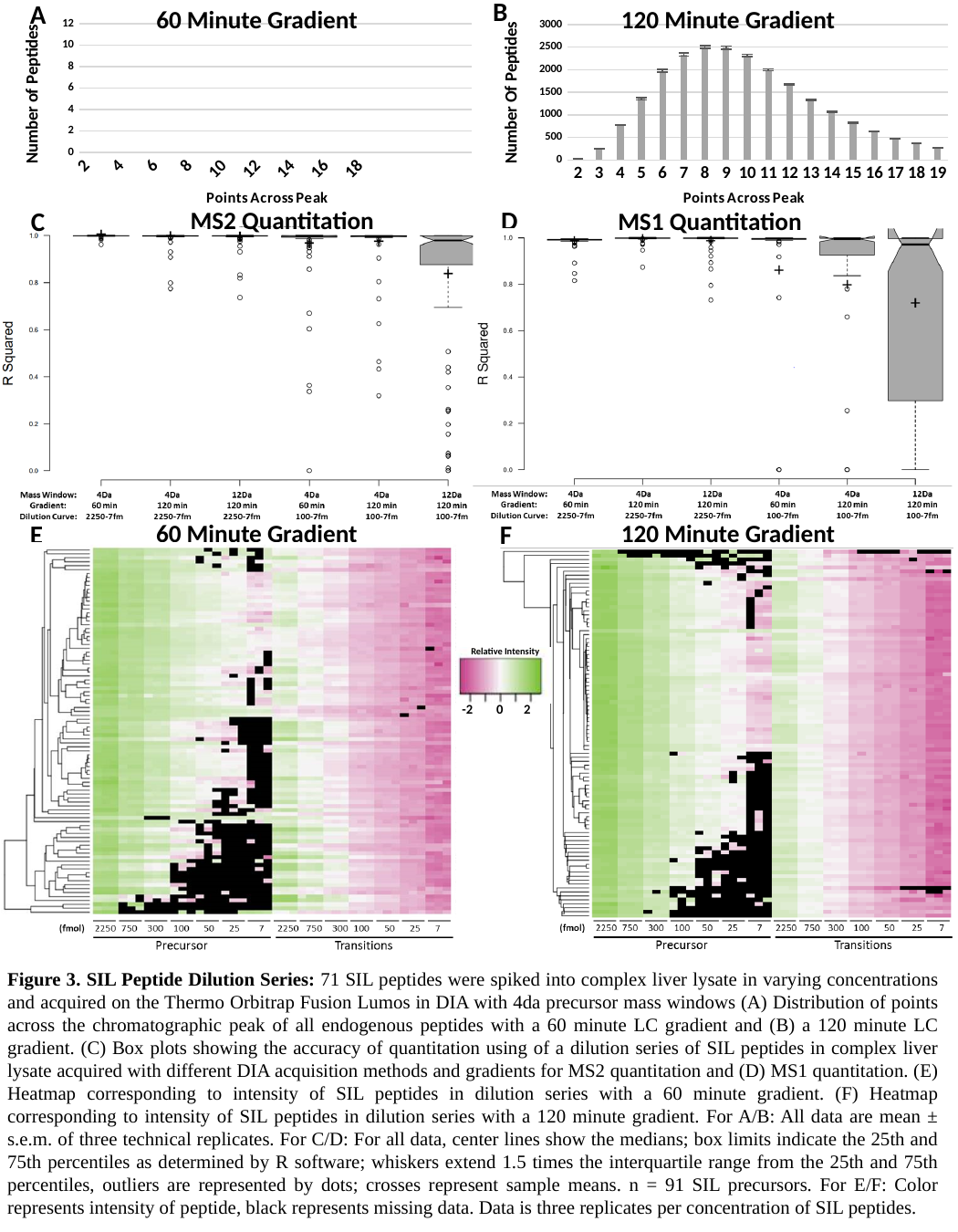

B
A
60 Minute Gradient
120 Minute Gradient
#### Chart
| Category | Points Across Peak |
|---|---|
| 2 | 8.909090909090908 |
| 3 | 246.54545454545453 |
| 4 | 1260.8181818181818 |
| 5 | 2364.7272727272725 |
| 6 | 2471.818181818182 |
| 7 | 1822.8181818181818 |
| 8 | 1276.5454545454545 |
| 9 | 841.4545454545455 |
| 10 | 542.9090909090909 |
| 11 | 334.45454545454544 |
| 12 | 209.1818181818182 |
| 13 | 133.27272727272728 |
| 14 | 89.18181818181819 |
| 15 | 61.90909090909091 |
| 16 | 42.27272727272727 |
| 17 | 32.0 |
| 18 | 21.0 |
| 19 | 17.181818181818183 |
#### Chart
| Category | Points Across Peak |
|---|---|
| 2 | 21.0 |
| 3 | 252.875 |
| 4 | 776.375 |
| 5 | 1358.5 |
| 6 | 1976.75 |
| 7 | 2339.0 |
| 8 | 2503.875 |
| 9 | 2489.875 |
| 10 | 2315.375 |
| 11 | 1996.125 |
| 12 | 1668.625 |
| 13 | 1338.375 |
| 14 | 1065.625 |
| 15 | 824.25 |
| 16 | 634.25 |
| 17 | 473.25 |
| 18 | 371.375 |
| 19 | 263.125 |D
MS2 Quantitation
C
MS1 Quantitation
60 Minute Gradient
120 Minute Gradient
F
E
Relative Intensity
-2 0 2
Figure 3. SIL Peptide Dilution Series: 71 SIL peptides were spiked into complex liver lysate in varying concentrations and acquired on the Thermo Orbitrap Fusion Lumos in DIA with 4da precursor mass windows (A) Distribution of points across the chromatographic peak of all endogenous peptides with a 60 minute LC gradient and (B) a 120 minute LC gradient. (C) Box plots showing the accuracy of quantitation using of a dilution series of SIL peptides in complex liver lysate acquired with different DIA acquisition methods and gradients for MS2 quantitation and (D) MS1 quantitation. (E) Heatmap corresponding to intensity of SIL peptides in dilution series with a 60 minute gradient. (F) Heatmap corresponding to intensity of SIL peptides in dilution series with a 120 minute gradient. For A/B: All data are mean ± s.e.m. of three technical replicates. For C/D: For all data, center lines show the medians; box limits indicate the 25th and 75th percentiles as determined by R software; whiskers extend 1.5 times the interquartile range from the 25th and 75th percentiles, outliers are represented by dots; crosses represent sample means. n = 91 SIL precursors. For E/F: Color represents intensity of peptide, black represents missing data. Data is three replicates per concentration of SIL peptides.

### Slide 7
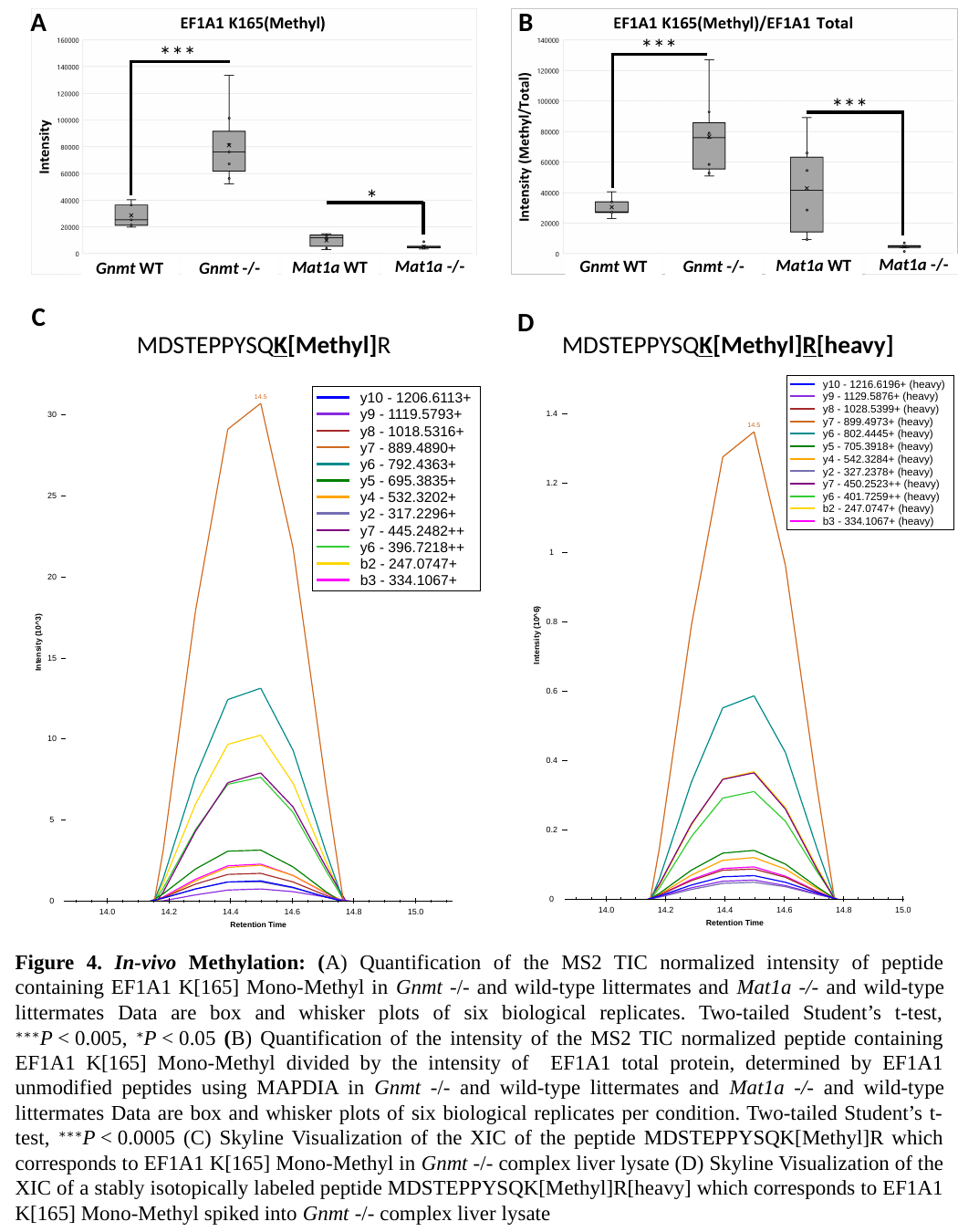

A
B
***
***
 ***
 *
Mat1a -/-
Mat1a WT
Gnmt WT
Gnmt -/-
Mat1a -/-
Mat1a WT
Gnmt WT
Gnmt -/-
C
D
MDSTEPPYSQK[Methyl]R
MDSTEPPYSQK[Methyl]R[heavy]
Figure 4. In-vivo Methylation: (A) Quantification of the MS2 TIC normalized intensity of peptide containing EF1A1 K[165] Mono-Methyl in Gnmt -/- and wild-type littermates and Mat1a -/- and wild-type littermates Data are box and whisker plots of six biological replicates. Two-tailed Student’s t-test, ^∗∗∗^P < 0.005, ^∗^P < 0.05 (B) Quantification of the intensity of the MS2 TIC normalized peptide containing EF1A1 K[165] Mono-Methyl divided by the intensity of EF1A1 total protein, determined by EF1A1 unmodified peptides using MAPDIA in Gnmt -/- and wild-type littermates and Mat1a -/- and wild-type littermates Data are box and whisker plots of six biological replicates per condition. Two-tailed Student’s t-test, ^∗∗∗^P < 0.0005 (C) Skyline Visualization of the XIC of the peptide MDSTEPPYSQK[Methyl]R which corresponds to EF1A1 K[165] Mono-Methyl in Gnmt -/- complex liver lysate (D) Skyline Visualization of the XIC of a stably isotopically labeled peptide MDSTEPPYSQK[Methyl]R[heavy] which corresponds to EF1A1 K[165] Mono-Methyl spiked into Gnmt -/- complex liver lysate

### Slide 8
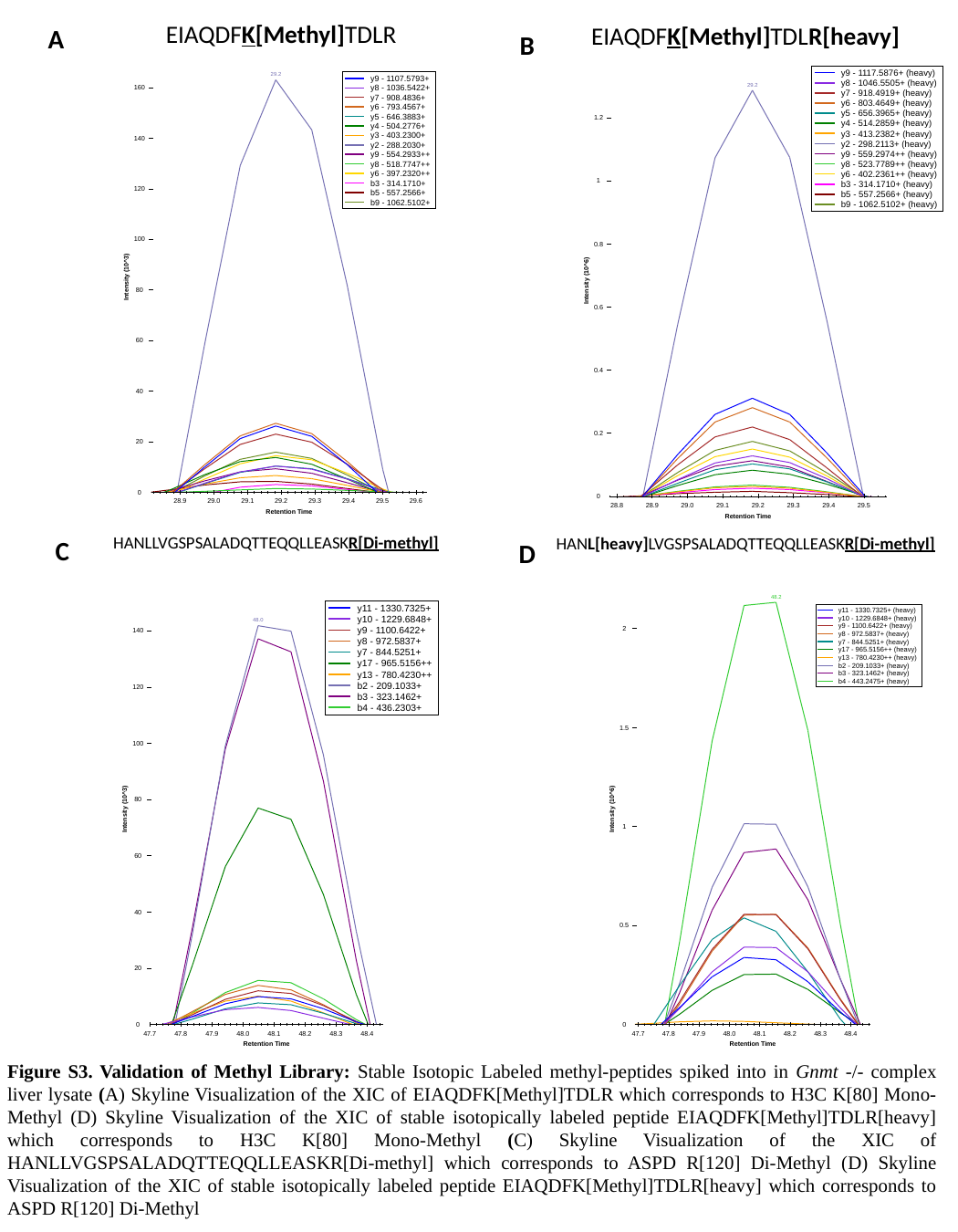

EIAQDFK[Methyl]TDLR
EIAQDFK[Methyl]TDLR[heavy]
A
B
HANLLVGSPSALADQTTEQQLLEASKR[Di-methyl]
HANL[heavy]LVGSPSALADQTTEQQLLEASKR[Di-methyl]
C
D
Figure S3. Validation of Methyl Library: Stable Isotopic Labeled methyl-peptides spiked into in Gnmt -/- complex liver lysate (A) Skyline Visualization of the XIC of EIAQDFK[Methyl]TDLR which corresponds to H3C K[80] Mono-Methyl (D) Skyline Visualization of the XIC of stable isotopically labeled peptide EIAQDFK[Methyl]TDLR[heavy] which corresponds to H3C K[80] Mono-Methyl (C) Skyline Visualization of the XIC of HANLLVGSPSALADQTTEQQLLEASKR[Di-methyl] which corresponds to ASPD R[120] Di-Methyl (D) Skyline Visualization of the XIC of stable isotopically labeled peptide EIAQDFK[Methyl]TDLR[heavy] which corresponds to ASPD R[120] Di-Methyl

### Slide 9
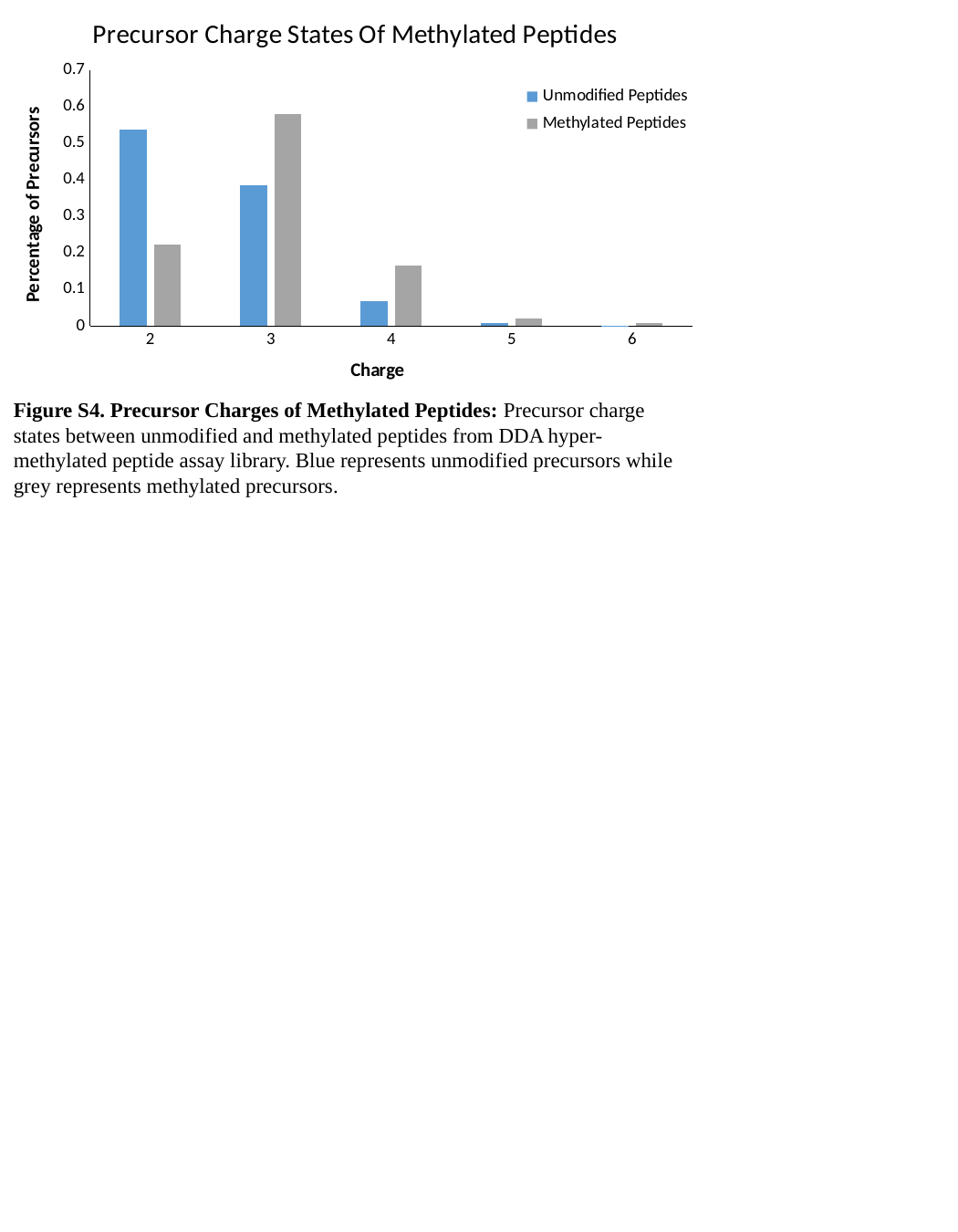

#### Chart: Precursor Charge States Of Methylated Peptides
| Category | Unmodified Peptides | Methylated Peptides |
|---|---|---|
| 2 | 0.5379916741236431 | 0.22193877551020408 |
| 3 | 0.3851106874683889 | 0.58078231292517 |
| 4 | 0.06898027467610784 | 0.16666666666666666 |
| 5 | 0.007061432517604949 | 0.021258503401360544 |
| 6 | 0.0008559312142551453 | 0.00935374149659864 |Figure S4. Precursor Charges of Methylated Peptides: Precursor charge states between unmodified and methylated peptides from DDA hyper-methylated peptide assay library. Blue represents unmodified precursors while grey represents methylated precursors.

### Slide 10
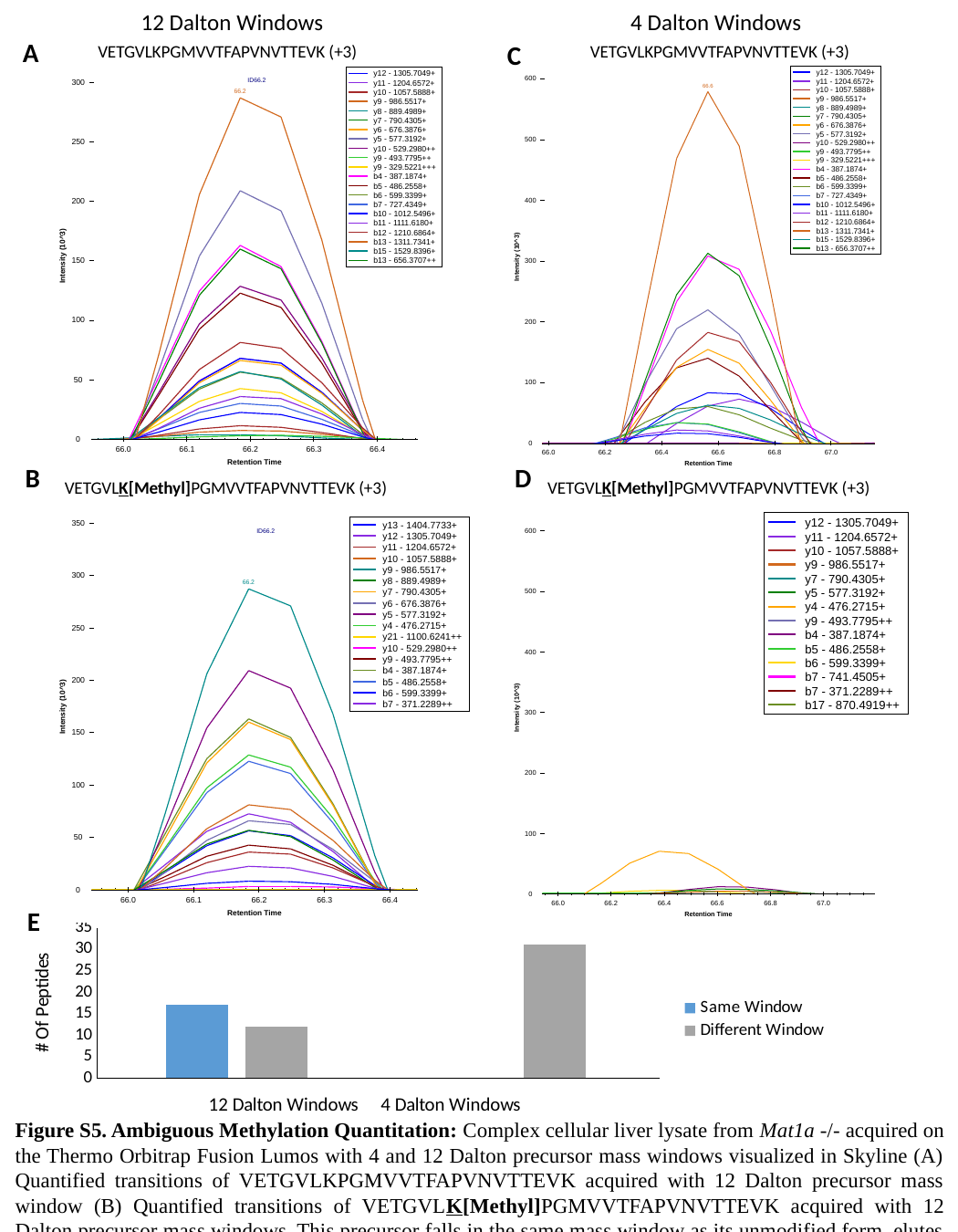

12 Dalton Windows
4 Dalton Windows
A
C
VETGVLKPGMVVTFAPVNVTTEVK (+3)
VETGVLKPGMVVTFAPVNVTTEVK (+3)
B
D
VETGVLK[Methyl]PGMVVTFAPVNVTTEVK (+3)
VETGVLK[Methyl]PGMVVTFAPVNVTTEVK (+3)
E
#### Chart
| Category | Same Window | Different Window |
|---|---|---|
| False Positives 12 Dalton Windows | 17.0 | 12.0 |
| False Positives 4 Dalton Windows | 0.0 | 31.0 |Figure S5. Ambiguous Methylation Quantitation: Complex cellular liver lysate from Mat1a -/- acquired on the Thermo Orbitrap Fusion Lumos with 4 and 12 Dalton precursor mass windows visualized in Skyline (A) Quantified transitions of VETGVLKPGMVVTFAPVNVTTEVK acquired with 12 Dalton precursor mass window (B) Quantified transitions of VETGVLK[Methyl]PGMVVTFAPVNVTTEVK acquired with 12 Dalton precursor mass windows. This precursor falls in the same mass window as its unmodified form, elutes at the same time, and contains one site specific transition (b7). (C) Quantified transitions of VETGVLKPGMVVTFAPVNVTTEVK acquired with 4 Dalton precursor mass window (D) Quantified transitions of VETGVLK[Methyl]PGMVVTFAPVNVTTEVK acquired with 4 Dalton precursor mass windows. There is no discernable peak; leading us to believe that the quantitation of the methylated peptide found with 12 Dalton precursor mass windows is incorrect due to many ambiguous transitions.(E) Graphical representation of unmodified, monomethyl, and dimethyl +2 and +3 precursors which can be differentiated from their unmodified forms using 4 and 12 Dalton precursor mass windows. Grey represents peptides eluting in different precursor mass windows while blue represents peptides co-eluting in the same precursor mass window.

### Slide 11
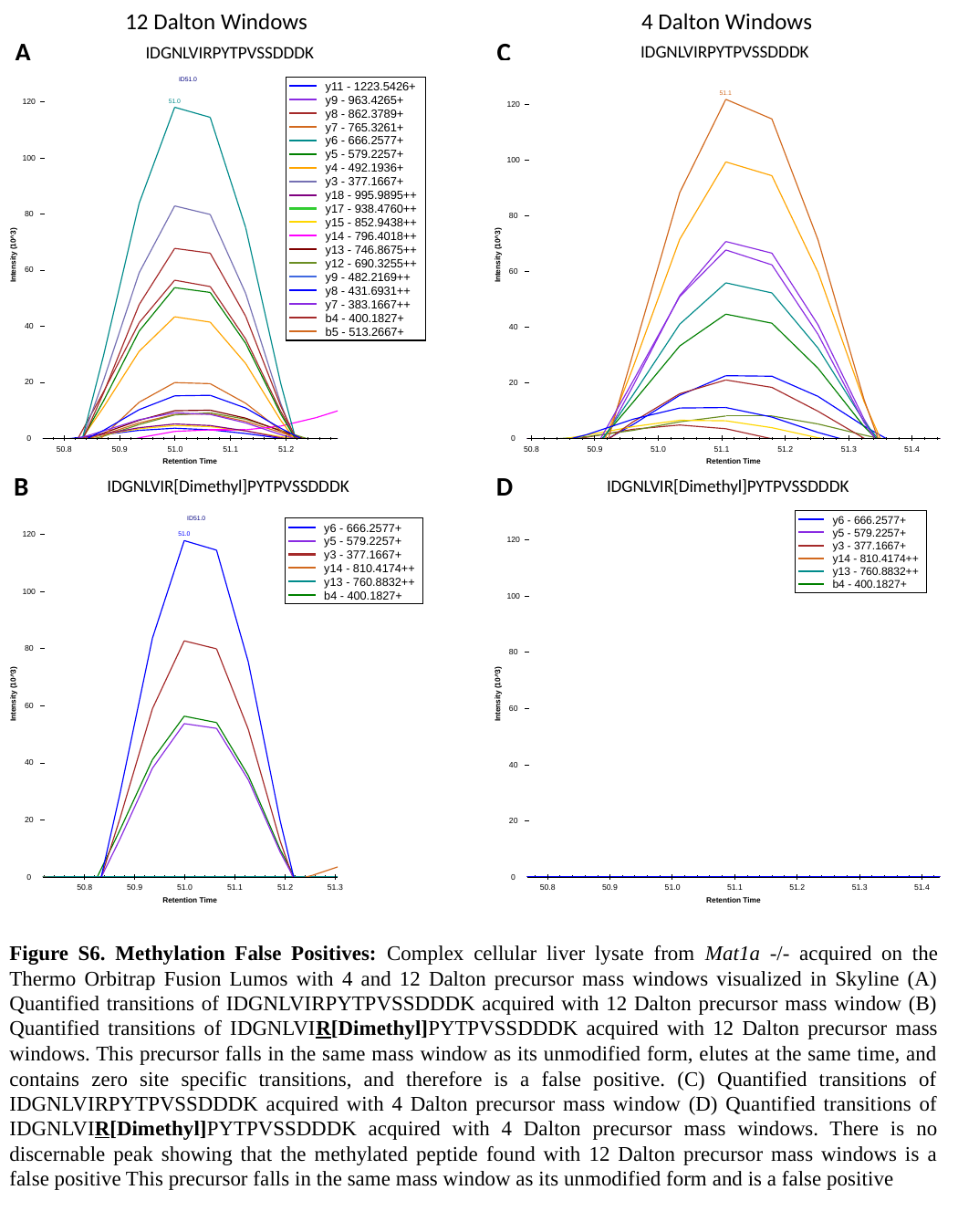

4 Dalton Windows
12 Dalton Windows
A
C
IDGNLVIRPYTPVSSDDDK
IDGNLVIRPYTPVSSDDDK
B
D
IDGNLVIR[Dimethyl]PYTPVSSDDDK
IDGNLVIR[Dimethyl]PYTPVSSDDDK
Figure S6. Methylation False Positives: Complex cellular liver lysate from Mat1a -/- acquired on the Thermo Orbitrap Fusion Lumos with 4 and 12 Dalton precursor mass windows visualized in Skyline (A) Quantified transitions of IDGNLVIRPYTPVSSDDDK acquired with 12 Dalton precursor mass window (B) Quantified transitions of IDGNLVIR[Dimethyl]PYTPVSSDDDK acquired with 12 Dalton precursor mass windows. This precursor falls in the same mass window as its unmodified form, elutes at the same time, and contains zero site specific transitions, and therefore is a false positive. (C) Quantified transitions of IDGNLVIRPYTPVSSDDDK acquired with 4 Dalton precursor mass window (D) Quantified transitions of IDGNLVIR[Dimethyl]PYTPVSSDDDK acquired with 4 Dalton precursor mass windows. There is no discernable peak showing that the methylated peptide found with 12 Dalton precursor mass windows is a false positive This precursor falls in the same mass window as its unmodified form and is a false positive

### Slide 12
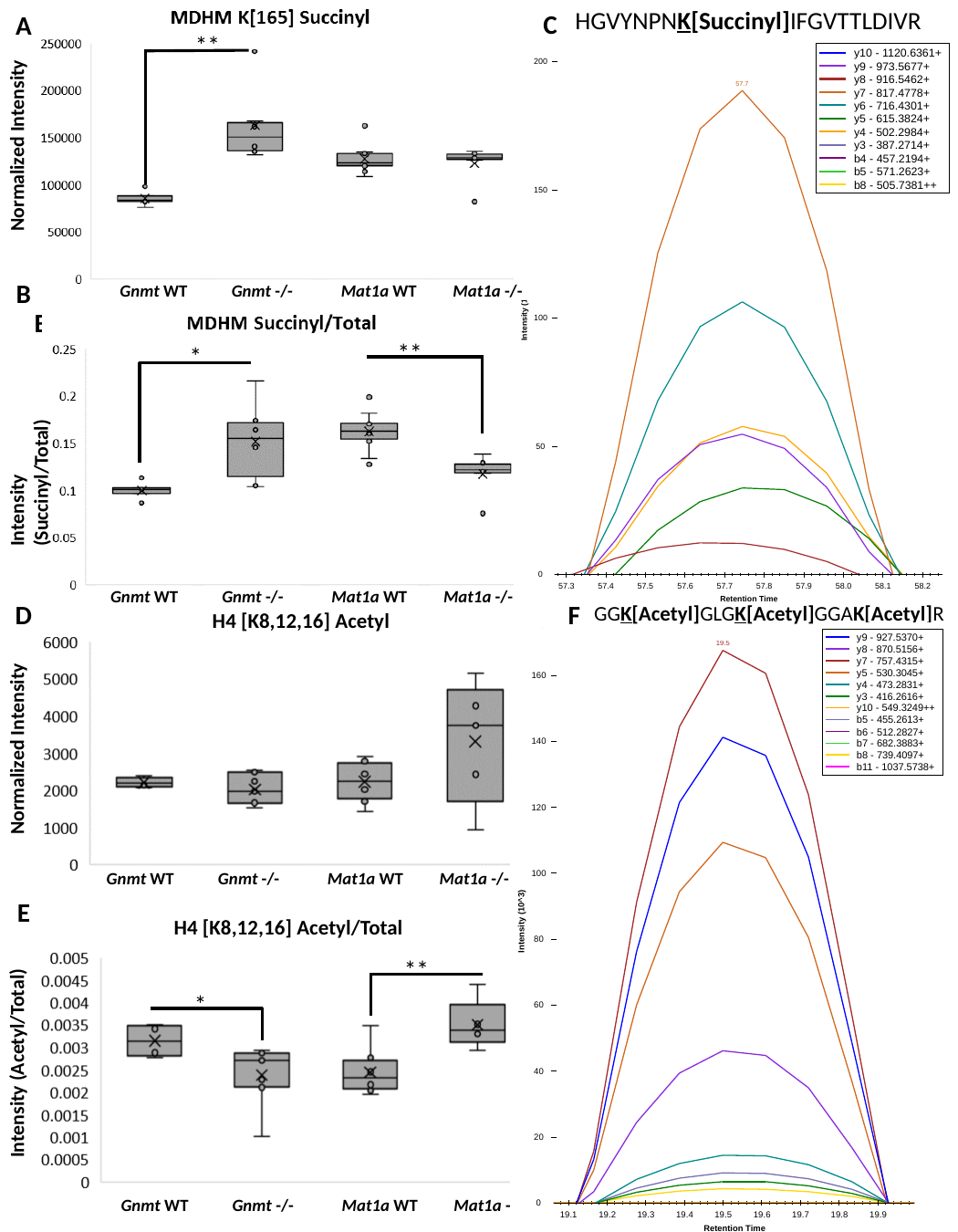

HGVYNPNK[Succinyl]IFGVTTLDIVR
C
A
Normalized Intensity
**
B
Gnmt WT
Gnmt -/-
Mat1a WT
Mat1a -/-
B
**
Intensity (Succinyl/Total)
*
Gnmt WT
Gnmt -/-
Mat1a WT
Mat1a -/-
D
F
GGK[Acetyl]GLGK[Acetyl]GGAK[Acetyl]R
H4 [K8,12,16] Acetyl
Normalized Intensity
Gnmt WT
Gnmt -/-
Mat1a WT
Mat1a -/-
E
H4 [K8,12,16] Acetyl/Total
Intensity (Acetyl/Total)
**
 *
Gnmt WT
Gnmt -/-
Mat1a WT
Mat1a -/-
Figure 5. In-vivo Succinylation and Acetylation: (A) Quantification of the MS2 TIC normalized intensity of peptide containing MDHM K[165] Succinyl in Gnmt -/- and wild-type littermates and Mat1a -/- and wild-type littermates (n=6/condition) Data are box and whisker plots of six biological replicates per condition. Two-tailed Student’s t-test, ^∗∗^P < 0.005 (B) Quantification of the intensity of the MS2 TIC normalized peptide containing MDHM K[165] Succinyl divided by the intensity of MDHM total protein, determined by MDHM unmodified peptides using MAPDIA. Data are box and whisker plots of six biological replicates per condition. Two-tailed Student’s t-test, ^∗∗^P < 0.005, ^∗^P < 0.05 (C) Skyline Visualization of the XIC of the peptide containing MDHM K[165] Succinyl (D) Quantification of the MS2 TIC normalized intensity of a proteotypic peptide mapping to H4 K[8,12,16] Acetyl in Gnmt -/- and wild-type littermates and Mat1a -/- and wild-type littermates. Data are box and whisker plots of six biological replicates per condition. (E) Quantification of the intensity of the MS2 TIC normalized peptide containing H4 K[8,12,16] Acetyl divided by the intensity of H4 total protein, determined by H4 unmodified peptides using MAPDIA in Gnmt -/- and wild-type littermates and Mat1a -/- and wild-type littermates. Data are box and whisker plots of six biological replicates per condition. Two-tailed Student’s t-test, ^∗∗^P < 0.005, ^∗^P < 0.05 (F) Skyline Visualization of the XIC of the peptide containing H4 K[8,12,16] Acetyl

### Slide 13
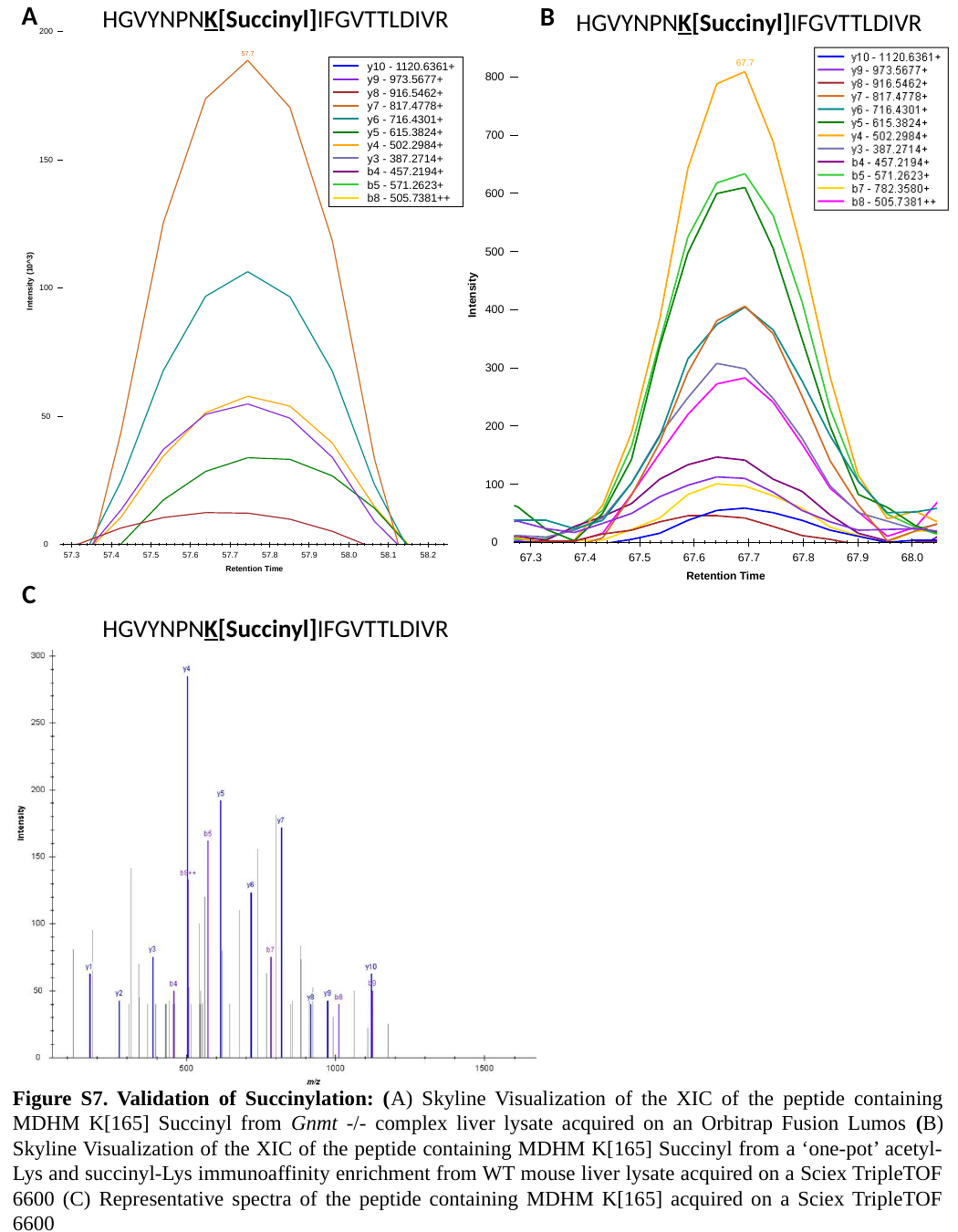

HGVYNPNK[Succinyl]IFGVTTLDIVR
A
B
HGVYNPNK[Succinyl]IFGVTTLDIVR
C
HGVYNPNK[Succinyl]IFGVTTLDIVR
Figure S7. Validation of Succinylation: (A) Skyline Visualization of the XIC of the peptide containing MDHM K[165] Succinyl from Gnmt -/- complex liver lysate acquired on an Orbitrap Fusion Lumos (B) Skyline Visualization of the XIC of the peptide containing MDHM K[165] Succinyl from a ‘one-pot’ acetyl-Lys and succinyl-Lys immunoaffinity enrichment from WT mouse liver lysate acquired on a Sciex TripleTOF 6600 (C) Representative spectra of the peptide containing MDHM K[165] acquired on a Sciex TripleTOF 6600

### Slide 14
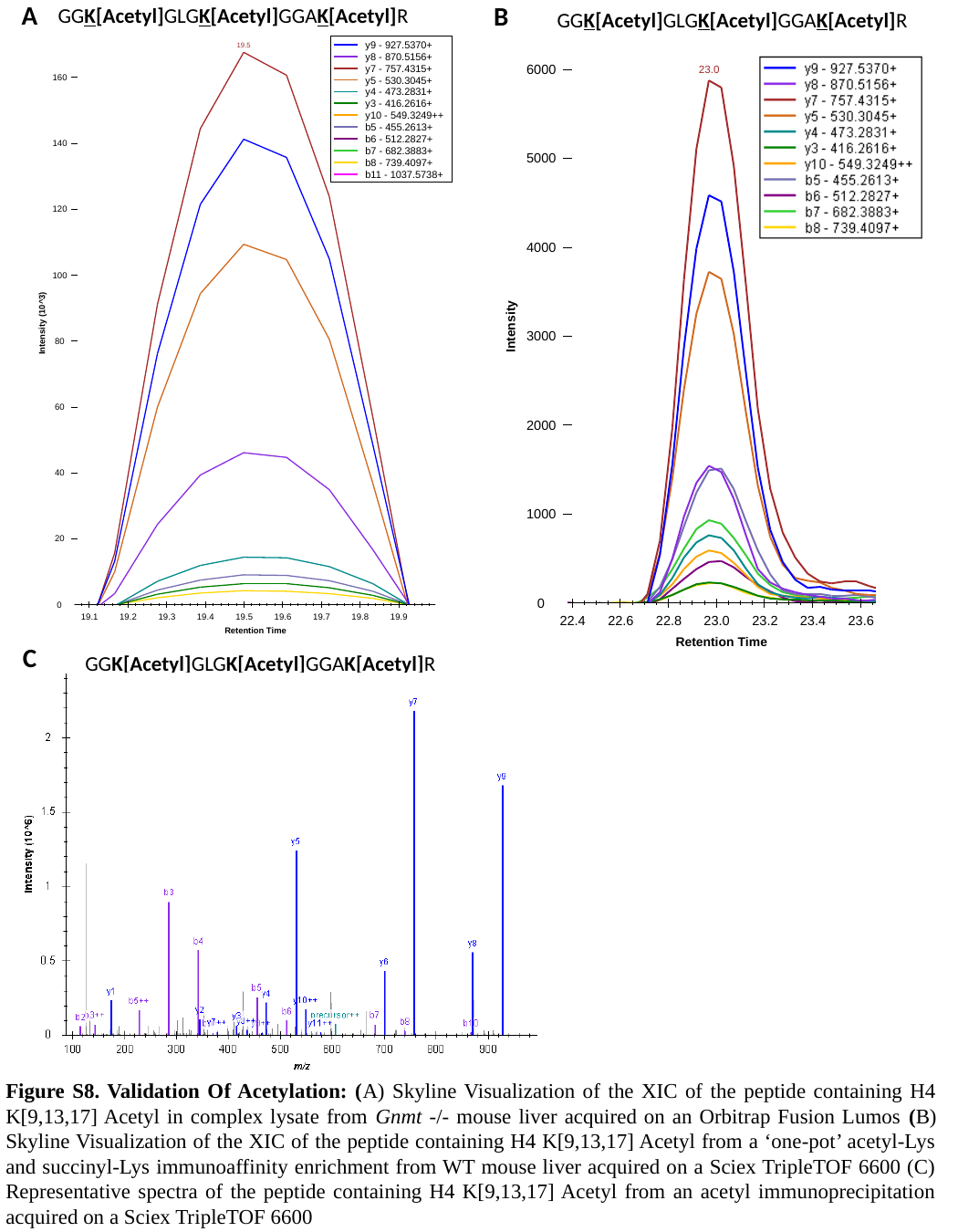

GGK[Acetyl]GLGK[Acetyl]GGAK[Acetyl]R
A
B
GGK[Acetyl]GLGK[Acetyl]GGAK[Acetyl]R
C
GGK[Acetyl]GLGK[Acetyl]GGAK[Acetyl]R
Figure S8. Validation Of Acetylation: (A) Skyline Visualization of the XIC of the peptide containing H4 K[9,13,17] Acetyl in complex lysate from Gnmt -/- mouse liver acquired on an Orbitrap Fusion Lumos (B) Skyline Visualization of the XIC of the peptide containing H4 K[9,13,17] Acetyl from a ‘one-pot’ acetyl-Lys and succinyl-Lys immunoaffinity enrichment from WT mouse liver acquired on a Sciex TripleTOF 6600 (C) Representative spectra of the peptide containing H4 K[9,13,17] Acetyl from an acetyl immunoprecipitation acquired on a Sciex TripleTOF 6600

### Slide 15
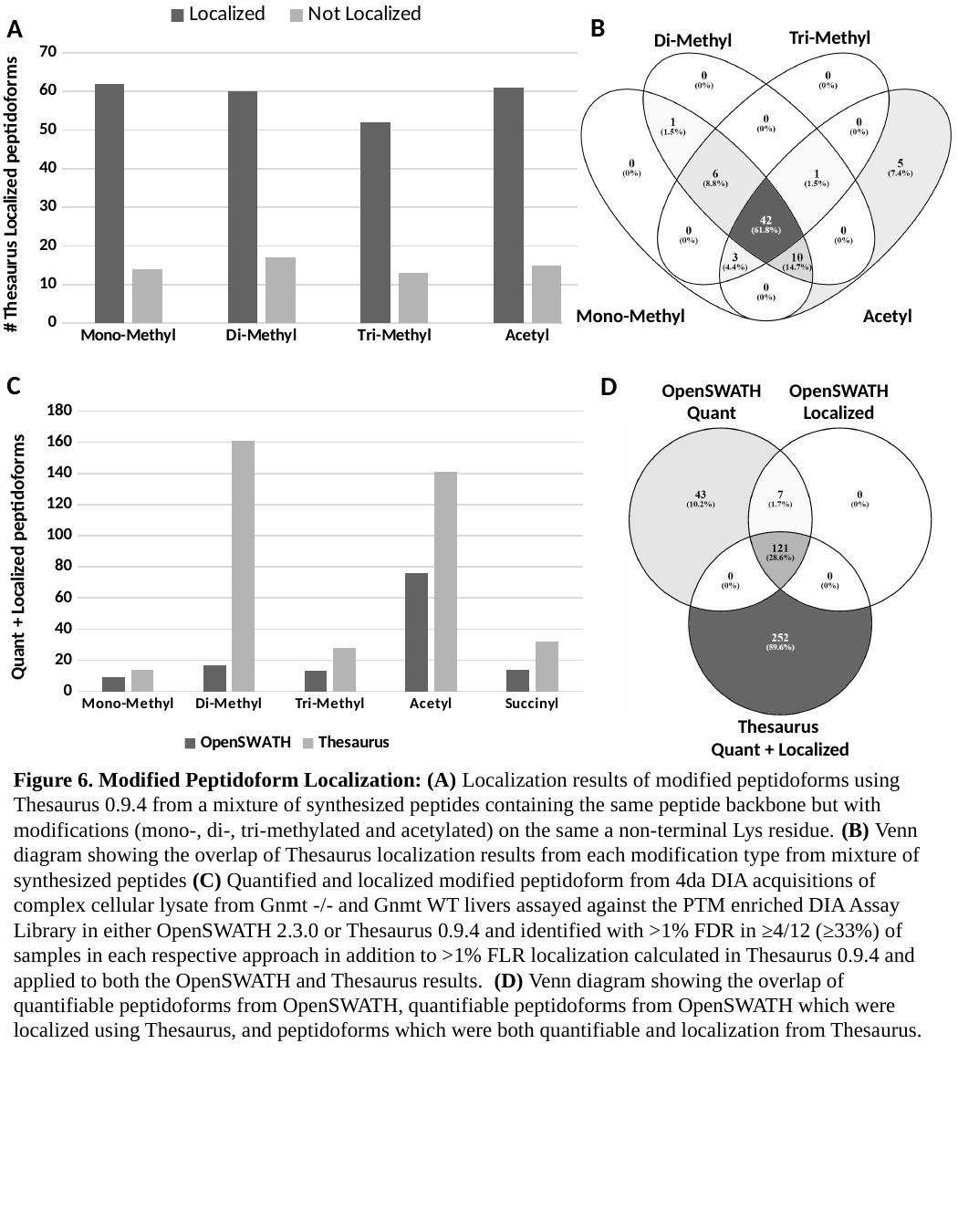

#### Chart
| Category | Localized | Not Localized |
|---|---|---|
| Mono-Methyl | 62.0 | 14.0 |
| Di-Methyl | 60.0 | 17.0 |
| Tri-Methyl | 52.0 | 13.0 |
| Acetyl | 61.0 | 15.0 |B
A
Tri-Methyl
Di-Methyl
Mono-Methyl
Acetyl
C
D
OpenSWATH Quant
OpenSWATH
Localized
#### Chart
| Category | OpenSWATH | Thesaurus |
|---|---|---|
| Mono-Methyl | 9.0 | 14.0 |
| Di-Methyl | 17.0 | 161.0 |
| Tri-Methyl | 13.0 | 28.0 |
| Acetyl | 76.0 | 141.0 |
| Succinyl | 14.0 | 32.0 |
Thesaurus
 Quant + Localized
Figure 6. Modified Peptidoform Localization: (A) Localization results of modified peptidoforms using Thesaurus 0.9.4 from a mixture of synthesized peptides containing the same peptide backbone but with modifications (mono-, di-, tri-methylated and acetylated) on the same a non-terminal Lys residue. (B) Venn diagram showing the overlap of Thesaurus localization results from each modification type from mixture of synthesized peptides (C) Quantified and localized modified peptidoform from 4da DIA acquisitions of complex cellular lysate from Gnmt -/- and Gnmt WT livers assayed against the PTM enriched DIA Assay Library in either OpenSWATH 2.3.0 or Thesaurus 0.9.4 and identified with >1% FDR in ≥4/12 (≥33%) of samples in each respective approach in addition to >1% FLR localization calculated in Thesaurus 0.9.4 and applied to both the OpenSWATH and Thesaurus results. (D) Venn diagram showing the overlap of quantifiable peptidoforms from OpenSWATH, quantifiable peptidoforms from OpenSWATH which were localized using Thesaurus, and peptidoforms which were both quantifiable and localization from Thesaurus.

### Slide 16
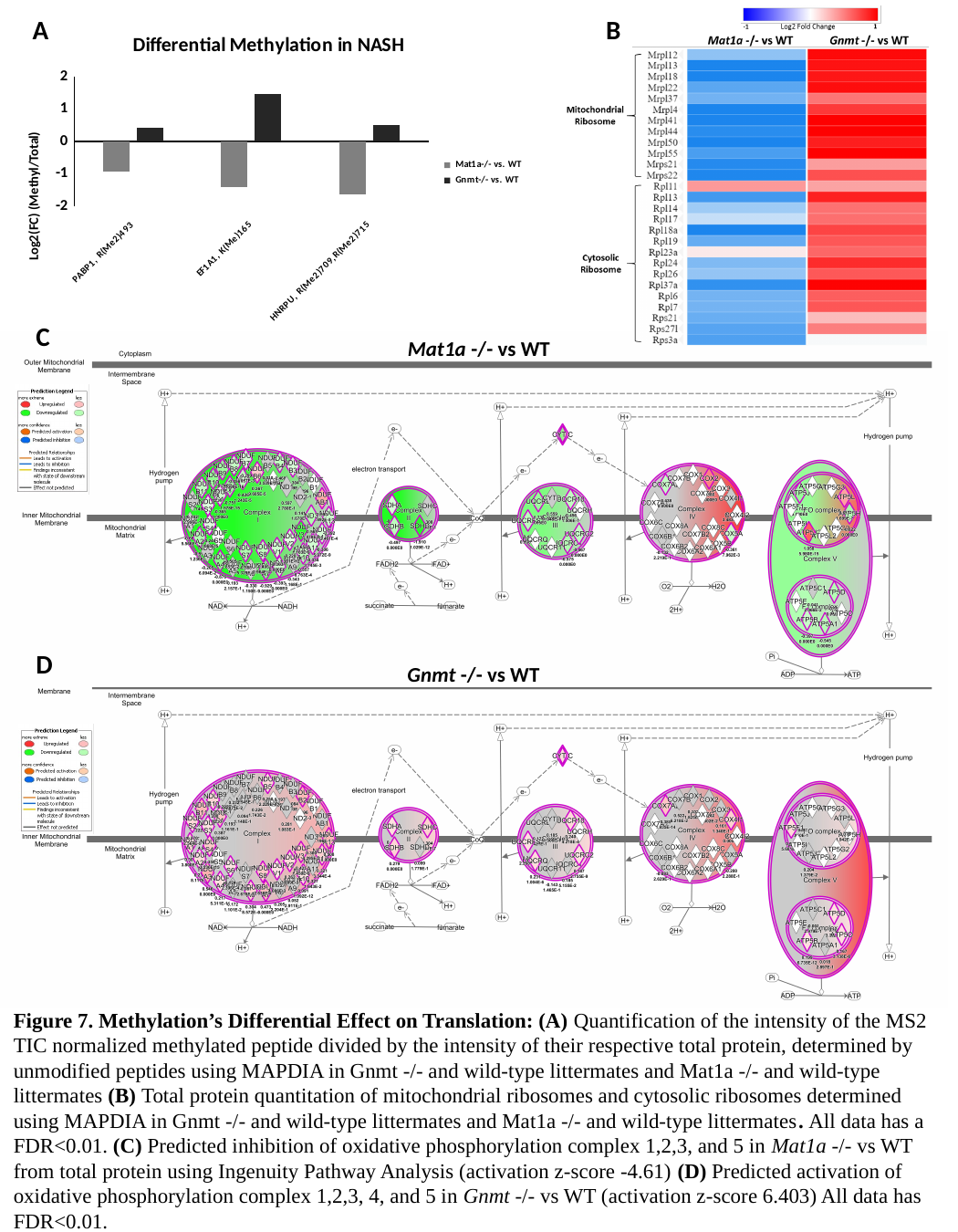

A
B
#### Chart: Differential Methylation in NASH
| Category | Mat1a-/- vs. WT | Gnmt-/- vs. WT |
|---|---|---|
| PABP1, R(Me2)493 | -0.9260987478263825 | 0.4282536067737104 |
| EF1A1, K(Me)165 | -1.397770491508122 | 1.4715467937839652 |
| HNRPU, R(Me2)709,R(Me2)715 | -1.6204002403315947 | 0.4975153792742217 |C
Mat1a -/- vs WT
D
Gnmt -/- vs WT
Figure 7. Methylation’s Differential Effect on Translation: (A) Quantification of the intensity of the MS2 TIC normalized methylated peptide divided by the intensity of their respective total protein, determined by unmodified peptides using MAPDIA in Gnmt -/- and wild-type littermates and Mat1a -/- and wild-type littermates (B) Total protein quantitation of mitochondrial ribosomes and cytosolic ribosomes determined using MAPDIA in Gnmt -/- and wild-type littermates and Mat1a -/- and wild-type littermates. All data has a FDR<0.01. (C) Predicted inhibition of oxidative phosphorylation complex 1,2,3, and 5 in Mat1a -/- vs WT from total protein using Ingenuity Pathway Analysis (activation z-score -4.61) (D) Predicted activation of oxidative phosphorylation complex 1,2,3, 4, and 5 in Gnmt -/- vs WT (activation z-score 6.403) All data has FDR<0.01.

### Slide 17
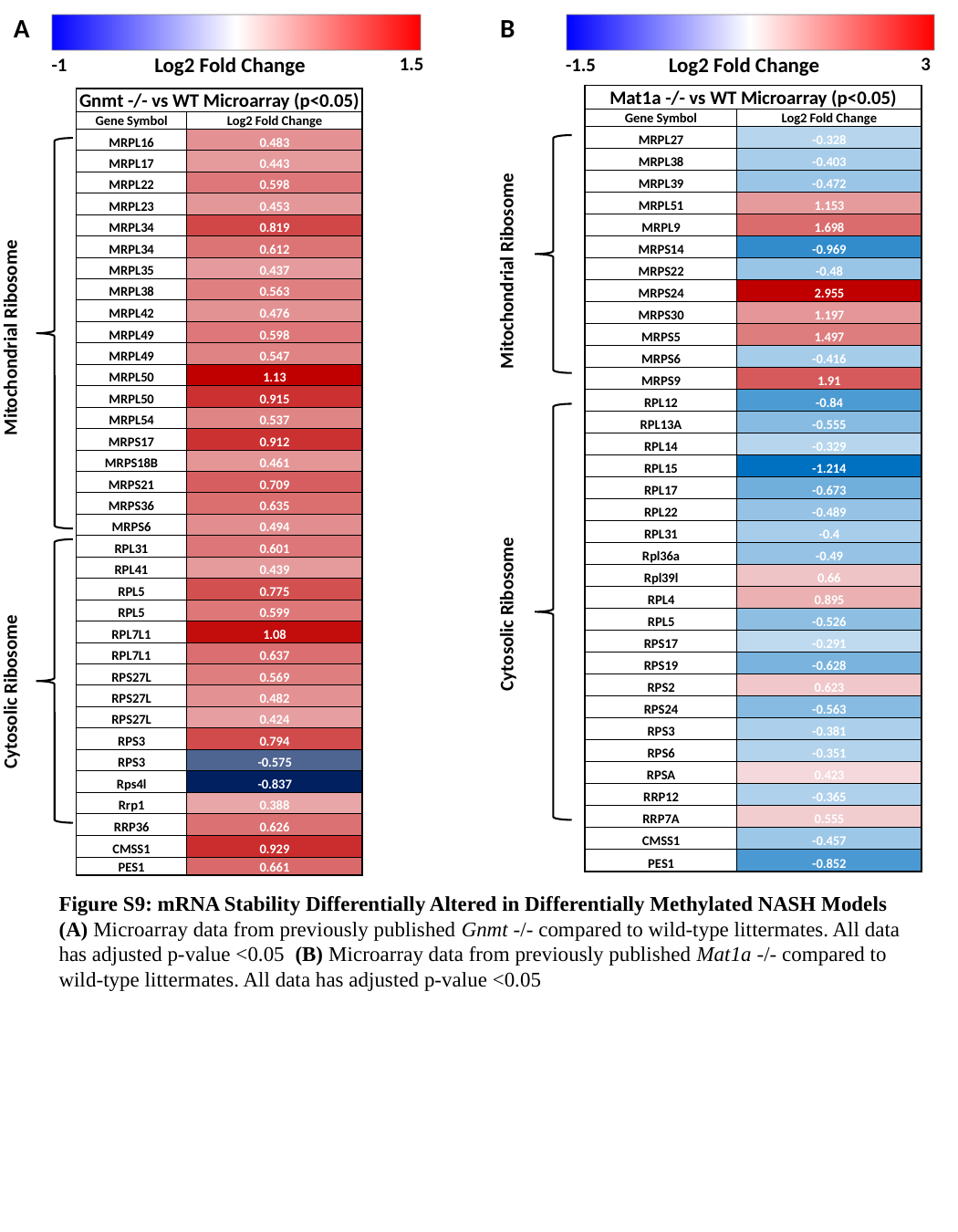

A
B
Log2 Fold Change
1.5
-1
Log2 Fold Change
3
-1.5
| Mat1a -/- vs WT Microarray (p<0.05) | |
| --- | --- |
| Gene Symbol | Log2 Fold Change |
| MRPL27 | -0.328 |
| MRPL38 | -0.403 |
| MRPL39 | -0.472 |
| MRPL51 | 1.153 |
| MRPL9 | 1.698 |
| MRPS14 | -0.969 |
| MRPS22 | -0.48 |
| MRPS24 | 2.955 |
| MRPS30 | 1.197 |
| MRPS5 | 1.497 |
| MRPS6 | -0.416 |
| MRPS9 | 1.91 |
| RPL12 | -0.84 |
| RPL13A | -0.555 |
| RPL14 | -0.329 |
| RPL15 | -1.214 |
| RPL17 | -0.673 |
| RPL22 | -0.489 |
| RPL31 | -0.4 |
| Rpl36a | -0.49 |
| Rpl39l | 0.66 |
| RPL4 | 0.895 |
| RPL5 | -0.526 |
| RPS17 | -0.291 |
| RPS19 | -0.628 |
| RPS2 | 0.623 |
| RPS24 | -0.563 |
| RPS3 | -0.381 |
| RPS6 | -0.351 |
| RPSA | 0.423 |
| RRP12 | -0.365 |
| RRP7A | 0.555 |
| CMSS1 | -0.457 |
| PES1 | -0.852 |
| Gnmt -/- vs WT Microarray (p<0.05) | |
| --- | --- |
| Gene Symbol | Log2 Fold Change |
| MRPL16 | 0.483 |
| MRPL17 | 0.443 |
| MRPL22 | 0.598 |
| MRPL23 | 0.453 |
| MRPL34 | 0.819 |
| MRPL34 | 0.612 |
| MRPL35 | 0.437 |
| MRPL38 | 0.563 |
| MRPL42 | 0.476 |
| MRPL49 | 0.598 |
| MRPL49 | 0.547 |
| MRPL50 | 1.13 |
| MRPL50 | 0.915 |
| MRPL54 | 0.537 |
| MRPS17 | 0.912 |
| MRPS18B | 0.461 |
| MRPS21 | 0.709 |
| MRPS36 | 0.635 |
| MRPS6 | 0.494 |
| RPL31 | 0.601 |
| RPL41 | 0.439 |
| RPL5 | 0.775 |
| RPL5 | 0.599 |
| RPL7L1 | 1.08 |
| RPL7L1 | 0.637 |
| RPS27L | 0.569 |
| RPS27L | 0.482 |
| RPS27L | 0.424 |
| RPS3 | 0.794 |
| RPS3 | -0.575 |
| Rps4l | -0.837 |
| Rrp1 | 0.388 |
| RRP36 | 0.626 |
| CMSS1 | 0.929 |
| PES1 | 0.661 |
Mitochondrial Ribosome
Mitochondrial Ribosome
Cytosolic Ribosome
Cytosolic Ribosome
Figure S9: mRNA Stability Differentially Altered in Differentially Methylated NASH Models (A) Microarray data from previously published Gnmt -/- compared to wild-type littermates. All data has adjusted p-value <0.05 (B) Microarray data from previously published Mat1a -/- compared to wild-type littermates. All data has adjusted p-value <0.05

### Slide 18
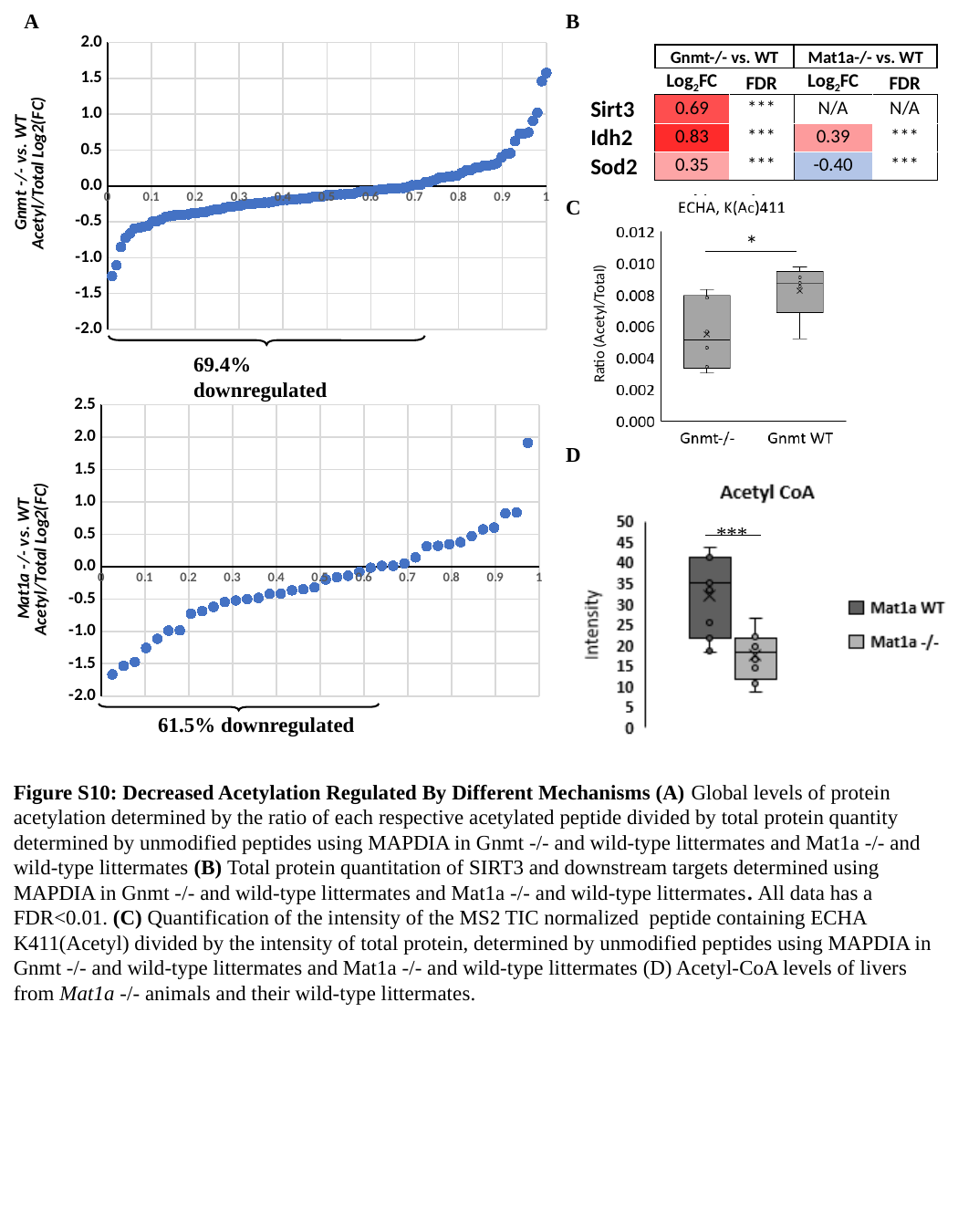

#### Chart:
| Category | |
|---|---|A
B
| | Gnmt-/- vs. WT | | Mat1a-/- vs. WT | |
| --- | --- | --- | --- | --- |
| | Log_2_FC | FDR | Log_2_FC | FDR |
| Sirt3 | 0.69 | \*\*\* | N/A | N/A |
| Idh2 | 0.83 | \*\*\* | 0.39 | \*\*\* |
| Sod2 | 0.35 | \*\*\* | -0.40 | \*\*\* |
C
*
Ratio (Acetyl/Total)
#### Chart:
| Category | |
|---|---|69.4% downregulated
D
***
61.5% downregulated
Figure S10: Decreased Acetylation Regulated By Different Mechanisms (A) Global levels of protein acetylation determined by the ratio of each respective acetylated peptide divided by total protein quantity determined by unmodified peptides using MAPDIA in Gnmt -/- and wild-type littermates and Mat1a -/- and wild-type littermates (B) Total protein quantitation of SIRT3 and downstream targets determined using MAPDIA in Gnmt -/- and wild-type littermates and Mat1a -/- and wild-type littermates. All data has a FDR<0.01. (C) Quantification of the intensity of the MS2 TIC normalized peptide containing ECHA K411(Acetyl) divided by the intensity of total protein, determined by unmodified peptides using MAPDIA in Gnmt -/- and wild-type littermates and Mat1a -/- and wild-type littermates (D) Acetyl-CoA levels of livers from Mat1a -/- animals and their wild-type littermates.

### Slide 19
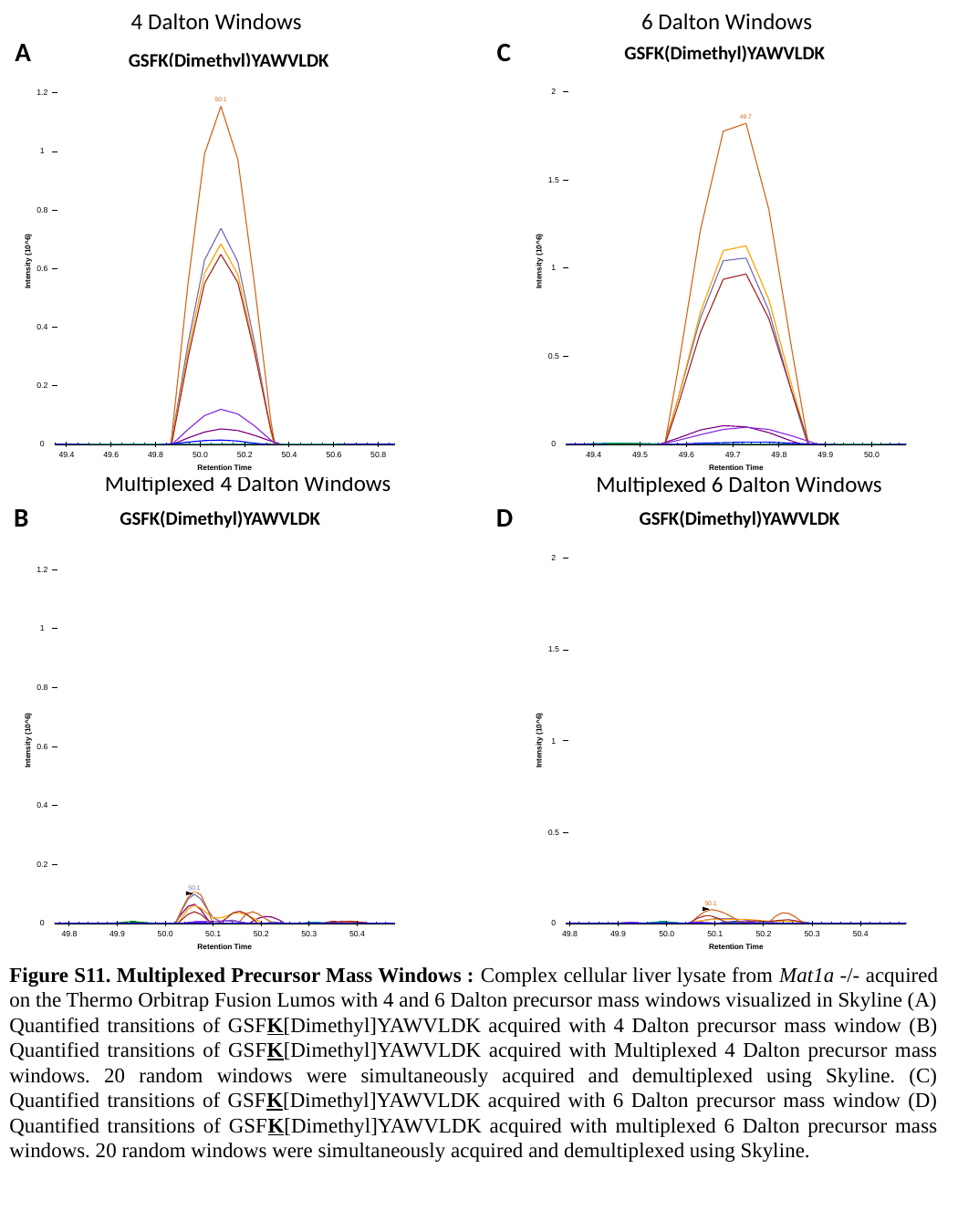

6 Dalton Windows
4 Dalton Windows
A
C
GSFK(Dimethyl)YAWVLDK
GSFK(Dimethyl)YAWVLDK
Multiplexed 4 Dalton Windows
Multiplexed 6 Dalton Windows
B
D
GSFK(Dimethyl)YAWVLDK
GSFK(Dimethyl)YAWVLDK
Figure S11. Multiplexed Precursor Mass Windows : Complex cellular liver lysate from Mat1a -/- acquired on the Thermo Orbitrap Fusion Lumos with 4 and 6 Dalton precursor mass windows visualized in Skyline (A) Quantified transitions of GSFK[Dimethyl]YAWVLDK acquired with 4 Dalton precursor mass window (B) Quantified transitions of GSFK[Dimethyl]YAWVLDK acquired with Multiplexed 4 Dalton precursor mass windows. 20 random windows were simultaneously acquired and demultiplexed using Skyline. (C) Quantified transitions of GSFK[Dimethyl]YAWVLDK acquired with 6 Dalton precursor mass window (D) Quantified transitions of GSFK[Dimethyl]YAWVLDK acquired with multiplexed 6 Dalton precursor mass windows. 20 random windows were simultaneously acquired and demultiplexed using Skyline.

### Slide 20
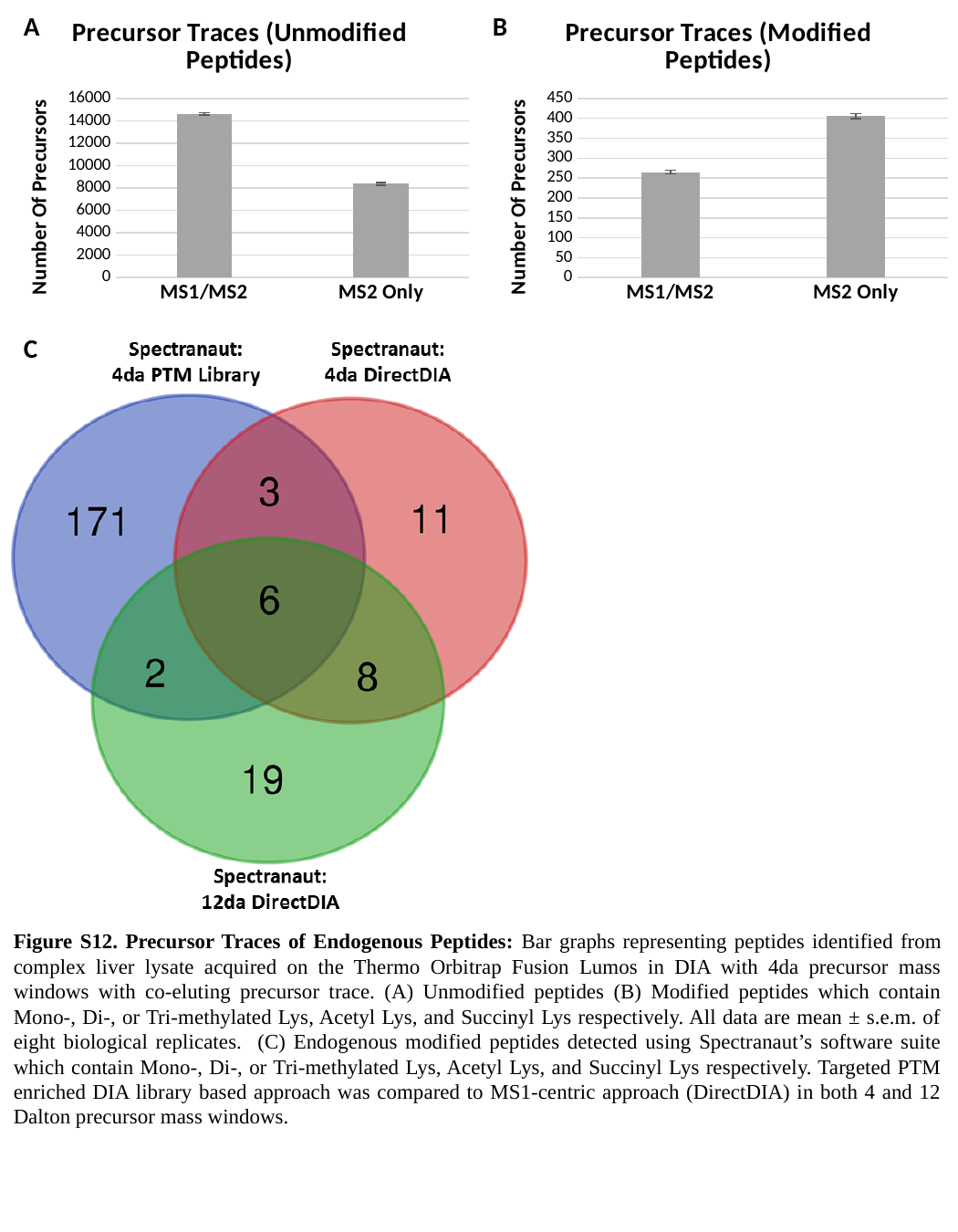

#### Chart: Precursor Traces (Unmodified Peptides)
| Category | MS1 Traces |
|---|---|
| MS1/MS2 | 14629.0 |
| MS2 Only | 8368.25 |
#### Chart: Precursor Traces (Modified Peptides)
| Category | MS1 Traces (Modified Peptides) |
|---|---|
| MS1/MS2 | 264.75 |
| MS2 Only | 405.25 |B
A
C
Figure S12. Precursor Traces of Endogenous Peptides: Bar graphs representing peptides identified from complex liver lysate acquired on the Thermo Orbitrap Fusion Lumos in DIA with 4da precursor mass windows with co-eluting precursor trace. (A) Unmodified peptides (B) Modified peptides which contain Mono-, Di-, or Tri-methylated Lys, Acetyl Lys, and Succinyl Lys respectively. All data are mean ± s.e.m. of eight biological replicates. (C) Endogenous modified peptides detected using Spectranaut’s software suite which contain Mono-, Di-, or Tri-methylated Lys, Acetyl Lys, and Succinyl Lys respectively. Targeted PTM enriched DIA library based approach was compared to MS1-centric approach (DirectDIA) in both 4 and 12 Dalton precursor mass windows.

### Slide 21
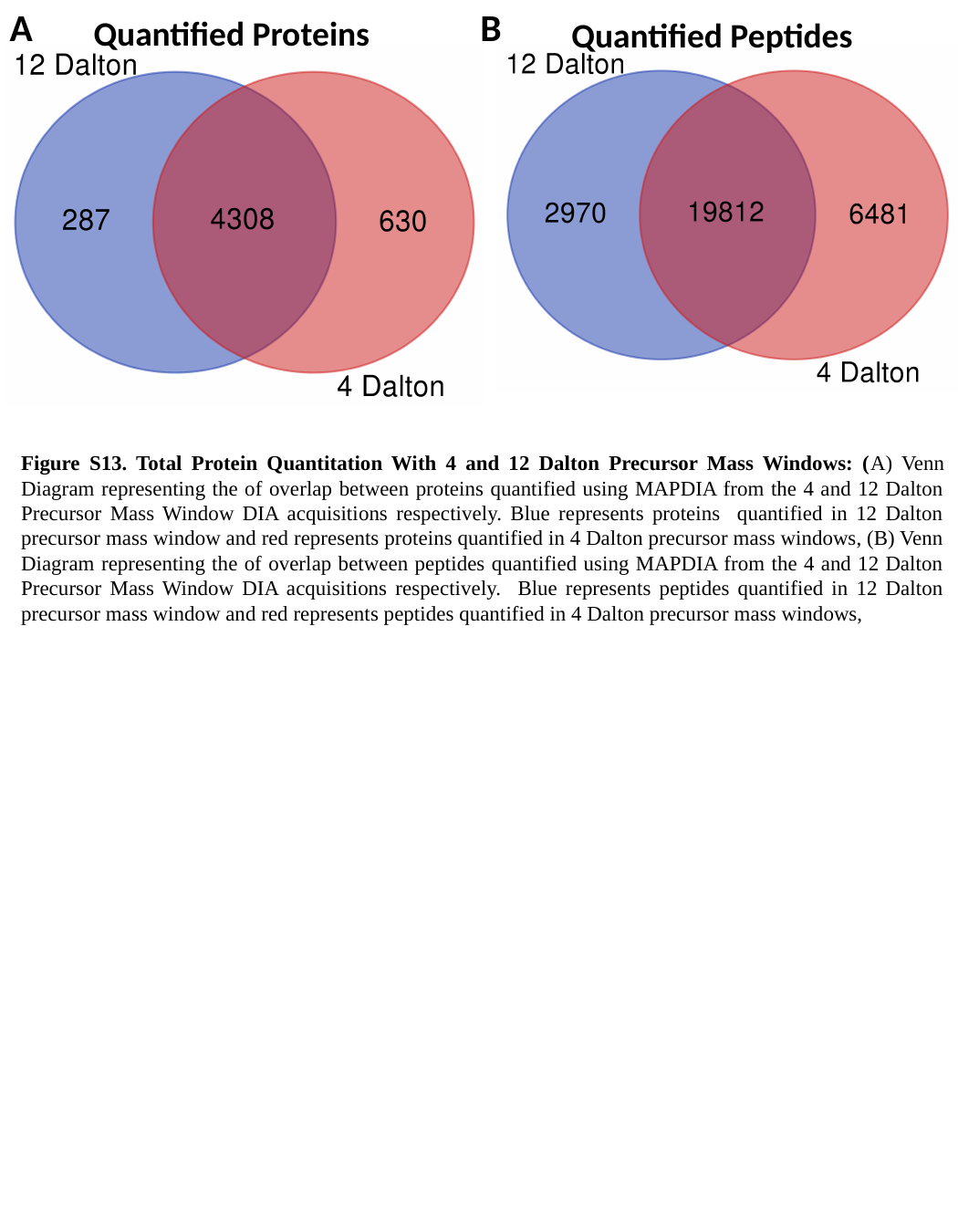

A
B
Quantified Proteins
Quantified Peptides
Figure S13. Total Protein Quantitation With 4 and 12 Dalton Precursor Mass Windows: (A) Venn Diagram representing the of overlap between proteins quantified using MAPDIA from the 4 and 12 Dalton Precursor Mass Window DIA acquisitions respectively. Blue represents proteins quantified in 12 Dalton precursor mass window and red represents proteins quantified in 4 Dalton precursor mass windows, (B) Venn Diagram representing the of overlap between peptides quantified using MAPDIA from the 4 and 12 Dalton Precursor Mass Window DIA acquisitions respectively. Blue represents peptides quantified in 12 Dalton precursor mass window and red represents peptides quantified in 4 Dalton precursor mass windows,

### Slide 22
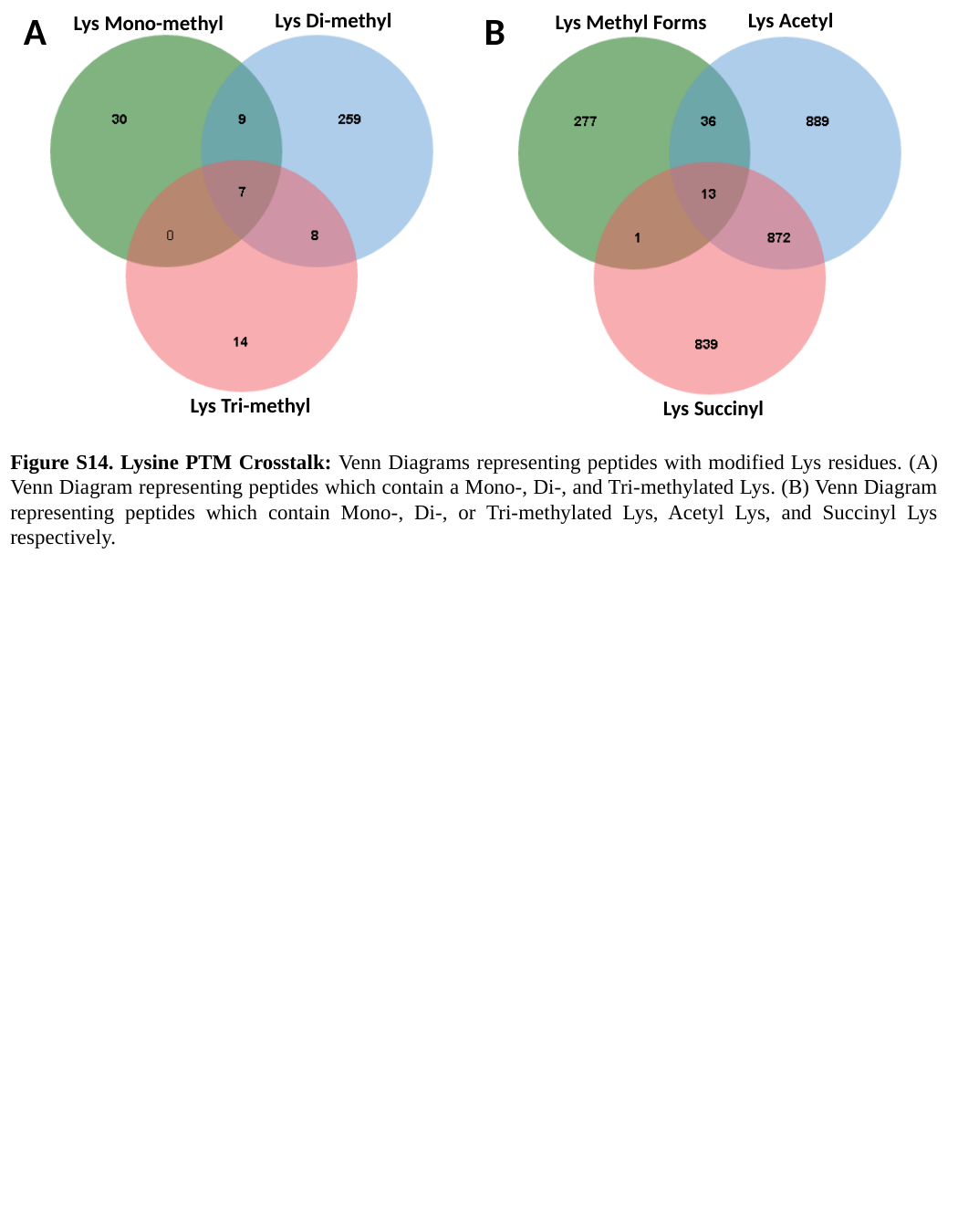

Lys Di-methyl
A
B
Lys Acetyl
Lys Methyl Forms
Lys Mono-methyl
Lys Tri-methyl
Lys Succinyl
Figure S14. Lysine PTM Crosstalk: Venn Diagrams representing peptides with modified Lys residues. (A) Venn Diagram representing peptides which contain a Mono-, Di-, and Tri-methylated Lys. (B) Venn Diagram representing peptides which contain Mono-, Di-, or Tri-methylated Lys, Acetyl Lys, and Succinyl Lys respectively.
